# Longitudinal multi-omic profiling uncovers immune escape and predictors of response in multiple myeloma

**DOI:** 10.1101/2025.05.27.656392

**Authors:** Denis Ohlstrom, William C. Pilcher, Marina E Michaud, Chaitanya R. Acharya, Sarthak Satpathy, Edgar Gonzalez-Kozlova, Reyka G. Jayasinghe, Katherine E. Ferguson, Hope Mumme, Shivani Nanda, Yizhe Song, Sowmitri Karthikeya Mantrala, Dimitra Karagkouni, Jessica Schulman, Nick Pabustan, Junia Vieira Dos Santos, Daniel Sherbenou, Jonathan Keats, Alex Gout, Steven Foltz, Alessandro Lagana, Taxiarchis Kourelis, Ravi Vij, Madhav V Dhodapkar, David Avigan, Hearn J Cho, Linda B Baughn, Ajay Nooka, Sagar Lonial, Shaji Kumar, Mehmet K Samur, Ioannis S Vlachos, Li Ding, Sacha Gnjatic, George Mulligan, Manoj Bhasin

## Abstract

Multiple myeloma (MM) is an incurable malignancy of clonally expanded plasma cells shaped by complex interactions with the immune microenvironment. To investigate immune factors driving treatment response and resistance, we conducted multi-omics profiling including CD138^neg^ single-cell RNA sequencing of 243 bone marrow samples from 102 patients (631,226 cells) and CD138^pos^ bulk RNA and whole-genome sequencing from 209 samples. Longitudinal analyses revealed that interferon gamma signaling impairs T cell memory after autologous stem cell transplant, while naïve B cell abundance and immunoglobulin diversity correlated with improved progression-free survival (HR = 0.48, p = 2.3e^-4^). At disease progression, MM cells upregulated cancer-testis antigens and immune effector genes, with concurrent B cell depletion, enrichment of myeloid-derived suppressor cell genes in monocytes, and T cell exhaustion. These findings highlight dynamic immune-tumor interactions, identifying naïve B cell reconstitution as a biomarker of durable response, and cancer-testis antigens as potential targets for high-risk disease at progression.

**Statement of Significance:** Longitudinal profiling of multiple myeloma and the immune microenvironment revealed dynamic immune-tumor interactions across the disease course. Dysfunctional CD8⁺ T cells limited memory formation post-transplant, while naïve B recovery associated with sustained treatment response. At progression, cancer-testis antigen expression associated with immunosuppression, revealing novel mechanisms of immune escape.

## Introduction

Multiple myeloma (MM) is an incurable malignancy of clonally expanded plasma cells. Disease incidence has steadily increased over the past 30 years to an estimated 36,000 new diagnoses and 13,000 deaths per year in the United States(1). Advances in MM-targeted therapies, autologous stem cell transplantation (ASCT), and immunotherapy have improved 5-year survival rates(2). However, MM remains incurable, with most patients ultimately experiencing disease progression(3,4). Several features contribute to MM’s response to therapy, including cytogenetics, intra-tumoral heterogeneity, and composition of the bone marrow immune microenvironment (IME)(5,6). In our recent analysis of baseline bone marrow samples from newly diagnosed MM patients in the MMRF Immune Atlas we identified immune alterations associated with rapid disease progression—namely reduced B cell abundance, expanded myeloid populations, and a shift toward terminally differentiated cytotoxic T cells—that remained prognostic even after adjusting for cytogenetics and patient demographics(7). However, the dynamics of the IME through treatment and interplay between plasma cells and the IME during disease progression remains incompletely understood.

Recent studies have uncovered quantitative abundance changes in the IME along with alterations in gene expression associated with the progression of MM(8,9). Key pathologic features of the IME include the development of dysfunctional T cells, accumulation of myeloid-derived suppressor cells (MDSCs), and depletion of B cells(7,8,10). Treatment with ASCT and pre-transplant conditioning with high-dose cytotoxic agents induces profound depletion and alteration of the IME, creating a period of suboptimal immune surveillance while the immune system reconstitutes. Previous studies have described patient-specific differences in the IME’s response post-ASCT and its relationship with the duration of response using flow cytometry(11), mass cytometry(10), and single-cell RNA sequencing(12). Even after reconstitution, MM cells persist and escape control by the IME through several mechanisms including direct alteration of the IME via secretion of soluble factors, indirect alteration by recruitment of immunosuppressive cell types, evasion by decreased expression of immunoreactive epitopes, and overcoming the IME by aberrant proliferative and anti-apoptotic pathways(13–16). However, the dynamics of these features through the MM disease course require additional study. Longitudinal evaluation of both MM and IME-specific features through disease and treatment stages has the potential to reveal critical insights into the mechanisms by which MM overcomes the IME.

To this end, we longitudinally evaluated MM and the IME at disease onset, response after therapy, and disease progression in patients in the Multiple Myeloma Research Foundation (MMRF) CoMMPass study (NCT01454297). To facilitate a comprehensive characterization of the disease course, we generated a single-cell profile of 243 CD138^neg^ bone marrow samples collected at disease diagnosis, response, or progression from 102 patients. Combining the analysis of this IME single-cell RNA sequencing data with bulk RNA and whole-genome sequencing data from the malignant CD138^pos^ compartment enabled deep characterization to identify IME features associated with treatment response and mechanisms by which MM escapes treatment and leads to disease progression.

## Results

### Characteristics of longitudinal multiple myeloma cohort: Clinical and single-cell landscape

To evaluate the dynamics of the IME through the MM disease course, we generated and analyzed single-cell profiles for 243 CD138^neg^ sorted bone marrow (BM) aspirates from 102 patients with longitudinal samples collected as a part of the CoMMpass Immune Atlas initiative (**Fig 1A**). The longitudinal-sample cohort was broadly representative of the clinical characteristics of the CoMMpass cohort (n=1,143). The longitudinal cohort had comparable distributions of age at diagnosis (longitudinal cohort mean = 61.6 years, CoMMpass mean = 62.9 years, p = 0.29), sex (60.4% vs 60.4% male, p = 1.0), International Staging System (ISS) stage at diagnosis (Stage I: 34.3% vs 35.1%, Stage II: 39.2% vs 35.1%, Stage III: 26.5% vs 29.8% p = 0.38, **Supplemental Fig 1A-C**). There was also comparable composition of cytogenetic groups (e.g. 1q21 gain: 35.6% vs 38.1%, p = 0.78, 17p13 deletion: 10.1% vs 10.3%, p = 1.0), and International Myeloma Working Group (IMWG)-defined risk category(17) (29.9% vs 26.3% high risk, p = 0.55, **Fig 1B, Supplemental Fig 1D**). The majority of patients in both cohorts received triplet first-line therapy, though the longitudinal cohort was much more homogenous with 85.3% receiving a proteasome inhibitor (PI), immunomodulatory drug (IMiD), and steroid (12.7% doublet, 1.0% with chemotherapy, PI, and steroid) as compared to 44.3% PI, IMiD, and steroid, 16.7% chemotherapy IMiD and steroid, 1.5% PI, chemotherapy, IMiD, and steroid in the CoMMpass cohort. There was a higher rate of ASCT as first-line therapy for the longitudinal cohort as compared to CoMMPass (67.6% vs 49.6%, p = 4.6e^-4^, **Supplemental Fig 1E-F**). Progression-free survival (p = 0.82) and overall survival (p = 0.21) were equivalent between the longitudinal and CoMMpass cohorts (**Supplemental Fig 1G-H**). Of the 102 patients with longitudinal samples, 53 had samples collected at baseline and first response, 51 had samples collected at baseline and first progression, nine of which had longitudinal samples from baseline, first response, and first progression, and 15 patients had samples taken at second/third responses/progressions (**Fig 1C**). Thus, our longitudinal cohort provides a large and diverse representative sample of MM to evaluate the IME characteristics through MM disease stages.

**Figure 1:**
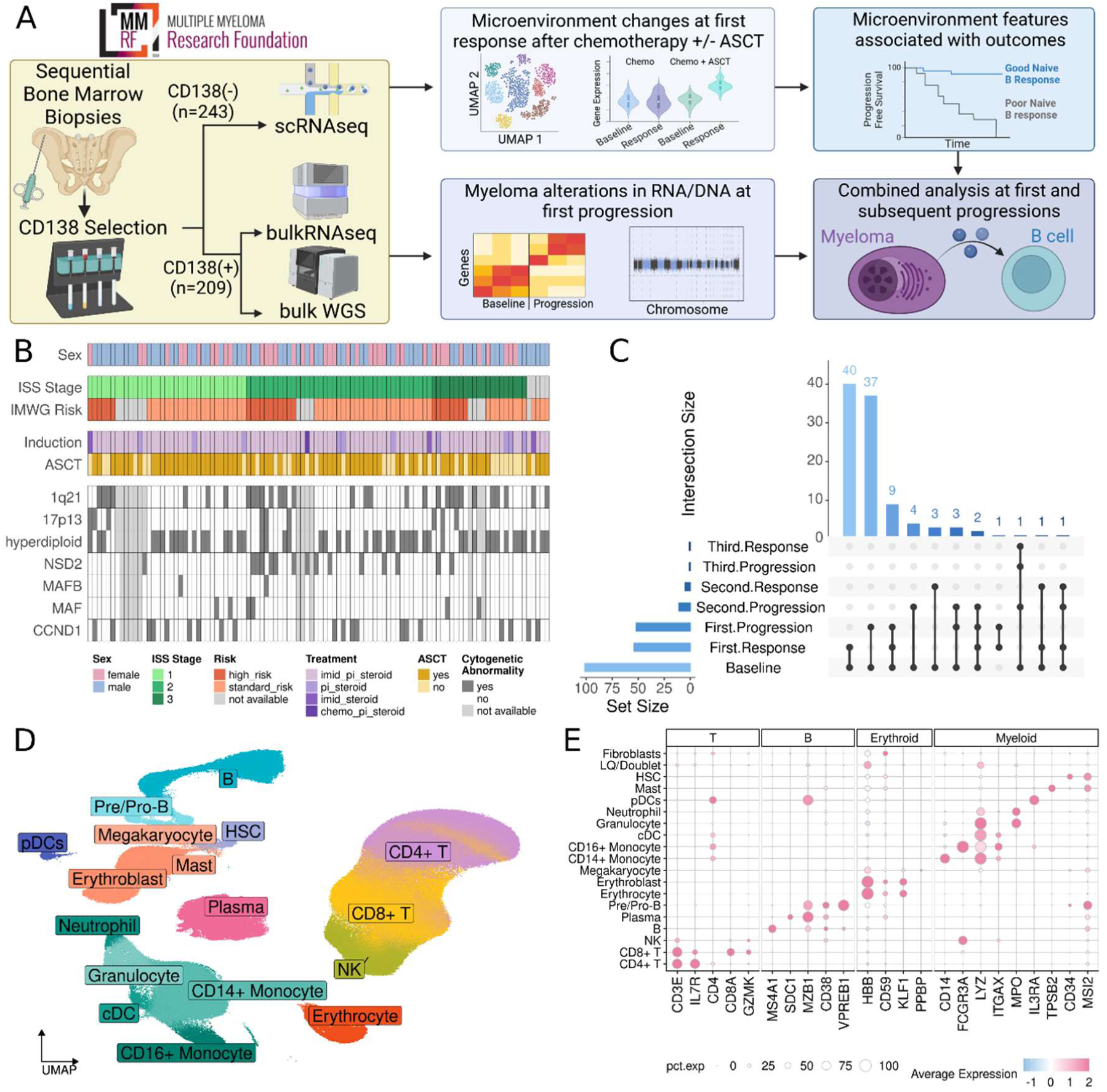
Longitudinal-sample single cell RNA sequencing of the immune microenvironment in multiple myeloma. **A.** Schematic overview of longitudinal CD138^neg^ single cell RNA sequencing (scRNAseq) and CD138^pos^ bulk RNA and whole genome sequencing analyses of multiple myeloma (MM). **B.** Tile plot of clinical and genetic features of the 102 patients with longitudinal samples in the CoMMpass scRNAseq cohort. Colors indicate reported sex (blue-male, pink-female), shades of green for international staging system (ISS) stage (1-light, 2-medium, 3-dark green), shades of orange for international myeloma working group risk category (standard-light, high-dark orange, gray for not calculable), shades of purple for induction therapy with light purple for immunomodulatory drug (IMiD), proteasome inhibitor (PI), and steroid, and progressively darker shards for PI and steroid, IMiD and steroid, and chemotherapy, PI and steroid respectively. Autologous stem cell transplant is given in shades of yellow (light yellow - no, dark yellow – yes), and mutations are dark grey for yes, white for no, and light gray for not available. **C.** Upset plot displaying the number of patients with combinations of samples across baseline and first, second, or third responses and progressions. **D.** Uniform manifold approximation and projection (UMAP) embeddings for the 631,226 cells (243 samples from 102 patients) after quality filtering and doublet removal. Colors indicate cell type. **E.** Dot plot of canonical cell type genes. The X axis has marker genes grouped by cell lineage, and the y axis shows the cell types presented in D. Dot color indicates average normalized, scaled expression with blue for low values and pink for high values. Dot size indicates the percent of cells in the corresponding cell type that expresses the corresponding gene. Schema created in BioRender.

Single-cell RNA sequencing (scRNAseq) generated profiles of 631,226 high-quality cells after quality control and doublet filtering from 243 CD138^neg^ sorted BM aspirates samples (**Fig 1D**). Cells were annotated into different cell types and states using cell type canonical marker expression information adopted from the cell annotation dictionary prepared as part of the recent Immune Atlas study(7) (**Fig 1E**). The IME of the longitudinal samples were composed of 51.8% T cells (Baseline [B] = 57.0%, response [R] = 52.1%, progression [P] = 42.5%), 7.8% NK (B = 7.4%, R = 8.3%, P = 8.1%), 12.3% B cells (B = 9.4%, R = 14.3%, P = 15.4%), 13.0% myeloid cells (B = 10.1%, R = 15.5%, P = 15.7%), 8.1% erythroid cells (B = 7.4%, R = 8.0%, P = 10.0%), and 1.4% (B = 1.4%, R = 1.4%, P = 1.3%) other small clusters (e.g. hematopoietic stem cells, mast cells, **Supplemental Fig 2A**)(7). To assess the distribution of cell types across disease stages and patients, we calculated the per-cell type entropy and found that all disease stages and patients were well represented among cell types, indicating patient and disease stage-specific cell types were not present (**Supplemental Fig 2A-D**). In addition to the expected immune populations and despite the use of the CD138^neg^ fraction, 5.6% of cells were annotated as plasma cells (B = 7.4%, R = 0.5%, P = 7.0%). We learned that samples with a greater percentage of plasma cells detected by pre-sorting flow cytometry also had a higher percentage of plasma cells in IME scRNAseq after CD138-based sorting (R = 0.76, p = 2.2e^-16^, **Supplemental Fig 2E**). Furthermore, the inferred copy number state of the plasma cells overlapped with the alterations detected in whole genome sequencing (**Supplemental Fig 3A-G**), indicating that most of the plasma cells were malignant. This suggests that samples with high myeloma loads may have experienced cell flow through the CD138 selection column due to the saturation of its capacity.

After annotating broad cell types, we further subclustered cells and annotated them based on characteristic gene expression for specific classes (e.g., naïve, memory, effector) and functional markers (e.g., NFKB, interferon response). The T and NK compartment was divided into 11 CD4^+^ subclusters, 15 CD8^+^ subclusters, and four NK subclusters (**Supplemental Fig 4A-B**). The CD4^+^ T compartment was comprised of two central memory (25.2% of CD4^+^ T cells), one naïve (44.7%), two effector (9.7%), one helper (1.0%), one regulatory (6.4%), and four other subclusters (13.0%) (**Supplemental Fig 4C**). The CD8^+^ T compartment was comprised of five effectors (36.3% of CD8^+^ T cells), one naïve (12.3%), three effector memory (19.9%), two central memory (12.7%), and four other subclusters (18.8%) (**Supplemental Fig 4D**). The four NK subclusters included one CD56^bright^ (15.6% of NK cells), one CD56^dim^ (58.7%), and two others (25.7%, **Supplemental Fig 4E**). The B compartment was divided into 8 subclusters, including one pro-B (8.2% of B cells), two pre-B (8.3%), two immature (33.5%), one naïve (33.4%), and two memory (16.4%, **Supplemental Fig 5A-B**). The myeloid compartment was divided into 14 subclusters including five CD14^+^ monocyte (61.3% of myeloid cells), one CD16^+^ monocyte (8.7%), two macrophage (4.5%), two neutrophil (5.5%), one granulocyte (8.1%), one granulocyte-monocyte progenitor (8.8%), and two cDC (3.1%, **Supplemental Fig 5C-D**) clusters. This high-resolution analysis enabled the identification of both canonical and rare immune cell populations within the bone marrow, providing a comprehensive cellular framework for downstream correlation with disease progression and response to therapy.

### Induction therapy and autologous stem cell transplantation induces compositional alterations in the B and T compartments not observed with induction therapy alone

To evaluate how first-line MM treatment impacts the IME, we compared the cellular composition and gene expression of CD138^neg^ scRNAseq in longitudinal bone marrow aspirates from baseline and first response (**Fig 2A**). Twenty-seven patients had biopsies taken after induction (median days after diagnosis = 112) and 31 patients had biopsies taken after induction and ASCT (median days after diagnosis = 272, median days after ASCT = 99, **Supplemental Fig 6A**). Comparing the clinical characteristics from the cohort of patients who had first-response samples after induction to those after induction and ASCT, the induction group was older on average (induction average age at diagnosis = 63.3 years, induction and ASCT = 55.6 years, p = 0.036), had a lower percentage of males (57.6% vs 72.7%, p = 0.045), and comparable distributions of ISS stage (p = 0.73), and IMWG risk (p = 0.80) (**Supplemental Fig 6B-E**). The majority of patients in both cohorts received PI, IMID, and steroid-based induction therapy (p = 0.8, **Supplemental Fig 6F**).

**Figure 2:**
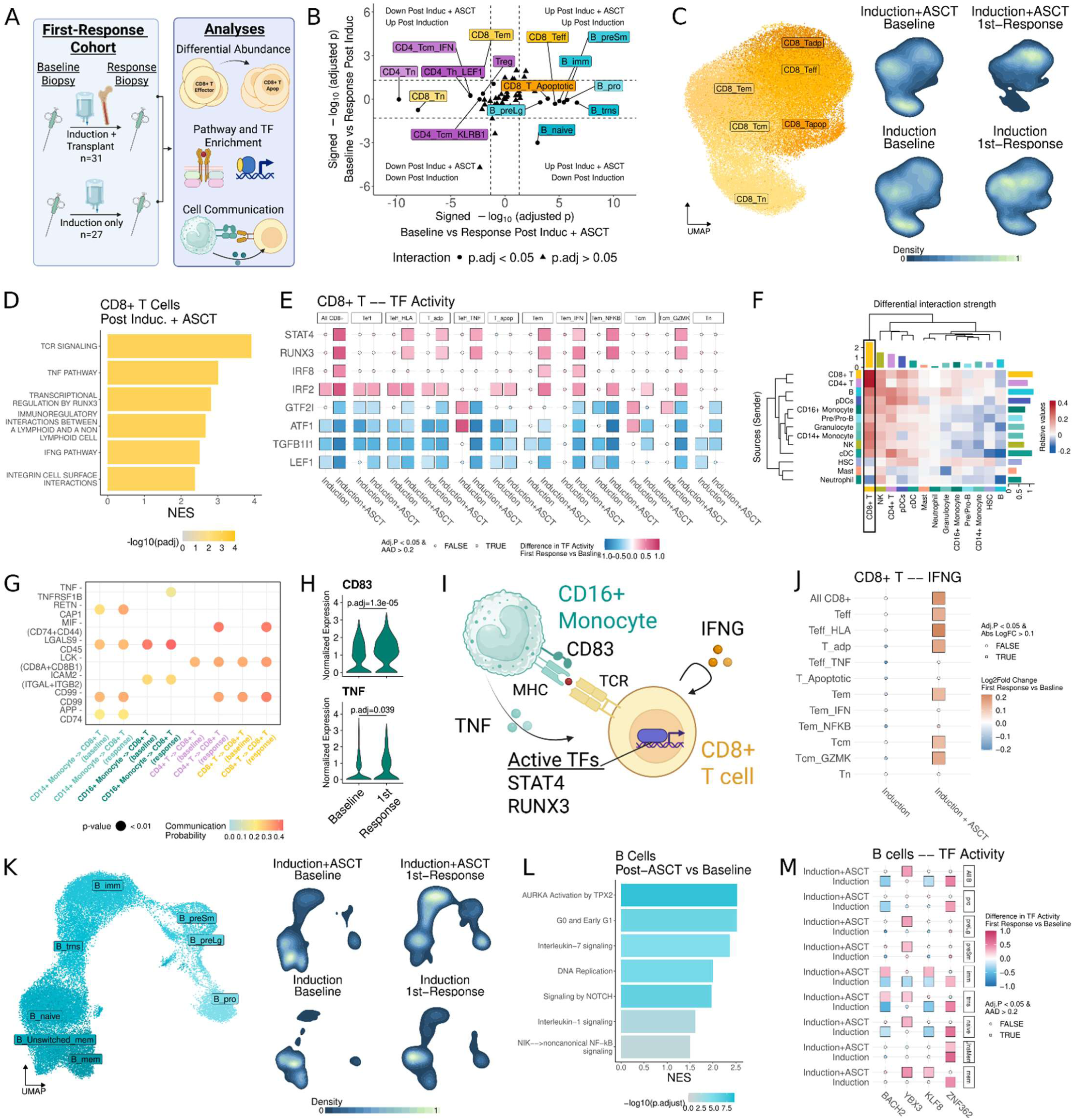
Induction therapy and autologous stem cell transplantation induces compositional alterations in the B and T compartments not observed with induction therapy alone. **A.** Schematic overview of the analyses performed. Thirty-one patients had longitudinal baseline and first response post-induction with autologous stem cell transplant (ASCT), and 27 patients had baseline and first response post-induction samples. Overview created in BioRender. **B.** Dot plot displaying the increases and decreases in immune cell subcluster proportion from baseline to first response after induction on the Y axis, and from baseline to first response post induction and autologous stem cell transplant (ASCT) on the X axis. A linear mixed effect model was used to compare subcluster proportions and changes were considered significant based on multiple comparisons adjusted p value (adj. p) < 0.05 for interaction between time and treatment (i.e. the change from baseline to first response was different in the induction and induction with ASCT groups) and an adj. p < 0.05 by Tukey post-hoc testing (i.e. the change from baseline to first response was significant in at least one of the treatment groups). Dot shape indicates the adj. p value of the interaction term with position in the plot indicating the log_2_ fold change signed adj. p for Tukey post hoc testing. The X axis depicts induction and ASCT, with values on the right indicating significant increase in proportion at first response and values on the left significant decreases. The Y axis depicts induction, with values on the top of the graph indicating significant increases at first response and values at the bottom significant decreases. **C.** Uniform manifold approximation and projection (UMAP) embeddings for CD8^+^ T cells and corresponding density plots displaying the normalized density of cells. Color in the density plots indicates density of cells with lighter colors indicating higher density. **D.** Bar plot of gene set enrichment analysis of CD8^+^ T effector cells comparing post-ASCT response to baseline. The X axis indicates normalized enrichment score (NES) and the bar color indicates -log_10_(adj. p) with grey for adj. p approaching 1.0 and yellow for small adj. p values. **E.** Heatmap of differential transcription factor (TF) activity comparing first response to baseline calculated using Decoupler. Decoupler uses a univariate linear model to calculate activity of a TF based on the expression of known downstream genes, with positive values indicating relatively high activity and negative values relatively decreased activity. Heatmap colors indicate TF activity with pink for increased activity and blue for decreased activity. Dot shape indicates significance with boxes for TFs with adjusted p < 0.05 and an absolute average activity difference > 0.2, and circles for non-significant changes. **F.** Heatmap depicting the difference in interaction strength between immune cells at first-response post-ASCT versus baseline as calculated by CellChat. Heatmap colors indicate the relative strength of interaction between two cell types at first-response post-ASCT as compared to baseline with red for increased and blue for decreased. Bars along the top of the plot indicate cumulative relative strength of incoming signals, and bars along the right side indicate cumulative relative strength of outgoing signals. **G.** Dot plot of pathways with differential communication probability between first response post-ASCT and baseline. The X axis indicates the interacting cell types and timepoint, the Y axis indicates the genes predicted to interact. Dots are colored based on communication probability with blue for low probability and red for high probability. **H.** Violin plots of gene expression in CD16^+^ monocytes. Differential expression was executed using limma voom with Benjamini-Hochberg multiple comparisons correction. The Y axis displays normalized counts. **I.** Schematic synopsis of potential interactions between CD16^+^ monocytes and CD8^+^ T cells. **J.** Heatmap displaying the log_2_ fold change of *IFNG* expression comparing first response to baseline in CD8^+^ T cells. Color indicates log_2_ fold change with orange values for positive values and blue for negative values. Shape indicates statistical significance with boxes for significant changes and circles for non-significant. **K.** UMAP embeddings for B cells and corresponding density plots displaying the normalized density of cells as in C. **L.** Bar plot of gene set enrichment analysis of B cells comparing post-ASCT response to baseline. The X axis indicates NES and the bar color indicates -log_10_(adj. p) with grey for adj. p approaching 1.0 and blue for small adj. p values. **M.** Heatmap of transcription factor activity in B cells calculated using Decoupler as in Fig E.

Given that ASCT is associated with profound disruption of the IME(10), we hypothesized that compositional changes would be observed to a greater extent after induction and ASCT as compared to after induction alone. Consistent with previous reports(11,18), at the broad cell type level, cellular proportion analysis depicted a significant increase after ASCT in B cells (log_2_-fold proportion, L_2_FP = 0.93, p. adj = 1.3e^-4^) and pre/pro B cells (L_2_FP = 1.75, adj. p = 2.1e^-7^) as well as depletion of CD4^+^ T cells (L_2_FP = -1.2, adj. p = 3.8e^-10^, **Supplemental Figure 7A-F**). Strikingly, cellular proportion analysis at the subcluster level identified 15 subclusters from the B and T compartments that uniquely changed from baseline to first response after ASCT, whereas only two subclusters had significant changes from baseline to first response that were unique to induction only (**Fig 2B, Supplemental Figure 8A-E**). These findings indicate that ASCT, but not induction therapy alone, drives substantial remodeling of the immune microenvironment. To further characterize the pathways and interactions driving changes in the T cell compartment, we next performed a focused analysis of its composition and gene expression.

### Interferon and tumor necrosis factor signaling support CD8^+^ T effector bias towards apoptosis over effector memory formation in the post-ASCT IME

Comparison of the proportion of T subclusters from baseline to first response after ASCT revealed a markedly inflammatory IME not observed after induction alone. As expected, naïve CD4^+^ (L_2_FP = -1.84, adj. p = 1.7e^-10^), and CD8^+^ (L_2_FP = -1.93, adj. p = 9.9e^-9^) T cells were significantly depleted, likely reflecting delayed reconstitution due to age-related thymic involution(19,20). Consistent with an inflammatory IME, cytotoxic CD8^+^ T effector cells were significantly expanded post-ASCT (L_2_FP = 1.35, adj. p = 2.4e^-5^) while immunosuppressive T regulatory cells were significantly reduced (L_2_FP = -0.7, adj. p = 9.5e^-3^, **Supplemental Fig 8D-E**). Notably, CD8^+^ T effector appeared to be biased towards apoptosis over effector memory formation, with early apoptotic CD8^+^ effector T cells depicting significant increase post-ASCT (L_2_FP = 1.57, adj. p = 1.1e^-4^), while CD8+ effector memory T cells were significantly decreased (L_2_FP = -0.94, adj. p = 8.0e^-3^), **Fig 2C, Supplemental Fig 8D-E**). The skew of CD8^+^ T cells toward apoptosis rather than effector memory formation may limit the durability of anti-tumor T cell responses post-ASCT, ultimately constraining long-term immune surveillance. To investigate the mechanisms driving the bias away from effector memory formation, we next characterized signaling pathways within and between immune populations that influence CD8⁺ effector T cell fate in the post-ASCT IME.

To investigate the transcriptional programs underlying the expansion of effector CD8⁺ T cells and their bias toward apoptosis over effector memory formation, we compared the gene expression profiles of CD8⁺ T cells following ASCT versus induction alone. Differential expression and gene-set enrichment analysis (GSEA) identified several effector-promoting pathways including increased tumor necrosis factor signaling (NES = 3.0, adj.p = 8.6e^-4^) and interferon-gamma signaling (NES = 2.51, adj.p = 1.5e^-3^, **Fig 2D**) that were not found after induction only (**Supplemental Fig 9A**). Notably, interferon-gamma signaling has been shown to drive CD8^+^ effector T cells towards apoptosis(21,22), aligning with our observation of increased early apoptotic effectors post-ASCT. Transcription factor analysis further identified enrichment for several transcription factors downstream of interferon-gamma stimulation including STAT4 (average activity difference [AAD] = 0.70, adj.p = 1.0e^-250^) and RUNX3 (AAD = 0.73, adj.p = 1.0e^-250^, **Fig 2E**).

Given that GSEA identified enrichment in CD8^+^ T cells for interaction with a non-lymphoid cell (NES = 2.68, adj.p = 8.6e^-4^), we next performed differential cell communication analysis(23) to identify potential intracellular signals influencing T cell fate. This analysis revealed that CD8⁺ effector T cells had the highest incoming signal strength post-ASCT (**Fig 2F**), in contrast to only moderate incoming signaling following induction alone (**Supplemental Fig 9B**).Examining the differentially active pathways between baseline and post-ASCT response, we found TNF signaling between CD16^+^ monocytes and CD8^+^ T cells was significant only in the post-ASCT samples (p < 0.01, **Fig 2G, Supplemental Fig 9C**), suggesting a monocytic contribution to the inflammatory state of CD8+ T cells. Given TNF’s dual role in promoting effector function and driving apoptosis during sustained activation(24), this signaling axis likely contributes to the skewing of CD8⁺ T cells toward apoptosis over long-lived memory formation.

Together, the GSEA and cell communication analyses highlight a multifaceted mechanism shaping the CD8⁺ T cell compartment post-ASCT. Compared to baseline samples, CD16^+^ monocytes exhibit increased TNF (L_2_FC = 0.18, adj. p = 0.039) corresponding with the enriched TNF signaling observed in CD8⁺ T cells and supporting a shift towards early-apoptosis (**Fig 2D, H-I**). Corresponding with the enriched T cell receptor signaling observed in CD8^+^ T cells, CD16^+^ monocytes also have increased expression of *CD83* (L_2_FC = 0.38, adj. p = 1.3e^-5^) which supports MHC stability(25). In parallel, CD8^+^ effector T cells displayed increased expression of *IFNG* (L_2_FC = 0.20, adj. p = 1.2e^-70^), consistent with an autocrine or paracrine loop sustaining *IFNG* driven inflammatory signaling and the observed pathway and transcription factor enrichment (**Fig 2I-J**). Collectively, these findings suggest that reciprocal interactions between activated myeloid and T cell populations in the post-ASCT bone marrow foster a hyperinflammatory environment that promotes expansion of effector CD8⁺ T cells while biasing them toward apoptosis rather than durable memory formation. Targeting key inflammatory mediators such as *IFNG* and *TNF* may offer therapeutic opportunities to restore immune balance and enhance long-term immune surveillance following ASCT.

### Post-ASCT B cell reconstitution is supported by pro-proliferative gene expression

In addition to the changes observed in the T cell compartment, six B cell subclusters were significantly expanded following ASCT, but not after induction therapy alone (**Fig 2B**). ASCT significantly enhanced pro-B (L_2_FP = 1.5, adj. p = 1.8e^-6^), large pre-B (L_2_FP = 1.47, adj. p = 4.5e^-4^), small pre-B (L_2_FP = 2.2, adj. p = 3.7e^-6^), immature B (L_2_FP = 9.7e^-6^), transitional B (L_2_FP = 2.11, p = 2.4e^-7^), and naïve B (L_2_FP = 1.1, adj. p = 9.9e^-4^) populations. These changes were not observed in the induction-only cohort (**Fig 2B, Supplemental Fig 7**), suggesting that ASCT—rather than induction alone—drives active B cell repopulation.

To investigate the underlying mechanisms of this expansion, we examined the transcriptional profiles of B cell subpopulations post-ASCT (**Fig 2K**). Differential gene expression and GSEA revealed enrichment of pro-proliferative pathways, including aurora kinase A (AURKA) activation by TPX2 (NES = 2.53, adj. p = 6.1e^-9^) and DNA replication (NES = 2.0, adj. p = 7.6e^-6^), as well as pro-activation/differentiation pathways such as NOTCH signaling (NES = 2.0, adj. p = 7.6e^-6^) and non-canonical NFKB signaling (NES = 1.5, adj. p = 0.026, **Fig 2L**). These transcriptomic programs suggest coordinated proliferation and maturation of the B cell compartment post-ASCT. Supporting the GSEA findings, transcription factor analysis found that post-ASCT B-cells sustained levels BACH2 activity, a known driver of B cell development(26), while post-induction B cells had decreased BACH2 activity (AAD = -0.38, adj.p = 2.6e^-34^, **Fig 2M**). Moreover, immature (AAD = 0.29, adj. p = 3.2e^-50^) and memory (AAD = 0.26, adj. p = 2.7e^-5^) B cells were enriched for KLF8 activity, a transcription factor associated with cell cycle progression(27). These transcriptional shifts underscore the presence of a proliferative B compartment following ASCT that may help drive effective B cell reconstitution.

### Patients with longer than median progression-free survival after autologous stem cell transplant have a more robust B cell recovery

Next, we aimed to evaluate the association between post-ASCT changes in the IME and the duration of clinical response to determine whether specific immune subpopulations induced by ASCT are linked to improved outcomes. To achieve this goal, we stratified the 31 patients with baseline and post-ASCT first-response samples based on greater than or less than the median progression-free survival (PFS) (1,491 days). Samples were annotated as either greater than the median PFS (GMpfs, n = 15) or less than the median PFS (LMpfs, n = 9) with seven censored before the median PFS (**Fig 3A, Supplemental Fig 10A-D**). Comparative analysis of the proportion of broad cell types between PFS groups found no differences in change from baseline that are unique to either group (**Supplemental Fig 11A-D**). However, subcluster analysis depicted that patient with GMpfs had a nearly 4-fold increase in naïve B cells (L_2_FP = 1.96, adj. p = 8.6e^-4^) that was not observed in the LMpfs group (L_2_FP = 0.19, adj. p = 0.79, **Fig 3B-C**). The enrichment of naïve B cells in GMpfs patients may reflect a more robust capacity for developing antigen-specific immune responses post-ASCT, potentially contributing to sustained disease control—an immunologic feature notably lacking in patients with shorter PFS.

**Figure 3:**
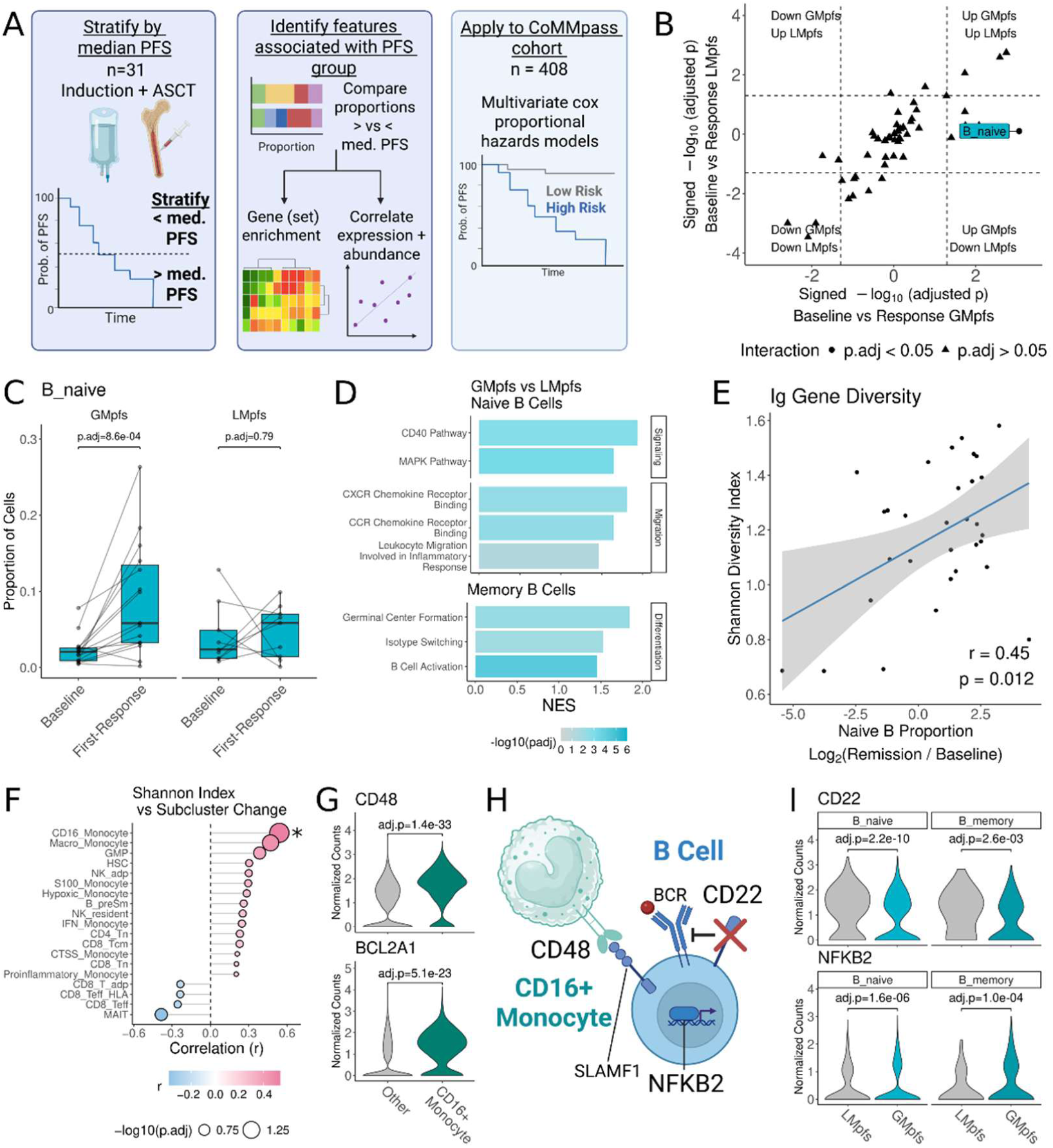
Patients with longer than median progression free survival after autologous stem cell transplant have a more robust B cell recovery. **A.** Schematic representation of analyses performed. Thirty-one patients with longitudinal baseline and first response post autologous stem cell transplant (ASCT) samples were split into greater than median progression free survival (GMpfs, n=15), less than median PFS (LMpfs, n = 9), with seven patients censored prior to median PFS. Cell type proportions and expression were compared between GMpfs and LMpfs to examine what changes in the IME associate with clinical outcomes. **B.** Dot plot displaying the increases and decreases in immune cell subcluster proportion from baseline to first response after ASCT in the LMpfs group on the Y axis, and from baseline to first response post ASCT for the GMpfs group on the X axis. A linear mixed effect model was used to compare cell subcluster proportions and changes were considered significant based on multiple comparisons adjusted p value (adj. p) < 0.05 for interaction between time and PFS group (i.e. the change from baseline to first response was different in the GMpfs and LMpfs groups) and an adj. p < 0.05 by Tukey post-hoc testing (i.e. the change from baseline to first response was significant in at least one of the PFS groups). Dot shape indicates the adj. p value of the interaction term with position in the plot indicating the log_2_ fold change signed adj. p for Tukey post hoc testing. The X axis depicts GMpfs, with values on the right indicating significant increase in proportion at first response and values on the left significant decreases. The Y axis depicts LMpfs, with values on the top of the graph indicating significant increases at first response and values at the bottom significant decreases. **C.** Box and dot plots of cell proportions for naïve B cells. Lines connect the dots corresponding to the same patient at each timepoint. P values were calculated using Tukey post-hoc testing as described in A. **D.** Bar plot of gene set enrichment analysis of naïve B cells (top) and memory B cells (bottom) comparing GMpfs to LMpfs. The X axis indicates normalized enrichment score (NES) and the bar color indicates -log_10_(adj. p) with grey for adj. p approaching 1.0 and blue for small adj. p values. **E.** Dot plot displaying the relationship between naïve B abundance and diversity of immunoglobulin gene expression in post-bone marrow B cells. Based on the enrichment for migration in naïve B cells and germinal center formation/isotype switching in memory B cells, we evaluated the diversity of immunoglobulin gene expression in these subclusters (i.e. the B subclusters that have typically left the bone marrow). The X axis displayed the log_2_ fold proportion of naïve B cells comparing first response post ASCT to baseline. The Y axis displays the Shannon diversity index on immunoglobulin gene expression in post-bone marrow B cells. For each patient, the expression of genes for each immunoglobulin isotype (IgD, IgM, IgG, IgE, IgA) were averaged and normalized. Then the five normalized isotype expression values were used to calculate the Shannon diversity index. R value and significance were calculated using Pearson correlation. **F.** Lollipop plot for correlation of other subclusters abundance with immunoglobulin diversity. To identify other subclusters that associated with high immunoglobulin diversity we correlated each other subcluster’s log_2_ fold proportion with the same Shannon index values calculated in E. Lollipop colors indicate the r value with pink for positive values and blue for negative values. Lollipop size indicates -log_10_ multiple comparisons adjusted p value, with an asterisk indicating adjusted p < 0.05. **G.** Violin plots of normalized counts of *CD48* (top) and *BCL2A1* (bottom) in CD16^+^ monocytes and all other monocytes in first response post-ASCT samples. Benjamini-Hochberg adjusted P values were calculated in limma. **H.** Schematic representation of possible interaction between CD16^+^ monocytes and B cells. **I.** Violin plots of normalized counts of *CD22* (top) and *NFKB2* (bottom) in naïve and memory B cells comparing LMpfs to GMpfs patients in first response post-ASCT samples. Multiple comparisons adjusted P values were calculated using limma.

To characterize the transcriptomic changes associated with the high naïve B proportion in GMpfs patients, we compared the expression of naïve B cells at first response between the GMpfs and LMpfs groups. Naïve B cells in GMpfs to have enrichment of activation/survival signals (CD40 pathway: NES = 1.92, adj. p = 1.7e^-3^, MAPK pathway: NES = 1.64, adj. p = 4.8e^-4^) and migratory pathways (CCR chemokine receptor binding: NES = 1.65, adj. p = 3.5e^-3^, CXCR receptor binding: NES = 1.81, adj. p = 3.0e^-3^, **Fig 3D**). Based on these findings, we posit that the relative abundance, activation, and migration of naïve B cells in GMpfs patients would promote activity in developmentally downstream B subtypes as well. Comparing the memory B cells between GMpfs and LMpfs patients we also observed enrichment for germinal center formation (NES = 1.86, adj. p = 2.5e^-3^), isotype switching (NES = 1.52, adj. p = 6.7e^-3^), and activation (NES = 1.45, adj. p = 3.2e^-5^) pathways, suggesting that abundant naïve B cells supported production of a diverse immunoglobulin (Ig) repertoire. To assess the potential relationship of abundant naive B cells supporting diverse Ig expression, we calculated the Shannon diversity index on Ig gene expression in the B cells that undergo isotype switching (i.e., naïve to memory cells). The analysis showed that proportion of naïve B cells is significantly positively associated with Ig gene Shannon diversity index (R = 0.45, p = 0.012, **Fig 3E**). Thus, increased naïve B cell proportion, transcriptional activation, and migratory capacity collectively contribute to a more diverse and durable immune repertoire in patients with extended post-ASCT remission.

Finally, to determine which other cell types/subtypes in the IME may contribute to diverse Ig production, we correlated the post-ASCT change (L_2_FP) of each non-B immune cell type with Ig Shannon index. Interestingly, CD16^+^ monocytes were the only cell type with a significant correlation with Ig diversity (R = 0.54, adj. p = 0.032) and have been previously reported to associate with B cell differentiation and Ig production(28), suggesting a possible interaction with B cells that would contribute to diverse Ig production (**Fig 3F**). Compared to the other clusters of monocytes which were not correlated with Ig diversity, CD16^+^ monocytes had increased levels of *CD48* (L_2_FC = 0.76, adj. p = 1.4e^-33^), a cell surface receptor that binds *SLAMF1*/CD244 to promote B cell activation(29), as well as *BCL2A1* (L_2_FC = 0.72, adj. p = 5.1^-23^) which promotes monocyte survival in the IME(30) (**Fig 3G-H**). Consistent with the increased activation of B cells from the GMpfs group observed in GSEA, both naïve and memory B cells had decreased expression of the BCR antagonist, *CD22*(31), (naïve: L_2_FC = -0.41, adj. p = 2.2e^-10^, memory: L_2_FC = -0.42, adj. p = 2.6e^-3^), and increase in downstream /activation genes *NFKB2* (naïve: L_2_FC = 0.25, adj. p = 1.6e^-6^, memory: L_2_FC = 0.47, adj. p = 1.0e^-4^, **Fig 3I**)(32). These results indicate a potential supportive role for CD16⁺ monocytes in fostering B cell activation and sustaining Ig repertoire diversity following ASCT. Through the expression of costimulatory molecules, and in concert with heightened BCR signaling observed in GMpfs patients, this subset may represent critical cell types/subtypes linking innate immune remodeling to durable humoral immunity.

### Low post-ASCT serum immunoglobulin diversity is associated with early disease progression

Given that a high proportion of naïve B cells post-ASCT was associated with greater than median PFS and increased diversity of Ig gene expression, we next aimed to determine if post-ASCT serum Ig measurements were also associated with patient outcomes. Using the CoMMpass single-cell Immune Atlas dataset, we randomly split the 408 patients who underwent ASCT, while ensuring equal distribution of ISS stage, into derivation (n = 205) and validation (n = 203) cohorts (**Fig 4A**). The resulting derivation and validation cohorts had comparable distributions of clinical features, including age, BMI, IMWG risk, sex, and induction therapy (all p > 0.05, **Fig 4A**). The optimal cut point calculated from the derivation cohort using the CutP method(33) (0.768) was applied to both the derivation and validation cohorts, stratifying 20.5% of patients as Shannon low and 79.5% of patients as Shannon high, consistent with 75.0% of patients increasing naïve B cells post-ASCT (**Fig 3C**). The low Shannon index samples were associated with shorter PFS in both the derivation (hazard ratio, HR = 2.3, p = 4.9e^-5^) and validation cohorts (HR = 2.1, p = 2.3e^-4^, **Fig 4B-C**). Overall survival (OS) was also poorer for the low Shannon group in the derivation cohort (HR = 2.2, p = 5.2e^-3^) and did not reach significance in the validation cohort (HR = 1.5, p = 0.13, **Supplemental Fig 12A-B**). The worse outcomes observed in low Shannon index patients suggest that inadequate B cell reconstitution post-ASCT may reflect poor immune function and limited ability of the IME to maintain anti-tumor immune surveillance. To further characterize whether differences in outcomes were seen by the Shannon group while accounting for the ISS stage, we compared high and low Shannon within ISS stage 1, 2, and 3 patients. In the derivation cohort, a low Shannon index was associated with worse outcomes in all ISS stages (ISS 1: HR = 2.0, p = 3.3e^-2^, ISS 2: HR = 2.3, p = 0.028, ISS 3: HR = 3.3, p = 2.5e^-3^), though in the validation cohort differences were detected in ISS stage 1 and 2 (ISS 1: HR = 2.0, p = 0.039, ISS 2: HR = 2.8, p = 1.9e^-3^, ISS 3: HR = 1.6, p = 0.26, **Fig 4D-E, Supplemental Fig 12C-D**). Consistent with our observation that naïve B proliferation post-ASCT is associated with good outcomes and diverse Ig gene expression, patients with higher Shannon index on serum Igs had longer PFS in derivation and validation cohorts from the CoMMpass single-cell dataset. These findings support the use of serum Ig diversity as a surrogate for humoral reconstitution and a potential prognostic biomarker for risk stratification following ASCT in MM.

**Figure 4:**
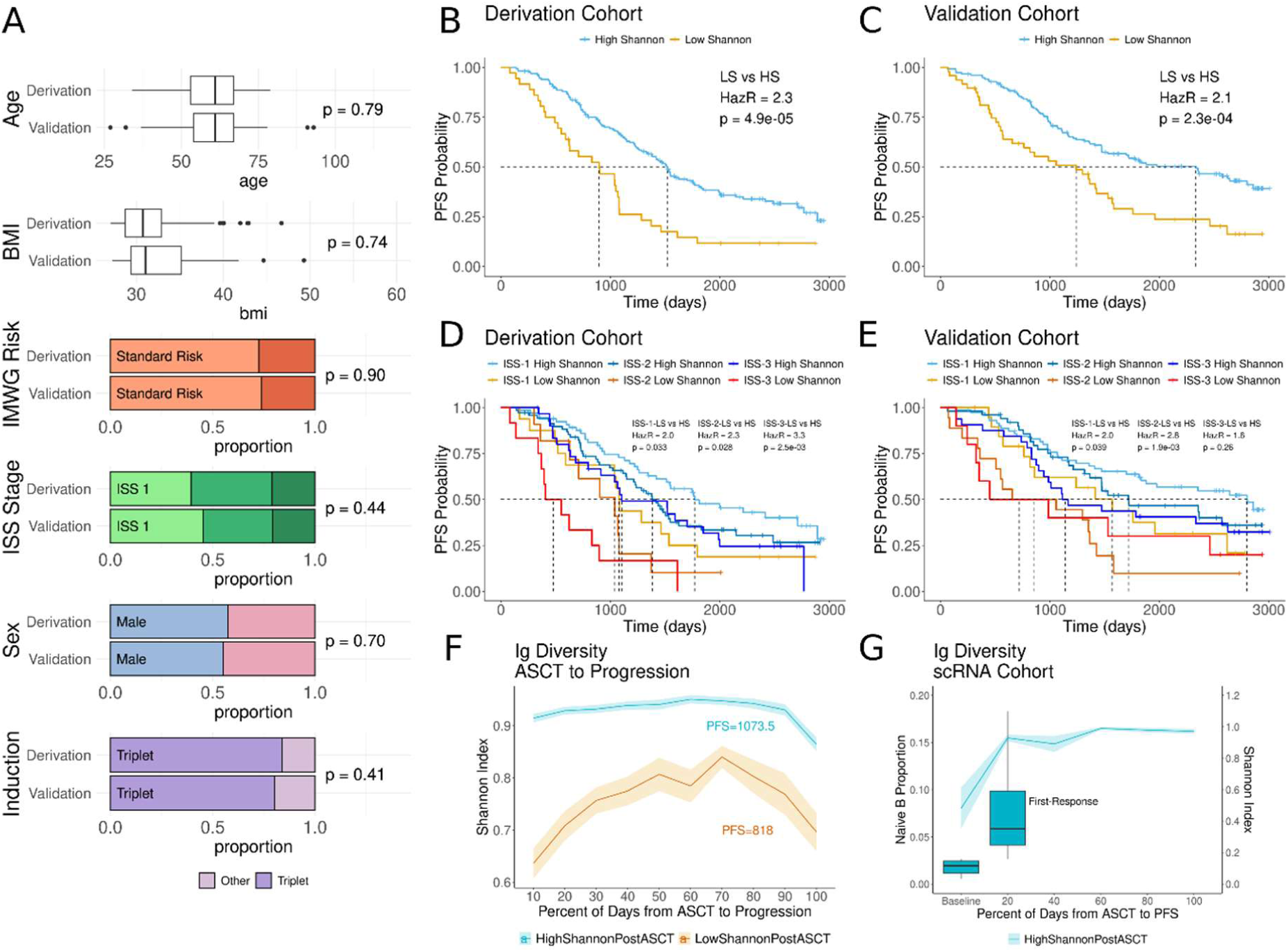
Low post-ASCT serum immunoglobulin diversity associated with early progression. **A.** Box plots and bar plots displaying the clinical characteristics of the derivation and validation cohorts. Four hundred and eight patients in the CoMMpass dataset who underwent autologous stem cell transplant were randomly split while ensuring equal distribution of International Staging System (ISS) disease stages into derivation (n=205) and validation (n=203) cohorts. For age and body mass index (BMI) p values were computed using the student’s t-test. For International Myeloma Working Group (IMWG) risk, ISS stage, sex, and induction therapy p values were computed using the chi-squared test. IMWG risk is displayed in shades of orange (light-standard risk, dark-high risk), ISS stage in shades of green (light-1, medium-2, dark-3), blue for males and pink for females, and shades of purple for induction therapy. **B-C.** Kaplan Meier curve for the derivation (B) and validation (C) cohorts of progression-free survival (PFS). Lines are colored by group with yellow for low Shannon index on serum immunoglobulins and blue for high Shannon index. Patients were grouped into high and low Shannon index groups using the CutP method on the derivation cohort. Hazard ratio (HazR) and p value calculated by Cox proportional hazards model. **D-E.** Multivariate Kaplan Meier curves for PFS of ISS stage and Shannon index group on serum immunoglobulins. Line colors indicate the ISS-Shannon group with shades of blue for high Shannon and shades of yellow-orange for low Shannon index. **F.** Line plot showing the Shannon Index on serum immunoglobulins from ASCT to progression. For the 215 patients who experienced progression on the study, the day of ASCT was treated as day 0 and each visit between ASCT and disease progression was partitioned into buckets of 10% (e.g. 0-10%, 10-20% of the days from ASCT to progression) to visualize patients with different PFS on a comparable scale. The line indicates the average Shannon index and the highlighted color indicates the standard error on the mean. Lines are blue for the high Shannon index and yellow for the low Shannon index. **G.** Line and box plot of Shannon index in clinical serum measurements and naïve B proportion in CD138^neg^ single-cell RNA sequencing. Eleven patients with naïve B increase above the lower tertile after ASCT had a progression on the study and were used for this analysis. As with F, days to progression were binned (by 20%) and the X axis displays percent of days to progression to compare patients with different durations of response on a comparable scale. The left Y axis and boxplot displays the proportion of naïve B cells at baseline and first response. The line and right Y axis display the average Shannon index.

To determine how Ig diversity changed over time in patients with high/low Shannon index post-ASCT, we evaluated the 215 patients who were not censored (i.e., progressed in the study). Treating the day of ASCT as day 0, we observed that patients with high post-ASCT Shannon index on serum Igs sustained high Shannon values until progression (**Fig 4F**). While patients with low post-ASCT Shannon index increased their Ig diversity over time, it did not reach comparable values to the patients with high post-ASCT Shannon (**Fig 4F**). To assess if naïve B proportion in scRNAseq was associated with sustained serum Ig Shannon index values we evaluated the 11 uncensored patients with high naïve B proportion (above the lower tertile, 0.063, **Fig 3C**) at first-response post-ASCT. All 11 patients were in the high serum Shannon index post-ASCT group, and consistent with the larger cohort, depicted sustained Ig diversity until progression (**Fig 4G**). Together, these analyses suggest that low Ig diversity post ASCT is a poor prognostic biomarker and is associated with persistently restricted diversity in the Ig repertoire. Impaired B cell reconstitution may reflect a broader failure of immune recovery and provide potential value for disease monitoring after ASCT.

### Cancer-testis antigen expression MM associates with B cell depletion at first progression

To understand how myeloma escapes the IME at first progression, we compared the gene expression of CD138^pos^ bulk RNA sequencing and whole genome sequencing from 67 patients with matched baseline and first-progression biopsies, searching for genes strongly associated with this relapse event (**Fig 5A**). Interestingly, differential expression of myeloma cells from first progression versus baseline depicted significantly increased expression of several cancer-testis antigens (CTAg), genes that are typically epigenetically silenced outside of the developing germ cells and placenta but exhibit pleiotropic functions in human cancers and are associated with elevated risk in MM(34–36) (**Fig 5B**). Paradoxically, CTAgs are frequently expressed in myeloma cells and can elicit a natural anti-tumor immune response, suggesting that these cells leverage additional mechanisms to escape IME control(36). Examining the expression of CTAgs, we observed significant sample-wise heterogeneity with a subset of samples (n = 27) driving the differential expression of CTAg genes (e.g. CTAG2: 35.8% of samples, SSX1: 32.0%, PAGE2B: 8.2%, MAGEA3: 32.8%, **Supplemental Fig 13A-B**). These twenty-seven of the 134 samples were labeled CTAg-enriched; four came from baseline samples, out of which three had persistent enrichment to progression, and 20 samples were not enriched at baseline but became enriched at first progression (**Supplemental Fig S14 A**). This pattern of a gene expression profile that is rare at baseline with increased prevalence at progression was also observed for a group of MM patients with high expression of proliferation-related genes in a recent CoMMpass study project by Skerget *et al.*(37). The proliferation-related genes enrichment analysis demonstrated that CTAg enriched samples expressed significantly higher proliferation-associated genes and pathways (p = 1.9e^-8^, **Fig 5C, Supplemental Fig S14B**). Also consistent with Skerget *et al.*, we found that patients with CTAg enrichment at first progression had considerably shorter time to second progression (HR = 2.9, p = 1.5e^-3^) and overall survival (HR = 4.5, p = 5.8e^-6^, **Supplemental Fig 15A-D**). In addition to expressing higher levels of CTAgs, this subgroup of samples also expressed more unique CTAg genes, with a median of 21 CTAgs >1 TPM as compared to two CTAgs >1 TPM from all other samples suggesting epigenetic dysregulation of CTAg in MM reversing normal silencing to enhance pro-tumoral properties (p = 4.9e^-5^, **Fig 5C**). Supporting this hypothesis, we found that CTAg enrichment was not associated with common MM driving copy number alterations, single nucleotide variants, structural variants, or fusion transcripts (**Supplemental Fig 14C-D**), but rather had preferential alteration in genes that code for chromatin modifying proteins (**Supplemental Fig 16A-B, Supplemental Fig 14E-G**). Finally, CTAg-enriched samples highly expressed several genes encoding cytokines and chemokines, including IL-32, FAM3B, and CXCL8 (p = 5.7e^-10^, **Fig 5C**). Thus, it appears that CTAg-enriched MM leverages both proliferative signaling and inflammatory cytokine expression to subvert the immunologic constraints of the IME at disease progression. These findings suggest a distinct transcriptional program characterized by CTAg enrichment, proliferation, and immune modulation that may enable immune evasion and disease acceleration.

**Figure 5:**
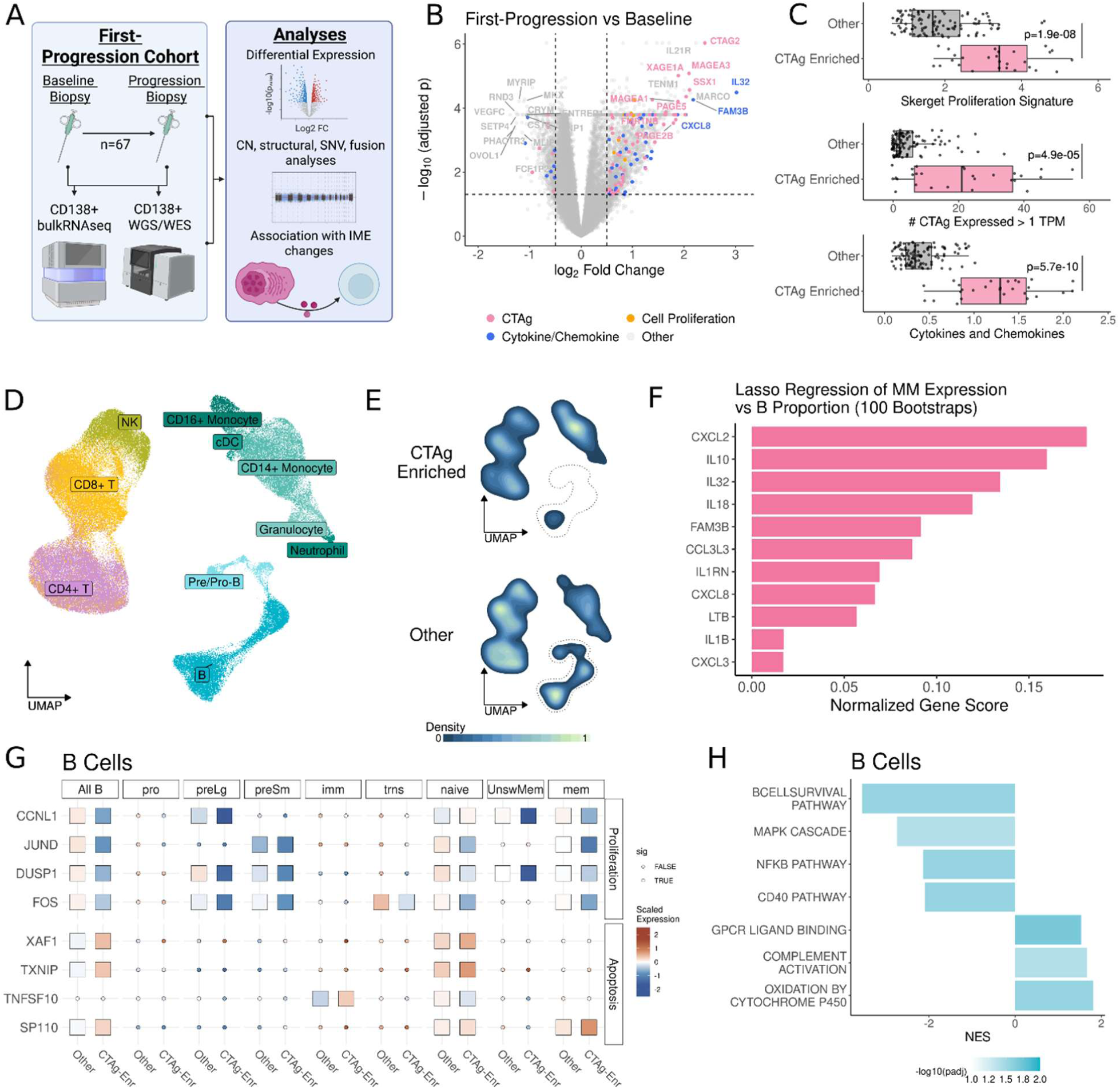
Cancer-testis antigen expression MM associates with B cell depletion at first progression. **A.** Schematic for the longitudinal analysis of CD138^pos^ bulk RNA sequencing and whole genome sequencing (n=67). **B.** Volcano plot displaying the differentially expressed genes (DEGs) comparing baseline vs first progression. Differential expression was executed using limma voom with significance cut point of absolute Log_2_ fold change > 0.5 and multiple comparisons adjusted p value < 0.05. Selected points are colored by category including cancer testis antigens (CTAg) in pink, pro-proliferative genes in orange, and cytokines/chemokines in blue. **C.** Box plots of proliferation gene signature published in Skerget et al., number of CTAgs with > 1 TPM, and enrichment for cytokine/chemokine DEGs. P value calculated by students t test. **D-E.** Uniform manifold approximation and projection (UMAP) embeddings for lymphoid cells in CD138^neg^ scRNAseq of samples from first progression and corresponding density plots displaying the normalized density of cells. Color in the density plots indicates density of cells with lighter colors indicating higher density. **F.** Bar chart displaying the cytokine/chemokine genes expressed in MM associated with naïve B depletion. One-hundred bootstrapped iterations of lasso regression were performed with cytokine/chemokine gene expression (CD138^pos^ bulk RNAseq) as the candidate predictors of proportion of naïve B cells (CD138^neg^ scRNAseq). Feature importance scores were calculated by the normalized, average r value weighted by the proportion of iterations in which the feature was included in the Lasso model. **G.** Heatmap displaying the patient-wise scaled expression of pro-proliferation and pro-apoptotic genes in B cells from first progression samples. Significance was calculated using limma-voom with Benjamini-Hochberg multiple comparison correction. Tiles are colored by normalized gene expression values were pseudo-bulked sample-wise, scaled, and averaged across samples within each group with orange for high values, blue for low values, and white for near-0 values. Shape indicates statistical significance with boxes for significant differences between CTAg enriched samples versus all other samples and circles for non-significant differences. **H.** Bar plot of gene set enrichment analysis of B cells comparing CTAg-enriched samples to all other samples. The X axis indicates normalized enrichment score (NES) and the bar color indicates -log_10_(adj. p) with grey for adj.p approaching 1.0 and blue for small adj.p values.

### Cancer testis antigen enriched myeloma associates with B cell depletion

Given that CTAg MM highly expressed several cytokines and chemokines, we hypothesized that these patients would exhibit alterations in their IME composition. Fifty-two patients had CD138^neg^ samples at first progression, with eight having CTAg-enriched MM in their CD138^pos^ bulk RNA sequencing. Strikingly, CTAg-enriched samples had significantly lower proportions of most B lineages including large pre-B (L_2_FP = -2.7, adj.padj. p = 0.028), small pre-B (L_2_FP = -2.6, adj.padj. p = 4.7e^-3^), immature B (L_2_FP = -2.8, adj.padj. p = 1.0e^-3^), naïve B (L_2_FP = -1.5, adj.padj. p = 0.028), transitional B (L_2_FP = -3.1, adj.padj. p = 1.0e^-3^), and memory B (L_2_FP = -2.8, adj.padj. p = 0.014, **Fig 5D-E, Supplemental Fig 17**). This finding was particularly notable given that naive B cells were previously found to be at higher proportion after ASCT in patients with the sustained response (**Fig 3B**). To identify which genes expressed in CTAg enriched MM associate with B cell depletion, we performed Lasso regression of cytokine/chemokine expression in MM cells with B cell proportion. Six genes explained most of the variation in naïve B proportion: *CXCL2*, *IL10*, *IL32, IL18*, *FAM3B*, and *CCL3L3* (**Fig 5F, Supplemental Fig 18A-B**). Of these, IL10(38) and IL32(39) are known to suppress B cells. Concurrently, B cells in CTAg enriched samples had increased expression of pro-apoptotic genes (*XAF1*: L_2_FC = 0.44, adj.p = 1.3.0e^-46^, *TXNIP:* L_2_FC = 0.33, adj.p = 3.4.0e^-9^, *SP110:* L_2_FC = 0.27, adj.p = 4.9e^-13^) and decreased expression of pro-proliferative genes *(JUND:* L_2_FC = -0.85, adj.p = 1.1.0e^-98^*, FOS:* L_2_FC = -0.65, adj.p = 6.1e^-42^*, CCNL1:* L_2_FC = -0.51, adj.p = 3.2e^-39^*, DUSP1:* L_2_FC = -0.40, adj.p = 2.7e^-26^, **Fig 5G**). Of the B cell subclusters, naïve cells appear to be particularly suppressed with both decreased expression of proliferative genes and increased expression of pro-apoptotic genes (**Fig 5G**). Consistent with these findings, GSEA identified significantly decreased enrichment for the B cell survival pathway in B cells from CTAg enriched samples (NES = -3.5, adj.p = 0.028) as well as B cell activation pathways such as NFKB (NES = -2.1, adj.p = 0.028, **Fig 5H**). These data suggest that CTAg-enriched MM promotes a cytokine/chemokine-driven reshaping of the IME that may suppress B cell development through direct and indirect mechanisms to support IME evasion. CTAg-enriched MM is associated with markedly low levels of B cell populations, particularly naïve B cells, potentially driven by elevated cytokines such as *IL10* and *IL32*. These samples showed increased B cell apoptosis, reduced proliferation, and suppressed survival pathways. Together, these findings suggest that CTAg expression reshapes the IME to impair B cell development, supporting immune evasion. However, several of the genes identified by Lasso regression are known to act on other cells in the IME, so we next aimed to characterize how myeloid and CD8^+^ T cells are altered in CTAg enriched samples.

### Cancer-testis antigen expressing myeloma supports myeloid derived suppressor cells and T cell exhaustion

The cytokines and chemokines over-expressed in CD138^pos^ bulk RNAseq of CTAg MM include known myeloid- attracting chemokines *CXCL2* and *CCL3*(40), as well the myeloid derived suppressor cell promoting cytokine*, IL18*(41), prompting us to evaluate the myeloid compartment as a potential indirect mechanism by which CTAg MM creates a suppressive IME. Differential expression of monocytes from CTAg-enriched samples to all other samples found hallmarks of MDSC gene expression, including increased leukocyte immunoglobulin-like receptor (LILR) genes (e.g. *LILRB4*: L_2_FC = 0.34, adj. p = 3.7e^-199^, *LILRB1*: L_2_FC = 0.40, adj. p = 4.3e^-104^) and decreased MHC-II (e.g. *HLA-DRB1*: L_2_FC = -0.51, adj. p = 2.8e^-99^, *HLA-DRA* L_2_FC = 0.30, adj. p = 2.8e^-28^, **Fig 6A-B**). Interestingly, the subclusters of classical CD14^+^ monocytes (including hypoxic, IFN stimulated, proinflammatory, and S100 enriched monocytes) which typically have a greater propensity for antigen presentation were the also the subclusters that had decreased expression of MHC class II genes, suggesting and IME with decreased potential for generating a response against CTAgs. Calculation of a monocytic MDSC gene expression enrichment score using a previously published gene signature(42) found enrichment in monocytes from CTAg-enriched samples relative to other samples (p = 1.9e^-156^, **Fig 6C**). Supporting the possible indirect role of monocytes in suppressing the IME in CTAg samples, GSEA identified significant enrichment immunosuppressive pathways including FCGR3A mediated IL-10 synthesis (NES = 2.3, adj. p = 3.7e^-3^) as well as decreased antigen presentation (NES = -3.6, adj. p = 4.5e^-3^, **Fig 6D**). These findings suggest that CTAg-expressing MM reshapes the immune microenvironment by inducing MDSC-like features in monocytes and suppressing antigen presentation, thereby fostering an immune-evasive niche that may hinder effective immune surveillance and contribute to disease progression.

**Figure 6:**
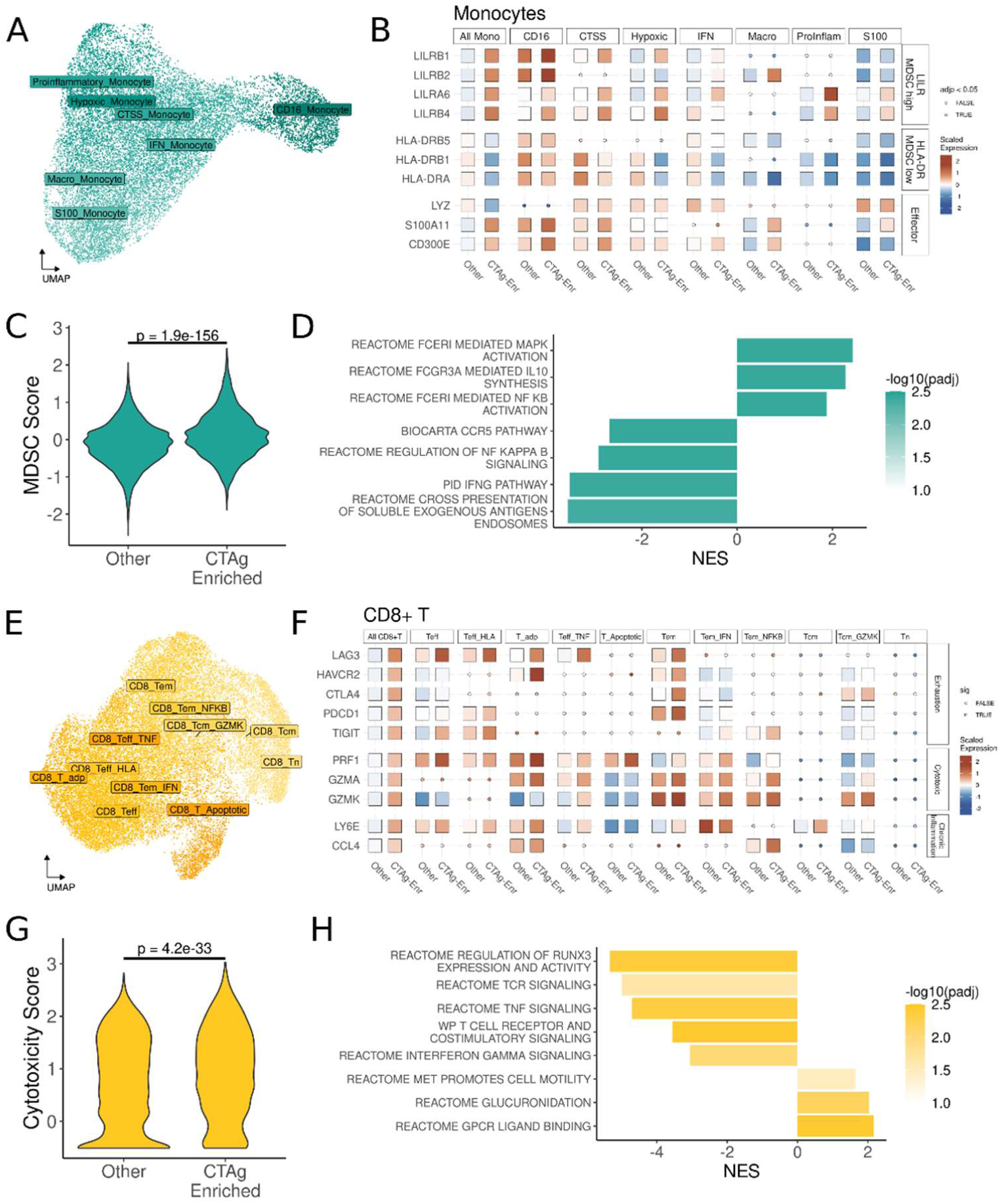
Cancer-testis antigen expressing myeloma supports myeloid derived suppressor cells (MDSC) and T cell exhaustion **A.** Uniform manifold approximation and projection (UMAP) embeddings for myeloid cells in CD138^neg^ scRNAseq of samples from first progression. **B.** Heatmap displaying the patient-wise scaled expression of MDSC-related genes in myeloid cells from first progression samples. Significance was calculated using limma-voom with Benjamini-Hochberg multiple comparison correction. Tiles are colored by normalized gene expression values were pseudo-bulked sample-wise, scaled, and averaged across samples within each group with orange for high values, blue for low values, and white for near-0 values. Shape indicates statistical significance with boxes for significant differences between CTAg enriched samples versus all other samples and circles for non-significant differences. **C.** Violin plot displaying MDSC score for myeloid cells from CTAg-enriched samples and all other samples. MDSC score calculated by the average of normalized scaled expression values for published MDSC markers. **D.** Bar plot of gene set enrichment analysis of myeloid cells comparing CTAg-enriched samples to all other samples. The X axis indicates normalized enrichment score (NES) and the bar color indicates -log_10_(adj. p) with grey for adj. p approaching 1.0 and teal for small adj. p values. **E.** Uniform manifold approximation and projection (UMAP) embeddings for CD8^+^ T cells in CD138^neg^ scRNAseq of samples from first progression. **F.** Heatmap displaying the patient-wise scaled expression of exhaustion and chronic inflammation related genes in CD8^+^ T cells from first progression samples. Significance was calculated using limma-voom with Benjamini-Hochberg multiple comparison correction. Tiles are colored by normalized gene expression values were pseudo-bulked sample-wise, scaled, and averaged across samples within each group with orange for high values, blue for low values, and white for near-0 values. Shape indicates statistical significance with boxes for significant differences between CTAg enriched samples versus all other samples and circles for non-significant differences. **G.** Violin plot displaying cytotoxicity score for CD8^+^ T cells from CTAg-enriched samples and all other samples. Cytotoxicity score calculated by the average of normalized scaled expression values for published MDSC markers. **H.** Bar plot of gene set enrichment analysis of CD8^+^ T cells comparing CTAg-enriched samples to all other samples. The X axis indicates normalized enrichment score (NES) and the bar color indicates -log_10_(adj. p) with grey for adj. p approaching 1.0 and teal for small adj. p values.

Finally, to determine if the CD8^+^ T compartment is influenced by the presence of CTAg MM we examined the gene expression in CTAg enriched samples (**Fig 6E**). Interestingly, CD8+ T cells had increased expression of cytotoxic genes including *PRF1* (L_2_FC = 0.38, adj. p = 6.6e^-69^), *GZMA* (L_2_FC = 0.49, adj. p = 4.3e^-64^), and *GZMK* (L_2_FC = 0.50, adj. p = 1.3e^-65^,**Fig 6F-G**). The cytotoxic enrichment observed appears to be an indication of prolonged stimulation, as CD8^+^ T cells from CTAg enriched samples depicted significantly higher markers chronic inflammation (*CCL4*: L_2_FC = 0.26, adj. p = 1.2e^-18^, *LY6E*: L_2_FC = 0.37, adj.p = 8.0e^-89^), and markers of exhaustion including *LAG3,* (L_2_FC = 0.32, adj. p = 2.8e^-161^), *HAVCR2* (L_2_FC = 0.28, adj. p = 1.9e^-250^), *CTLA4* (L_2_FC = 0.28, adj. p = 1.0e^-250^), *PDCD1* (L_2_FC = 0.27, adj. p = 1.0e^-250^), and *TIGIT* (L_2_FC = 0.25, adj. p = 5.3e^-88^, **Fig 6F**). This is further supported by the finding that the effector and effector memory populations had the greatest frequency of increased expression of exhaustion markers, potentially implying that an ineffective cytotoxic response in the context of an immunosuppressive IME lead to CD8^+^ T cell exhaustion. GSEA similarly identified decreased activity of canonical T cell activation pathways such as TCR signaling (NES = -4.8, adj.p = 0.021), RUNX3 activity (NES = -5.3, adj.p = 4.6e^-3^), and TNF signaling (NES = -4.7, adj.p = 4.8e^-3^, **Fig 6H**). Together, these data suggest that CTAg enriched MM profoundly reshapes the immune microenvironment by promoting immune dysfunction—depleting B cells, inducing MDSC-like monocytes, and driving CD8⁺ T cell exhaustion—thereby limiting immune-mediated disease control at relapse. These insights highlight CTAg expression as a biomarker of immune resistance and suggest that targeting CTAg-associated immunosuppressive pathways may offer new therapeutic strategies to restore anti-tumor immunity in relapsed MM.

### Patients with three or more longitudinal samples depict progressive enrichment of cancer testis antigen expression and clonality of mutations in chromatin modifying genes

To evaluate how MM genetics and transcriptomics progress through the disease course we analyzed 13 patients with three or more matched CD138^pos^ bulk RNA sequencing and whole genome sequencing samples (**Fig 7A**). Six of the 13 patients had consistent increases in the number of CTAg genes expressed (> 1 TPM) across longitudinal samples (**Fig 7B**). All six patients received a PI and steroid as a part of their induction chemotherapy regimen, with four also receiving an IMiD and two receiving a DNA alkylator (**Fig 7C, Supplemental Figs 19-23A**). Despite therapy, all six patients exhibited not only increased in expression of CTAg genes (Log_2_ TPM) but also acquired expression of a greater number of CTAg families across progressions, suggesting increasing (epi)genetic dysregulation over time (**Fig 7D, Supplemental Figs 19-23B**). Consistent with the longitudinal two-sample analysis (**Fig 5**), there were not unifying cytogenetic changes across samples, with three patients having odd-number-trisomes, four 1q gain, and three with chr13 deletion (**Fig 7E, Supplemental Figs 19-23C**). While many of these lesions are associated with poor outcomes, none were consistently observed across CTAg-expressing samples, suggesting that CTAg upregulation during disease progression is not driven by a shared cytogenetic mechanism. Instead, these data point toward an alternative driver—potentially epigenetic in nature—underlying the stepwise accumulation of CTAg expression across samples.

**Figure 7:**
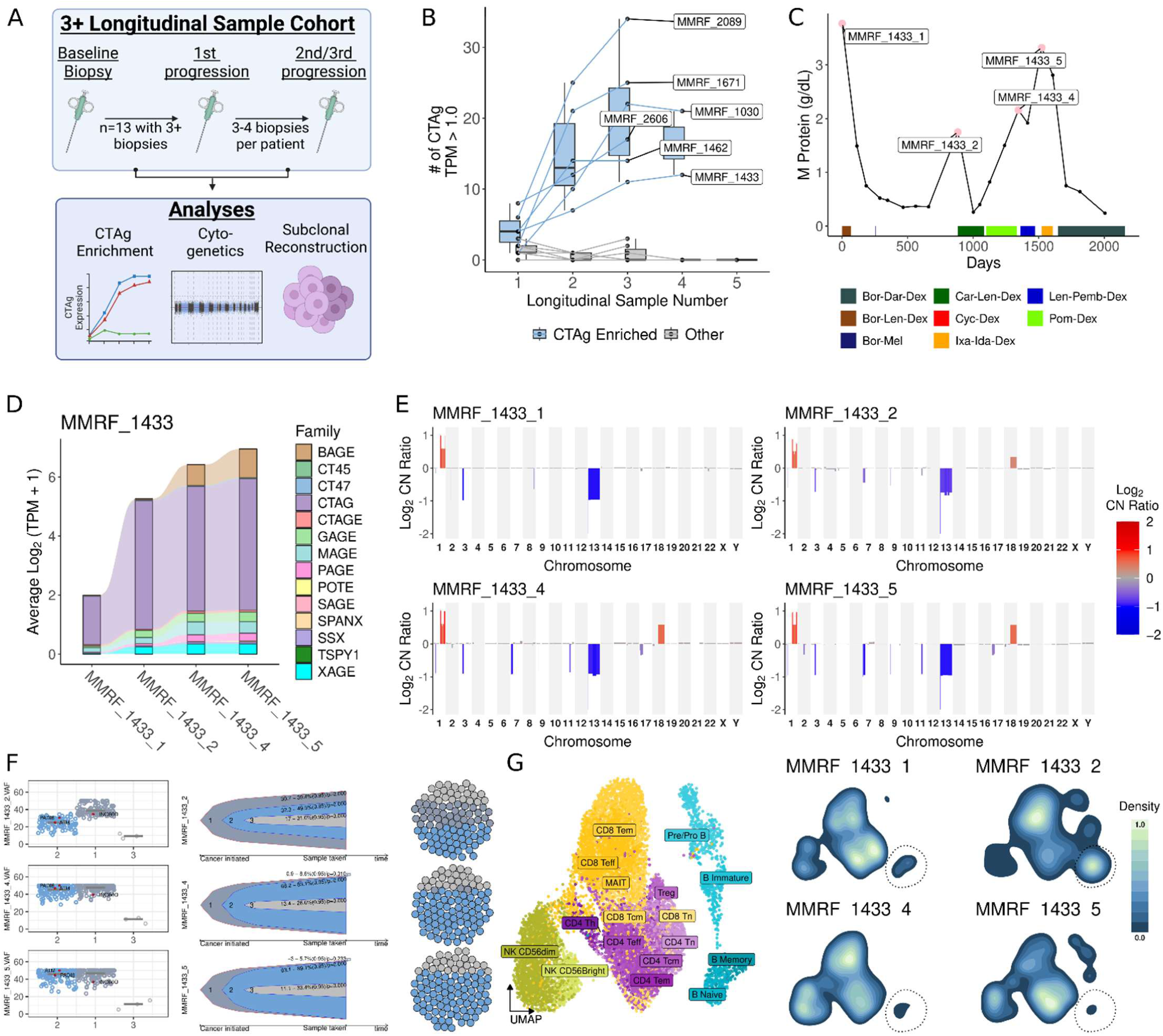
Longitudinal evaluation of patients with three or more samples in CD138^pos^ DNA and RNA sequencing. **A.** Schematic representation of longitudinal analysis of 13 patients with three or more samples. **B.** Box and line plot displaying the number of cancer testis antigen (CTAg) genes expressed at > 1 TPM in CD138^pos^ bulk RNA sequencing. Six out of thirteen samples had progressive increases in CTAg expression (blue), while seven patients had three or fewer CTAgs expressed across longitudinal samples (gray). **C.** Line plot of serum M protein measurements over time. Pink dots and labels annotate the visits at which biopsies were taken for RNA and DNA sequencing. Bars below the line indicate lines of treatment. Patient 1433 is presented as a representative example, with corresponding data for the other five patients shown in Supplemental Figures 19–23. **D.** Alluvial plot of CTAg gene expression across longitudinal samples. Alluvium height indicates the average Log_2_ transcripts per million (TPM) for genes in each CTAg family. **E.** Copy number plot split by longitudinal samples. The X axis displays chromosome number, the Y axis and bar color displays the Log_2_ copy number ratio with red for copy number gain and blue for copy number loss. **F.** Subclonal reconstruction of longitudinal samples. Subclonal structure was inferred based on the cancer cell fraction of single nucleotide variants (SNV) using QuantumClone. SNVs in genes coding for chromatin modifying proteins are labeled and highlighted in red (left). Bell plots display inferred subclonal evolution from the SNV cancer cell fractions (middle). Dot chart depicts the relative proportion of each subclone across time points (right). **G.** Uniform manifold approximation and projection (UMAP) embeddings for lymphoid cells and corresponding density plots displaying the normalized density of cells. Color in the density plots indicates density of cells with lighter colors indicating higher density. Schematic produced in BioRender.

To determine if CTAg-expressing MM cells exhibited increased chromatin modifying mutations, we performed subclonal reconstruction on the SNVs detected across timepoints. Consistent with our longitduinal two-sample analysis, four of the samples depicted persistent or increasing clonality of non-synonymous mutations in chromatin modifying genes including *TRRAP*(43), a histone acetyltransferase, *KDM3B*(44), a demethylase, *INO80D*(45), a member of the INO80 chromatin remodeling complex, *PADI6*, a peptidyl arginine deiminase has been shown to influence transcriptional state via H4 acetylation(46), as well as genes involved in general DNA stability including *ATM* and *TP53*(47) (**Fig 7F, Supplemental Figs 19-23D**). Interestingly, patient 1433 did not have mutations detected in *INO80D* or *PADI6* at baseline when CTAg expression was relatively low, then at first progression had clonal *INO80D* mutation and subclonal mutation of *PADI6* and *ATM*, with all three being clonal in subsequent samples (**Fig 7F**). These changes correlated with the progressive increase in CTAg expression, potentially indicating connection between the two. Patient 1433 also had CD138^neg^ scRNAseq at the same four timepoints. Examining their corresponding IME profile through their disease course, their first progression had naïve B cells comprising 11.7% of the IME (**Figure 7G**). However, after a brief response (**Fig 7C**), patient 1143 had a second progression after which they had persistently low naïve B cell percents (Third sample: 1.8%, Fourth sample: 0.9%). Taken together, these longitudinal analyses suggest that CTAg expression may be shaped by the clonal evolution of mutations in chromatin-modifying genes, and that such transcriptional reprogramming may be linked to IME remodeling. In particular, the case of patient 1433 illustrates a potential trajectory in which acquisition and increased clonality of chromatin regulatory mutations coincides with both increasing CTAg expression and declining naïve B cell abundance, reinforcing the proposed connection between epigenetic dysregulation and immune evasion in relapsed MM.

## Discussion

In this study we longitudinally evaluate the dynamics of myeloma and its immune microenvironment across baseline at diagnosis, response to therapy, and disease progression. By utilizing CD138^neg^ scRNAseq, CD138^pos^ bulk RNAseq, and whole genome sequencing, we provided a deep mechanistic exploration of MM biology in the BM. Our analyses build on previous studies to provide a detailed characterization of how interactions between the B lymphoid, T lymphoid, and myeloid compartments shape the IME after induction with and without ASCT(7,10). Evaluation of outcomes in patients with CD138^neg^ scRNAseq, with subsequent extension to independent derivation and validation cohorts in the CoMMpass clinical data provided compelling evidence for robust B reconstitution as a positive biomarker post-ASCT. Finally, evaluation of myeloma profiles and the IME at first (and subsequent) progression identified CTAg enrichment as associated with development of malignant disease features and immune suppression. This integrative, multi-omic approach not only advances our understanding of the evolving immune landscape in MM but also establishes a framework for identifying actionable biomarkers and therapeutic targets, paving the way for more personalized and effective treatment strategies.

By evaluating the composition of the IME in matched samples at diagnosis, first response after induction, and first response after induction and ASCT we uncovered transcriptional changes and putative intracellular signaling interactions that hinder long term memory formation during the post-ASCT reconstitution period. Consistent with previous studies(10–12), we observed that post-ASCT samples had a proportional increase in early B subclusters as well as a proportional decrease in naïve, central memory, and helper T cells. In the CD8^+^ T compartment effector and early-apoptotic cells expanded post-ASCT, whereas effector memory cells were depleted, suggesting a skew in the effector cell that develop immunologic memory. This imbalance likely contributes to sub optimal tumor surveillance during the reconstitution period and may benefit from balancing the cell fate outcomes. Cell communication, transcription factor enrichment, and GSEA depicted tumor necrosis factor and type II interferon signaling from CD16^+^ monocytes and CD8^+^ T cells, both of which promote highly inflammatory IME and promote the observed bias in CD8^+^ effector T cells(24). Therapeutically inhibiting these signaling pathways has the potential to improve the balance of effector memory formation and promote productive tumor surveillance, potentially increasing the duration of response after ASCT.

The IME is critical for regulating MM, with previous studies identifying B cell repopulation as a positive prognostic biomarker and exhausted T cells as a poor prognostic biomarker(10,11). Building on these works, we found that naïve B cells appear to be the critical subcluster supporting sustained response. Given that naïve B cells typically have departed from the bone marrow and undergone further development in a secondary lymphoid organ, their presence as higher proportion in a bone marrow biopsy suggests a larger pool of mobile naïve B lymphocytes in the circulation to undergo maturation. This was reflected in the enrichment for migratory pathways in naïve B cells from patients with greater than median PFS as compared to those with less than median PFS. We additionally found that the proportional abundance of naïve B cells associated with downstream enrichment for isotype switching and germinal center formation in memory B cells, indicating a more robust B immune reconstitution. This was further supported by the increased diversity of Ig gene expression in patients with longer than median PFS. Enhanced maturation and germinal center formation likely supports immune surveillance to maintain disease response. We found that the differences in Ig diversity were also detectable in the serum of the 408 patients in the CoMMpass clinical serum Ig measurements. Consistent with our scRNAseq analysis, we found that high Ig diversity was associated with sustained response in our equally split derivation and validation cohorts derived from this dataset. This builds on existing studies that described the improved prognosis of patients with immunoglobulin recovery after ASCT(48,49), suggesting that increased immunoglobulin diversity and interactions between specific B subsets and other cells in the IME (e.g. myeloid cells) contribute to sustained response after ASCT. These findings suggest that post-ASCT immune composition plays a critical role in patient outcomes and further supports the application of serum Ig values to enhance post-ASCT risk stratification.

Longitudinal evaluation of the CD138^pos^ compartment further enabled us to uncover mechanisms by which MM circumvents the IME at disease progression. Differential expression of longitudinal baseline and first progression samples identified enrichment for CTAgs, proliferation, and cytokine/chemokine expression. Consistent with recent work by Skerget *et al.*(37), we found that this pattern of gene expression associated with inferior outcomes and was rare at baseline (4 of 67 patients) but became much more common at first progression (23 of 67 patients). Importantly, this gene expression pattern did not correlate with common MM cytogenetic alterations, structural variants, single nucleotide variants, or fusion events, but was instead enriched in cases harboring mutations in chromatin-modifying genes. Building on existing studies implicating CTAgs as a high-risk feature of MM(35,36), we found that patients with CTAg-enriched MM exhibited profound IME dysfunction, including B cell depletion, expansion of myeloid-derived suppressor cells (MDSCs), and T cell exhaustion These tumors also expressed high levels of immunosuppressive cytokines such as IL-10, suggesting possible therapeutic avenues to counteract CTAg-driven immune suppression. Longitudinal tracking in seven patients with 3–4 sequential samples demonstrated progressive increases in both CTAg expression and the cancer cell fraction of chromatin modifier mutations over time, supporting a role for CTAgs in clonal evolution and immune escape. Together, these findings highlight CTAg-enriched MM as a distinct and aggressive disease phenotype driven by epigenetic dysregulation and immune evasion—an entity not captured by current risk stratification schemes and one that may warrant targeted immunotherapeutic intervention, such as immunization approaches and tailored monitoring strategies(37,50,51).

While this study provides novel insights into the dynamics of MM and its IME following initial diagnosis and treatment, limitations exist and additional follow-up studies are warranted to extend the findings of this project. First, the majority of samples in this study were collected at disease baseline, first response, and first progression. Therefore, additional study of MM and its IME at second and third responses/progressions has the potential to enhance our understanding of how their interaction becomes increasingly perturbed over time. Secondly, although the CoMMpass project offers a large sample size, multi-omic data, and mature clinical outcomes, it underrepresents patients treated with newer therapies such as daratumumab and chimeric antigen receptor T cells. Longitudinal studies of patients receiving these agents may uncover both shared and distinct IME dynamics compared to those treated with induction regimens primarily using PIs, IMiDs, and steroids. Third, CTAgs are pleiotropic and their functional consequences can be context dependent(52). Detailed studies to isolate the relative contribution of CTAgs to MM proliferation, IME alteration, and other disease features will provide a deeper understanding of how they contribute disease persistence. Finally, while CTAg expression is highly suggestive of chromatin dysregulation, we did not directly assay the chromatin state of CTAg-enriched MM. Follow up studies will characterize the chromatin state of CTAg-enriched MM to determine the exact mechanisms by which progressive CTAg expression is developed.

Our longitudinal analysis of one of the largest cohorts of matching CD138^neg^ scRNAseq, CD138^pos^ bulk RNAseq, and whole genome sequencing adds to the valuable insights produced by the MMRF CoMMpass study. Naïve B repopulation and diverse Ig production post-ASCT are promising biomarkers to enhance post- ASCT risk stratification. CTAg enrichment, predominantly seen at disease progression, appears to promote multi-faceted IME escape mechanisms. Ongoing studies will build on these findings to further define how these features influence patient outcomes, providing critical insights to design targeted therapeutics to enhance response to therapy.

## Methods (Supplemental)

### Materials Availability

This study did not generate new unique reagents.

### Data availability

All the single-cell raw data is available at MMRF’s VLAB shared resource (www.mmrfvirtuallab.org). Bulk RNA sequencing, whole genome sequencing, and clinical metadata are available through the MMRF Research Gateway (research.themmrf.org). Requests to access these data will be reviewed by the data access committee at MMRF and data will be released under a data transfer agreement that will protect the identities of patients involved in the study. A Seurat R object with processed UMI counts and limited metadata can be accessed at Zenodo (10.5281/zenodo.15498983).

### Ethics approval and participant consent

All samples used in this study were obtained from the MMRF CoMMpass clinical trial (NCT01454297). All procedures involving human participants adhered to the ethical standards set by the MMRF research committee. Written informed consent was obtained from all participants for the collection and analysis of biospecimens and clinical data. The study protocol was approved by the Institutional Review Board (IRB) at each participating medical center. A complete list of participating institutions is available at ClinicalTrials.gov (NCT01454297).

### Sample collection

A total of 243 CD138^neg^ bone marrow mononuclear cell (BMMC) samples were collected from 102 multiple myeloma patients with enrolled in the MMRF CoMMpass study (NCT01454297), representing the patients with two or more samples in the Immune Atlas project(7). Patients were monitored with quarterly follow-ups for up to eight years following their initial diagnosis. Eligibility criteria included suitability for either standard triplet therapy (an immunomodulatory drug, proteasome inhibitor, and glucocorticoid) or doublet therapy, with the majority receiving triplet therapy as first-line treatment. Samples were collected at both pre-therapy (baseline) and post-therapy (response or progression) time points and processed at four institutions: Emory University, Mayo Clinic Rochester, Mount Sinai School of Medicine, and Washington University.

### CD138^neg^ cells isolation and cryopreservation of cell samples

Bone marrow aspirates obtained from the Multiple Myeloma Research Consortium (MMRC) tissue bank were fractionated into CD138^pos^ (myeloma cells) and CD138^neg^ (immune and other bone marrow cells) populations using immunomagnetic cell selection targeting CD138 surface expression (automated RoboSep and manual EasySep systems, StemCell Technologies Inc.). Prior to bead-based separation, each sample was assessed for malignant plasma cell content using flow cytometry. CD138^neg^ cells were subsequently centrifuged at 400 × g for 5 minutes, and the resulting pellet was resuspended in freezing medium (90% fetal calf serum [FCS], 10% dimethyl sulfoxide [DMSO]) at a concentration of 5–30 million cells per mL. Samples were aliquoted, documented for cell concentration and storage location, and preserved in liquid nitrogen for future analysis.

### Single-cell RNA-seq sample preparation, library construction, and sequencing

To ensure high-quality and consistent single-cell data across sites, we developed a detailed 3′ single-cell RNA sequencing protocol using the 10x Genomics Chromium platform(7). Aliquots of CD138^neg^ BMME samples were thawed in a 37°C water bath, washed with warm medium, and pelleted by centrifugation at 370 × g for 5 minutes at 4°C. Cell pellets were resuspended in ice-cold phosphate-buffered saline (PBS) with 1% bovine serum albumin (BSA), and viability was assessed. Samples with viability below 90% underwent dead cell removal using the Dead Cell Removal Kit (Miltenyi Biotec Inc.). Cells were incubated with 100 µL of dead cell removal microbeads at room temperature for 15 minutes, followed by magnetic separation using an MS column or the autoMACS® Pro Separator. The eluted live cells were pelleted and resuspended in ice-cold PBS with 1% BSA.

In selected samples, 100–150 murine sarcoma cells (NIH/3T3 – CRL-1658, ATCC) were spiked into the final single-cell suspension to evaluate batch effects across processing sites. Approximately 8,000 cells were loaded per sample onto the 10x Genomics Chromium Controller, targeting the capture of up to 5,000 individual cells per sample. Reverse transcription, cDNA amplification, and library preparation were performed using the Chromium Next GEM Single Cell 3′ GEM, Library & Gel Bead Kit v2.1. During reverse transcription, full-length poly-A mRNA transcripts were barcoded with a 16-nucleotide cell barcode and a 10-nucleotide unique molecular identifier (UMI). The resulting cDNA was enzymatically fragmented and size-selected (∼400 bp) for library construction following 10x Genomics’ guidelines. Final library concentrations were quantified by qPCR (Kapa Biosystems) to ensure optimal cluster density for paired-end sequencing on the NovaSeq 6000 platform (Illumina). Sequencing was performed at a target depth of 25,000–50,000 reads per cell, yielding gene expression profiles representing approximately 1,000–4,000 transcripts per cell.

### Single-cell RNA-seq genome alignment and quality control

Single-cell RNA sequencing data were processed using Cell Ranger (v6.0.1, 10x Genomics Inc.) to demultiplex sequencing reads into FASTQ files, align reads to the human reference genome (GRCh38), and generate gene-by-cell UMI count matrices. Empty droplets were identified and removed using DropletUtils(53) (v1.14.2) with a false discovery rate (FDR) threshold of <0.001. Ambient RNA contamination was addressed using CellBender(54) (v0.3.0) with a false positive rate (FPR) of 0.01. For quality control, cells with fewer than 1,000 UMIs, fewer than 200 detected genes, or more than 20% of UMIs mapping to mitochondrial genes were excluded using Seurat(55). To correct for batch effects arising from processing sites and shipment batches, Harmony(56) (v0.1) was applied to the resulting cell embeddings and cluster assignments.

### Mouse cell removal

To identify and exclude mouse cells, raw sequencing data were additionally aligned to a combined human (GRCh38) and mouse (mm10) reference genome. Clusters in which more than 80% of cells had fewer than 95% of reads mapped to the human genome were excluded. Additionally, two samples in which over 65% of cells were classified as mouse cells were removed from downstream analysis. Cell barcodes corresponding to mouse-derived cells were filtered out from the merged single-cell dataset across all samples.

### Clustering and cell annotation

After removing mouse cells, the raw UMI counts were log-normalized with a scaling factor of 10,000. To reduce the dimensionality of the data, principal component analysis (PCA) was performed using the top 3,000 most variable genes, and the first 25 principal components were retained. Harmony was applied to these principal components to correct for batch effects, with each unique combination of processing center and shipment batch treated as a separate variable. Louvain clustering was then performed on the batch-corrected embeddings using Seurat’s clustering function to group cells with similar transcriptomic profiles. Clusters were visualized using Uniform Manifold Approximation and Projection (UMAP).

Clusters that appeared closely connected in UMAP space were grouped into five major components, referred to as compartments. These compartments were annotated using a combination of SingleR(57) and expression of cell type–specific marker genes. The five identified compartments included: T/NK (T cells and natural killer cells), B-Ery (B cells, CD34-positive progenitor cells, and erythroblasts), Myeloid (monocytes, neutrophils, and dendritic cells), Plasma (plasma cells), and Ery (erythrocytes). A small, distinct cluster of fibroblasts was also observed in the initial UMAP but was not included in any of the five main compartments.

To achieve more precise annotation of individual cell types, the analysis described above was repeated separately for each major compartment, using variable genes specific to that compartment. The resulting groups of cells are referred to as “subclusters” throughout the text. Each subcluster was manually annotated based on the expression of canonical marker genes or the top differentially expressed genes within each group. When evidence of additional heterogeneity was observed within a cluster—such as distinct patterns of marker expression—further sub-clustering was performed using the same approach. The genes used for subcluster annotation are described in detail in the Immune Atlas Cell Population Annotation Dictionary(7). Multiple clustering resolutions were tested, and the final set of subclusters was chosen to best resolve biologically meaningful subpopulations while avoiding the formation of spurious or patient-specific clusters.

### Doublet detection

Doublets were identified by flagging clusters with high predicted doublet proportions using three independent tools: DoubletFinder(58), Scrublet(59) (v0.2.3), and Pegasus (v1.8.1, https://github.com/lilab-bcb/pegasus). Scrublet was run with an expected doublet rate of 0.06 and a detection threshold of 0.2. Clusters enriched for doublets were identified when flagged by at least two of the three methods, using a false discovery rate (FDR) threshold of <0.05 based on Fisher’s exact test. These clusters were then manually reviewed and confirmed as doublets. Criteria used during manual review included the co-expression of canonical markers from unrelated cell lineages—for example, simultaneous high expression of T cell markers (*CD3, CD8A, GZMK*) and myeloid markers (*LYZ, CST3, CD14*)—as well as unusually high UMI counts relative to similar cell types. A total of 17 clusters were identified as doublets and excluded from downstream analyses.

### Bulk RNA sequencing, whole genome sequencing, and clinical data

Bulk RNA sequencing, whole genome sequencing, and clinical data were procured from the MMRF Research Gateway using the IA22 release. For analysis of bulk RNA sequencing the raw count and transcript per million (TPM) files produced by the salmon(60) pipeline for bias-aware transcript quantification. For whole genome sequencing, we used the copy number estimate files produced by the gatk(61) pipeline, single nucleotide variant and indel files, and the structural variant files produced by manta(62). Fusion transcript files were produced by starfusion(63). Clinical flat files were downloaded with the per patient file providing information on clinical covariates, the per patient visit file providing clinical parameters assessed at quarterly visits, the survival file providing progression free and overall survival data used in survival analyses, and the treatment regimen file providing information regarding lines of treatment and agents used.

### Labeling samples as baseline, response, or progression

To classify biopsies as baseline, response, or progression, we applied a modified version of the International Myeloma Working Group (IMWG) criteria (https://www.myeloma.org/resource-library/international-myeloma-working-group-imwg-uniform-response-criteria-multiple). Samples collected during a patient’s screening visit (typically labeled with a "_1," e.g., MMRF_1000_1) were designated as “Baseline.” For subsequent visits, generally scheduled at 3-month intervals, patients were assessed for response based on the IMWG criteria. A response was defined as a >50% decrease in serum M-protein or a >90% reduction in 24-hour urinary M- protein, or a reduction to <200 mg/24hr. In cases where either serum or urinary M-protein levels were unavailable, the “or” criterion was applied, as opposed to the “and” requirement in the IMWG criteria. If both serum and urinary M-protein data were missing, a >50% decrease in the difference between involved and uninvolved serum light chain levels was used to indicate a response. In instances where both M-protein and light chain measurements were unavailable, a >50% reduction in bone marrow plasma cells (with a baseline value >30%) was considered indicative of a response, in line with the IMWG criteria.

Biopsies were classified as progression based on the IMWG progression criteria, which include an increase of 25% from the lowest response value along with meeting an absolute increase threshold in any of the following clinical parameters. The absolute thresholds for progression included: serum M-protein (0.5 g/dL), urinary M- protein (200 mg/24hr), serum light chain difference (10 mg/dL), bone marrow plasma cell percentage (10%). Additionally, consistent with the IMWG criteria, progressions were also identified by the definite development of new bone lesions, increased size of existing bone lesions, soft tissue plasmacytomas, or the development of hypercalcemia (2.65 nmol/L). To account for light chain restricted disease, we applied the light chain progression criteria when M-protein data were available. In addition, to address the real-world nature of the data, we created a flag for visits when new lines of therapy were initiated. These visits were manually reviewed to identify potential label updates. For instance, if a patient experienced a rise in M-protein from 0.0 to 0.4 and a new line of therapy was intiated, the label was updated from response to progression, as the initiation of a new treatment likely indicated that the patient was in the process of experiencing progression.

Once all visits were labeled as baseline, response, or progression, additional timing labels were applied. All contiguous response visits were labeled as “First-response” until progression criteria were met, at which point visits were labeled “First-progression.” If the patient later met response criteria, they were labeled “Second-response”, and so on.

### Assessment of clinical covariates

Clinical covariates—including age, sex, induction therapy, and International Staging System (ISS) disease stage at diagnosis—were assessed for balance across comparison groups (e.g., induction vs. induction with ASCT). For continuous variables such as age, differences were tested using a two-sample Student’s *t*-test with a significance threshold of 0.05. For categorical variables such as sex, significance was assessed using a chi-squared test when all groups had *n* > 5, or Fisher’s exact test when any group had *n* ≤ 5.

### Calculating cell proportions

For comparisons of cell type or subcluster abundance, we calculated the proportion of each cell type or subcluster within each sample. To determine the proportion of a given cell type or subcluster relative to the entire immune microenvironment, we excluded doublets, plasma cells, and erythroid cells. The proportion was calculated by dividing the number of cells annotated to a specific cell type or subcluster by the total number of cells in that sample after excluding doublets, plasma cells, and erythroid cells.

### Assessment of sample-wise and patient-wise distribution of cell types

To assess the balance of patients and disease status (i.e. baseline, response, progression) across cell types, we calculated the Shannon entropy for each cell type category. For each variable (patient identifier, disease status) we calculate the relative proportion of each cell type. Then, the Shannon entropy was calculated for each cell type with higher values indicating more event distribution of cells across patients/disease status.

### Assessment of plasma cells in single cell RNA sequencing data

After CD138^neg^ sorting and scRNAseq, plasma cells were detected in the resulting dataset. To determine whether these cells represented stochastic carryover of malignant plasma cells during sorting, we performed complementary analyses. First, we assessed the correlation between the proportion of plasma cells detected by scRNA-seq and those identified by clinical flow cytometry prior to sorting using Pearson correlation. Second, to evaluate whether the plasma cells exhibited genomic features consistent with malignancy, we used the inferCNV R package to infer large-scale copy number variations (CNVs) from scRNAseq data in patient samples with known chromosomal abnormalities (e.g., 17p13 deletion, 1q gain). Only samples with ≥50 malignant (P-lineage) cells were included; low-quality and erythroid cells were excluded. InferCNV objects were constructed using raw gene expression counts, reference annotations (B, T, NK, and myeloid cells), and gene position data from Gencode v27 (hg38). CNV inference was performed with a 0.1 expression cutoff, denoising enabled, and CNV calling using the HMM “i3” model.

### Differential abundance on cell type/subcluster proportions

To assess changes in cell type proportions over time (e.g., from baseline to first response) and how these changes differed between groups (e.g., induction vs. induction with autologous stem cell transplant [ASCT]), we applied linear mixed-effects models. The primary objective was to identify differences in temporal trends between groups. For instance, the model would highlight a cell type that increases from baseline to first response in the induction-only group while either decreasing, remaining stable, or changing at a significantly different rate in the induction + ASCT group.

To achieve this, we modeled cell type/subcluster proportions as a function of time, the covariate of interest (e.g., treatment group), and their interaction. Additional fixed effects, such as age and sex, were included as appropriate to account for clinical covariates, while patient identifiers were incorporated as random effects to adjust for inter-individual variability. Post-hoc Tukey tests were conducted to identify significant differences in specific comparisons (e.g., baseline vs. first response within the induction-only group). Statistical significance was defined using a Benjamini-Hochberg (BH) corrected p-value threshold of <0.05.

Cell types or subclusters with significant differences were identified based on two criteria: (1) an adjusted p- value < 0.05 for the interaction term, indicating a difference in how proportions changed over time between groups, and (2) an adjusted p-value < 0.05 for at least one timepoint comparison in post-hoc testing. These criteria ensured that only cell types with distinct temporal trends between groups were highlighted, while excluding those with similar changes over time or no significant change at all.

### Calculation of cell density in uniform manifold approximation and project plots

To visualize the relative abundance of cell subclusters across time points and patient groups in UMAP space, we used geom_density_2d in ggplot2(64). This function estimates a two-dimensional kernel density from the UMAP coordinates to produce contour lines representing regions of similar point density. It applies a Gaussian kernel to calculate the local density of points across the UMAP projection, effectively highlighting clusters or areas with higher concentrations of observations.

### Differential expression between time points and clinical groups

Differential expression was executed using linear models in the limma package(65). For comparisons using paired samples (e.g. comparing baseline to first response for longitudinal samples from the same patients) the model included patient identifier as a covariate. For comparisons between patients, models were adjusted for technical covariates such as processing site. Statistical significance was determined using a moderated t-test statistic with BH multiple comparison correction. An adjusted p value (adj. p) cut point of 0.05 was used for all comparisons.

### Gene set enrichment analysis

To identify pathways enriched in specific cell types or subclusters across time points (e.g., baseline vs. first response) or patient groups (e.g., patients with progression-free survival above vs. below the median), we performed gene set enrichment analysis (GSEA). Ranked gene lists were generated using -log_10_ p-values from limma, with the sign determined by the log_2_ fold change (see *Differential expression between time points and clinical groups*). As a result, genes with low p-values and positive fold changes were positioned at the top of the list, while genes with low p-values and negative fold changes were placed at the bottom. GSEA was conducted using the ReactomePA(66) and fgsea(67) packages in R.

### Transcription factor activity analysis

Transcription factor (TF) activity was inferred using the decoupler package(68). This method leverages collecTRI, a curated resource integrating data from 12 transcriptional regulatory databases that list TFs, their known targets, and interaction weights—where +1 indicates activation and −1 indicates repression. For each cell, decoupler fits a univariate linear model regressing the normalized expression values of a TF’s target genes against the corresponding interaction weights. The resulting t-statistic for the slope is used as the TF activity score, with positive values indicating greater concordance with activation and negative values indicating repression or inactivity. To test for significant differences in TF activity between groups, we used dc.rank_sources_groups with the method "t-test_overestim_var". This function performs a t-test for each TF across groups while conservatively estimating variance to account for group-specific variability and sample size differences. The resulting p-values are adjusted for multiple testing by the BH approach, enabling robust identification of TFs with significantly different activity between conditions.

### Cell-cell communication analysis

To identify changes in cell–cell communication within the IME across timepoints (e.g., baseline vs. first response after ASCT), we used the CellChat package(23). Separate CellChat objects were constructed using count matrices of immune cells from each timepoint. Prior to analysis, biologically similar subclusters were aggregated to improve interpretability and statistical power—for example, all CD14⁺ monocyte subclusters were combined into a single CD14⁺ monocyte group. CellChat infers intercellular communication by integrating known ligand–receptor interactions with single-cell transcriptomic data. For each sender–receiver cell pair, CellChat calculates a communication probability based on the expression levels of ligand and receptor genes, incorporating pathway-specific signaling models. After building individual communication networks for baseline and first response, we merged the objects and applied paired comparison functions within CellChat to identify signaling pathways and cell-type interactions that significantly differed between timepoints.

### Patient stratification by post ASCT progression free survival

After characterizing changes in IME composition and gene expression following ASCT, we next sought to determine which alterations were more frequently observed in patients with sustained responses compared to those who experienced earlier progression. To stratify patients by response duration, we calculated the median progression-free survival (PFS) among the 69 patients in our cohort who received ASCT, which was 1,491 days. Patients were then categorized into three groups: those with greater than median PFS (GMpfs), defined as progression or censoring after 1,491 days; those with less than median PFS (LMpfs), defined as disease progression before 1,491 days; and a third group labeled "Censored", consisting of patients censored prior to 1,491 days, who were excluded from downstream comparisons. Clinical covariates were evaluated between the GMpfs and LMpfs groups to assess baseline differences. Differential abundance, gene expression, and gene set enrichment analyses (GSEA) were performed as previously described (see *Differential abundance on cell type/subcluster proportions, Differential expression between time points and clinical groups*, *Gene set enrichment analysis*) to identify features associated with prolonged response.

### Immunoglobulin gene expression diversity analysis

GSEA revealed that naïve B cells from patients with GMpfs were enriched for pathways related to activation and migration, while memory B cells were enriched for germinal center formation and isotype switching (**Fig 3D**). Based on these findings, we hypothesized that increased proportions of naïve B cells following ASCT are associated with greater diversity in immunoglobulin (Ig) expression across naïve and memory B cells. To test this hypothesis, we calculated the Shannon diversity index on Ig gene expression and correlated the resulting diversity scores with the log_2_ fold change in naïve B cell proportion (first response vs. baseline).

For each cell, we computed the average normalized expression of genes corresponding to each Ig isotype— IgG, IgE, IgD, IgA, and IgM—yielding five values per cell. The cell-wise values were then pseudo-bulked by averaging expression across cells from each patient. To ensure equal weighting of isotypes, we normalized each isotype’s expression to its mean across all patients, generating five normalized Ig expression values per patient. Shannon diversity was then calculated on these five values to produce a single Ig diversity score per patient.

To assess whether naïve B cell expansion was associated with Ig diversity, we performed Pearson correlation between the log_2_ fold change in naïve B cell proportion and the corresponding Shannon index, with statistical significance defined as p < 0.05.

To identify additional immune subclusters that may support Ig diversity, we extended this analysis to other cell types in the IME. We included only subclusters with at least 10 cells per patient to avoid inflated log_2_ fold changes due to low abundance. For each subcluster, we computed the log_2_ fold change in abundance (first response vs. baseline) and correlated it with the Ig diversity score across patients using Pearson correlation. Subclusters with a BH-adjusted p < 0.05 were considered significantly associated with B cell Ig diversity and were further analyzed by differential gene expression to identify potential regulatory factors (e.g. surface receptors or ligands), that may support Ig diversification in B cells.

### Overview of serum immunoglobulin diversity analysis

Patients enrolled in CoMMpass underwent study visits every three months, during which clinical parameters were assessed following therapy. To evaluate whether Ig diversity after ASCT was associated with patient outcomes (e.g. PFS, OS) we applied the Shannon diversity index to serum immunoglobulin measurements collected at these three-month intervals. Of the 1,143 patients included in the IA22 CoMMpass clinical metadata release, 408 met the following inclusion criteria: (1) received a single ASCT, (2) available PFS and OS data, and (3) had complete clinical covariate information, including age, ISS stage at diagnosis, and induction regimen.

### Calculation of post ASCT Shannon diversity index on serum immunoglobulin values

In our CD138^neg^ scRNAseq data from post-ASCT biopsies, we observed that patients with GMpfs had elevated proportions of naïve B cells, which were associated with increased diversity in Ig gene expression. We aimed to extrapolate our findings in scRNAseq to clinical Ig measurements by calculating the Shannon diversity index for each patient using serum IgM, IgG, and IgA values collected within one year of ASCT. For patients with multiple study visits during this period, we computed the mean of their visit-wise Shannon index values.

To align this analysis with our hypothesis—derived from CD138^neg^ scRNAseq data—we excluded patients whose elevated serum Ig levels were likely driven by monoclonal Ig production from MM cells. Specifically, as a conservative filter, we excluded patients whose average serum IgG, IgM, or IgA values exceeded 1.5 times the upper limit of normal (Upper limits of normal, IgG: 16.0 g/L, IgM: 2.3 g/L, IgA: 4.0 g/L) within one year post-ASCT, as these abnormal values are most likely indicative of relapsed myeloma rather than biological diversity in non-malignant plasma cells. This step was necessary to remove patients with likely disease recurrence, who would exhibit low Ig diversity and low PFS, which could otherwise bias the results and artificially strengthen associations between higher Shannon index values and improved outcomes.

### Partitioning into derivation and validation cohorts

To robustly analyze the 408 patients, we generated derivation and validation cohorts, allowing cut points identified in the derivation cohort to be tested independently in the validation cohort to confirm that observed trends generalize to unseen data. To ensure balanced distribution of disease severity, we used the caret package(69) to perform a stratified random split, maintaining equal proportions of ISS stage between the derivation (n = 205) and validation (n = 203) cohorts. As in previous analyses, we evaluated clinical covariate balance between cohorts, including age, sex, BMI, and induction therapy, to rule out potential confounders.

### Identification of a Shannon index cutpoint in the derivation cohort

The cutP method in the survminer(70) package was used to identify the optimal Shannon index cutpoint that maximized the difference in PFS. This approach systematically tests possible cutpoints and selects the one that yields the highest log-rank test statistic, while also enforcing a minimum group size to avoid overly small subgroups. CutP was applied to the derivation cohort, and patients with values above the cutpoint were labeled “High Shannon Index,” while those with values below were labeled “Low Shannon Index.” The cutpoint identified in the derivation cohort was then applied to the validation cohort to assess whether it generalized to independent data.

### Univariate and multivariate survival analysis

To assess whether differences in Shannon index were associated with patient outcomes, we used Cox proportional hazards models to compare OS and PFS between High and Low Shannon Index groups. Univariate models were first constructed using the survival package(33), with Shannon group as the predictor and either PFS or OS as the outcome. Separate models were fit for OS and PFS; hazard ratios (HRs) and corresponding p-values were calculated using a significance threshold of 0.05. Then to account for disease stage at diagnosis, we then stratified patients by both Shannon group and ISS stage. Stratified models were fit within each ISS stage (e.g. ISS Stage I with low Shannon index versus ISS Stage I with high Shannon index) to compare survival between High and Low Shannon Index groups, allowing us to estimate HRs and p-values specific to each stage.

### Longitudinal evaluation of immunoglobulin diversity from ASCT to progression

To evaluate the dynamics of immunoglobulin diversity from ASCT to progression in the High and Low Shannon groups, we calculated the Shannon index on serum immunoglobulin measurements collected at clinical visits between ASCT and each patient’s first progression. Of the 408 patients, 215 experienced progression during the CoMMpass study period and were included in this analysis. To visualize longitudinal trends on a consistent scale, visit days were normalized such that the day of ASCT was designated as day 0, and subsequent visit days were scaled relative to each patient’s PFS to produce a “percent of time from ASCT to progression.” For visualization, these percentages were binned in 10% intervals (0–10%, 10–20%, etc.). Additionally, for patients who had an increase in naïve B cell proportion (greater than the lower tertile based on differential abundance analysis) from baseline to first response post-ASCT in scRNAseq data, we visualized longitudinal Shannon index values alongside the corresponding naïve B cell proportions.

### Differential expression of longitudinal CD138^pos^ bulk RNA sequencing samples

To identify changes in gene expression from baseline to first progression, we performed differential expression analysis on 67 paired CD138^pos^ bulk RNA-seq samples. Count matrices were obtained from the CoMMpass IA22 data release using outputs generated by the Salmon pipeline. Differential expression was assessed using linear models in the limma package(65), using covariates of timepoint (baseline vs. first progression) and patient to account for pairing. Statistical significance was determined using a moderated t-statistic with BH correction for multiple testing, applying a significance threshold of 0.05. In the corresponding volcano plot, cytokine and chemokine genes were annotated based on Gene Ontology terms GO:0005125 (cytokine activity) and GO:0008009 (chemokine activity). Proliferation-related genes were highlighted using the high-risk “Proliferative” gene signature described by Skerget et al(37).: *TYMS*, *TK1*, *CCNB1*, *MKI67*, *CKS1B*, *TOP2A*, *UBE2C*, *ZWINT*, *TRIP13*, and *KIF11*. GSEA was performed on the rank ordered gene list of log_2_ fold change signed –log_10_ p value using the same methods as described in *Gene set enrichment analysis*.

### Cancer testis antigen gene selection

Cancer testis antigens (CTAgs) are genes that encode proteins typically restricted to germ cells of the testis and placenta but are aberrantly reactivated in a variety of cancers. In malignant contexts, CTAgs can exhibit pleiotropic and context-dependent biological effects. Many potential CTAgs have been identified across sequencing studies; we employed a conservative, multi-faceted approach to define a high-confidence gene set of CTAgs. First, we curated a list of candidate CTAgs from 14 publications and databases, yielding an initial pool of 575 genes(71–84). This list was then refined using the following criteria: (1) presence in at least two of the 14 sources, (2) localization to the X chromosome to enrich for genes highly likely to be subject to epigenetic silencing in somatic tissues, (3) inclusion in CTAg families with at least three members (e.g., *MAGE* genes) as these genes are typically found in close proximity on the X chromosome, and (4) expression in fewer than 50% of CD138^pos^ bulk RNA-seq samples (TPM > 1.0). Applying these filters resulted in a final set of 154 CTAgs spanning 14 gene families, representing a high-confidence set of CTAgs for downstream analysis.

### Gene module scoring in CD138^pos^ bulk RNA sequencing

To identify samples enriched for specific gene categories, we applied a gene module scoring approach. For each gene list of interest, TPM values were log-transformed with a pseudocount of 1 and averaged across genes to generate a module score for each sample. Although differential expression analysis was performed across all 67 samples, many of the differentially expressed genes were upregulated in fewer than half of the samples (**Supplemental Fig. 13**), suggesting a subset of samples were responsible for the differential expression of CTAgs. To identify these samples, we calculated a module score using the set of genes with log_2_ fold change > 0.5 and adj. p < 0.05. Samples with module scores greater than one standard deviation above the mean were considered enriched. Given that many of the top differentially expressed genes were CTAgs, we designated these high-scoring samples as “CTAg-enriched.” Module scores were also calculated for proliferation genes and for the differentially expressed cytokine and chemokine genes (see *Differential expression of longitudinal CD138^pos^ bulk RNA sequencing samples*). To evaluate whether these pathways were elevated in CTAg-enriched samples, module scores were compared between CTAg-enriched and other samples using Student’s t-test with a significance threshold of 0.05. GSEA was performed using the ReactomePA(66) and fgsea(67) packages in R on ranked gene lists generated by the -log_10_ p-values from limma with the sign determined by the log_2_ fold(66,67).

### Survival analysis between patients with and without enrichment for cancer testis antigen gene expression

To evaluate whether CTAg enrichment was associated with differences in patient outcomes we used Cox proportional hazards models to compare PFS and OS between CTAg-enriched and non-enriched groups. HRs and corresponding p-values were calculated with a significance threshold of 0.05. At baseline, four biopsies were identified as CTAg-enriched, compared to 63 non-enriched samples. For this comparison, PFS and OS were calculated from the time of diagnosis. In a separate analysis based on biopsies collected at first progression from the same 67 patients, 23 samples were CTAg-enriched and 44 were not. Because these labels were derived from samples taken at first progression, we assessed time from first to second progression as a measure of PFS, while OS remained defined from the time of initial diagnosis. This approach enabled us to assess whether CTAg expression at different disease stages was predictive of outcomes.

### Comprehensive evaluation of cytogenetics and mutations in CD138^pos^ whole genome sequencing and gene fusions from CD138^pos^ bulk RNA sequencing

To comprehensively evaluate genetic alterations including copy number alterations (CNAs), structural variants (SVs), fusion transcripts, and single nucleotide variants (SNVs), we performed an integrative analysis similar to that described by Skerget et al.(37). The approach systematically evaluates each sample and gene for these cytogenetic changes and mutations, generating a series of gene-by-sample matrices indicating the presence or absence of each type of alteration. Separate matrices were created for each alteration type, enabling downstream identification of genes affected by multiple mechanisms (e.g., *TP53* loss through both monoallelic deletion and nonsynonymous SNVs). All analyses were conducted using data from the CoMMpass IA22 release, as described in the following sections.

For CNAs, we used the gene-by-sample matrix based on the lowest log_2_ segment mean in tumor samples relative to their matched normal (somatic) controls. Thresholds for identifying specific CNA states were derived based on theoretical log_2_ values corresponding to a scenario where 80% of tumor cells carry the alteration and 20% remain diploid. For instance, the cutoff for monoallelic loss was set at -0.737, calculated as log_2_((0.8 × 1 + 0.2 × 2) / 2). Separate flags were assigned for monoallelic and biallelic loss. High-level amplifications were defined using a segment mean threshold of 1.379, corresponding to 80% of cells with a copy number of 6, consistent with Skerget et al..

For gene fusion detection, we utilized both StarFusion and PairoScope outputs to identify genes altered by fusion events. From the StarFusion dataset, we filtered fusion transcripts to include only those with a fusion fragment per million (FFPM) ≥ 0.1, at least 5 junction reads, at least 5 spanning reads, and strong anchor support. Both genes involved in each high-confidence fusion event were flagged as altered in the corresponding sample. We then updated the gene-by-sample fusion matrix by incorporating the results from PairoScope, which specifically targeted known multiple myeloma driver fusions involving *NSD2, CCND1, MYC, MAFA, CCND2, MAF,* and *MAFB*. Any sample with a positive PairoScope fusion call for one of these genes was marked positive in the gene-by-sample matrix.

For structural variants, we used the output files generated by the Manta pipeline. All indels were filtered to retain those with a mapping quality of at least 20, paired-end support (PE) of at least 5, split-read support (SR) of at least 3, an absolute CIPOS less than 50, marked as precise, and a tumor genotype quality score of at least 30. The filtered structural variant file was then parsed into separate gene-by-sample matrices for deletions, translocations, and inversions.

For SNVs, we applied a multi-layered approach. To first inclusively identify SNVs, we used the gene-by-sample matrix of non-synonymous SNV counts. Separate flags were created for the presence of one non-synonymous SNV and for two or more non-synonymous SNVs. To further identify SNVs that occur frequently, we added two additional flags using the variant call format (VCF) files, following a method similar to Skerget et al. First, we applied quality filters to retain SNVs with at least 10 reads in both tumor and reference samples, an allelic ratio above 0.05, detection by at least two of five variant callers (MUTECT2, STRELKA2, VARDICT, OCTOPUS, LANCET), and a predicted impact of either "HIGH" or "MODERATE." From this filtered VCF file, mutations were flagged as recurrent if the same amino acid alteration occurred in at least two patients, or as clustered if at least five patients had non-synonymous mutations within 10% of the coding DNA sequence of the gene.

Finally, the separate gene-by-sample matrices for each mutation and cytogenetic alteration type were consolidated into a summary data frame in long format. For each gene-sample entry that was flagged, a row was created containing the sample name, gene, and type of mutation. For example, each flagged entry in the monoallelic loss gene-by-sample matrix became a row in the summary data frame with its corresponding sample name, gene, and the label “monoallelic loss.” This process was repeated for each type of mutation described above, resulting in a comprehensive summary of copy number alterations, fusion transcripts, structural variants, and SNVs used in downstream analyses.

### Identification of mutations in genes specific to samples with cancer testis antigen enrichment

To explore the possible mutations and cytogenetic changes associated with CTAg gene enrichment, we evaluated the prevalence of known MM-driving mutations. First, we compared the proportion of samples labeled as CTAg-enriched against those that were not for cytogenetic changes or mutations in: *CCND1, CCND2, TP53, CYLD, DIS3, FGFR3, KRAS, NRAS, MAF, MAFA, MAFB, BRAF, MYC, NSD2,* and *RB1*. We also evaluated the copy number state of each chromosome in CTAg-enriched samples. For the copy number heatmap, we transformed log_2_ segment_mean values to CN by using the formula 2 × (2^log2_segment_mean^).

Given that MM-driving mutations did not differentiate CTAg-enriched samples from non-CTAg-enriched samples, we next aimed to identify mutations that may contribute to CTAg enrichment. Since CTAgs are typically epigenetically repressed, we hypothesized that mutations in chromatin-modifying genes might be more prevalent in CTAg-enriched samples. We first identified a large list of candidate genes using the GO terms related to chromatin modification (GO:0016569, GO:0016570, GO:0006338, GO:0034728). For each gene, we calculated the proportion of samples in the CTAg-enriched and non-CTAg-enriched groups with any mutation (CNA, fusion, SV, CNV). To identify genes present more frequently in CTAg-enriched samples, we filtered to the genes that were present in less than 2% of non-CTAg samples and were present in at least twice the proportion of CTAg-enriched samples compared to non-CTAg-enriched samples. To isolate the minimum list of genes that maximized the difference in the proportion of CTAg-enriched samples versus non-CTAg-enriched samples, we performed recursive feature elimination. Starting from the list of genes that met the filtering criteria, we iteratively tested removing each gene and subsequently calculated the proportion of CTAg-enriched samples containing at least one mutation in the list and the proportion of non-CTAg-enriched samples with at least one mutation in the list. After testing all genes, the gene that resulted in the largest increase in the ratio of the proportion of CTAg-enriched samples to non-CTAg-enriched samples with at least one mutation was excluded. This process was repeated until no further gene exclusions increased the specificity of the gene list for CTAg-enriched samples versus non-CTAg-enriched samples.

### Differential abundance and expression of immune subclusters at first progression

To assess whether there were differences in the proportion of immune cells at first progression in the CD138^neg^ scRNAseq data, we compared patients identified as CTAg-enriched in their CD138^pos^ bulk RNA sequencing to those without CTAg enrichment. For each immune subcluster, we tested for differences in proportion using a linear model of proportions between CTAg-enriched samples and other samples. The log_2_ fold change in proportion was calculated by dividing the mean proportion in the CTAg-enriched group by the mean proportion in all other samples, and taking the log_2_ of that ratio. Statistical significance was determined using a BH corrected p-value threshold of < 0.05. Differential expression was performed using limma with a linear model comparing CTAg-enriched samples to all other samples. Statistically significant differences were defined by a BH corrected p-value of < 0.05.

### Lasso regression of CD138^pos^ cytokine and chemokine expression with naïve B proportion

Given that the proportion of naïve B cells was found to be increased in patients with greater than median PFS and decreased in samples with CTAg-enriched multiple myeloma (MM), we next aimed to identify genes expressed in CD138^pos^ bulk RNA sequencing of CTAg-enriched MM that are associated with the proportion of naïve B cells in CD138^neg^ scRNAseq. To generate a candidate list of genes potentially influencing naïve B cell abundance, we used GO lists of cytokines (GO:0005125) and chemokines (GO:0008009) that were differentially expressed in CTAg-enriched samples (log_2_ fold change > 1.0 and adjusted p-value < 0.05). This yielded 53 genes, which were included as potential predictors of naïve B cell proportion. To robustly test the association between gene expression and naïve B cell abundance, we performed 100 iterations of bootstrapped Lasso regression. In each iteration, the 27 samples with matched CD138^pos^ bulk RNAseq and CD138^neg^ scRNAseq were randomly resampled with replacement. Lasso regression was then performed between the log_2_-transformed TPM values of the 53 candidate cytokine/chemokine genes and the proportion of naïve B cells.

After completing 100 bootstrapped iterations, we next identified genes with the most consistent association with naïve B cell proportion. For each gene, we calculated a normalized gene score by multiplying the average Lasso regression coefficient by the frequency with which the gene was selected as a significant predictor of naïve B abundance (i.e., the number of bootstraps in which it was selected divided by 100). These scores were then normalized across all candidate genes to a 0-1 scale. Genes were ranked by their normalized gene scores to estimate their relative importance in association with naïve B cell proportion.

To identify the minimal set of cytokine and chemokine genes that collectively associate with the majority of the variance in naïve B cell proportion, we performed recursive feature addition. Beginning with the gene with the highest normalized score, we iteratively constructed 100 bootstrapped Lasso regression models and calculated the resulting R value for each model. The bootstraps were used to calculate mean and standard error on R values for each model. Then genes were added to the growing gene list in order of descending normalized gene score, up to a total of 11 genes. To estimate the point at which model performance plateaued, we fit a saturating exponential model to the curve of the number of genes versus R value. The resulting gene list was considered the minimal and most informative set of cytokine and chemokine genes associated with naïve B cell proportion.

### Gene set enrichment analysis and module scoring in CD138^neg^ scRNAseq at first progression

To identify biological pathways enriched in samples from patients with CTAg-enriched multiple myeloma, we performed gene set enrichment analysis (GSEA) using the fgsea package. Ranked gene lists were generated from the limma differential expression results using -log_10_ p-values, with the direction of change indicated by the sign of the log_2_ fold change. In addition to pathway-level enrichment, we calculated signature scores T cell cytotoxicity and myeloid derived suppressor cells(42) (MDSC) based on previously published gene sets. The T cell cytotoxicity score was computed as the average of log-normalized expression values for the following genes: *GZMB, PRF1, GZMA, GZMH, GNLY,* and *NGK7*. Statistical significance was assessed using limma as described above. The MDSC score included both positive (*LILRB4, LILRB2, LRFN4, WASF1, TPBG, F12, CD274*) and negative (*HLA-DRA, HLA-DRB1, HLA-DRB5, CD83, PRTM8, GIMAP6*) markers. Scores were calculated similarly to the cytotoxic signature, except that the expression values of the negative markers were multiplied by -1 prior to averaging across all markers.

### Evaluation of cancer testis antigen gene expression in CD138^pos^ bulk RNA sequencing for patients with three or more longitudinal samples

To jointly assess the longitudinal dynamics of gene expression and genomic alterations, we analyzed CD138^pos^ bulk RNA sequencing and whole genome sequencing data from 13 patients with three or more longitudinal samples in both assays. To evaluate CTAg expression over time within each patient, we applied two complementary approaches. First, we quantified the number of CTAgs (from the curated list of 154 genes; see *Cancer testis antigen gene selection*) with expression greater than 1 TPM as a measure of epigenetic escape. A threshold of more than three expressed CTAgs emerged as an empiric cutoff distinguishing patients with persistently low expression from those showing increasing CTAg expression over time. Second, we calculated the average log_2_ TPM (with a pseudocount of 1) across CTAg naming families for each sample. This approach provided a summary measure of CTAg expression, enabling characterization of both breadth and magnitude of expression across timepoints.

### Subclonal reconstruction in patients with three or more CD138^pos^ bulk whole genome sequencing samples

To infer subclonal progression of malignant cells across longitudinal samples, we performed subclonal reconstruction analysis on the CD138^pos^ whole genome sequencing data from six patients who had three or more samples and progressive enrichment for CTAg genes. For each sample, we used the SNV data from the IA22 CoMMpass release. Hot spot mutation regions in the immunoglobulin heavy chain, kappa light chain, and lambda light chain genes were excluded from the analysis. These regions included chromosome 14 (from 105,000,000 bp to the end of the chromosome), chromosome 2 (between 87,000,000 and 90,000,000 bp), and chromosome 22 (between 21,000,000 and 23,000,000 bp). To reduce the influence of technical false negatives, we limited the SNV list to those mutations present in all samples for that patient. Given that the primary objective was to identify subclonal dynamics related to mutations in chromatin-modifying genes, analyses were initiated with samples that exhibited mutations acquired during disease progression.

To account for copy number alterations, each SNV was assigned a copy number state based on the estimates produced by GATK. The thresholds for log_2_ segment mean values were derived from the theoretical value where 80% of tumor cells would exhibit the given copy number. For instance, the threshold for monoallelic loss was set at -0.737, corresponding to the expected reads if 80% of the cells were monoploid and 20% were diploid.

SNV clustering was performed using the One_step_clustering function in QuantumClone(85) with the "Flash" preclustering method and clone rangers set from 1:2 to 1:4 to minimize the identification of subclones driven by fewer than 10 SNVs. Clonal models were generated using the infer.clonal.models function in the clonevol(86) package with 1000 bootstrapped models. Clonal models were visualized using the plot.clonal.models function with annotation for missense and nonsense mutations in genes for chromatin modifying proteins.

