## Supplemental Figures for "Longitudinal multi-omic profiling uncovers immune escape and predictors of response in multiple myeloma"

**Supplemental Figures and Legends**

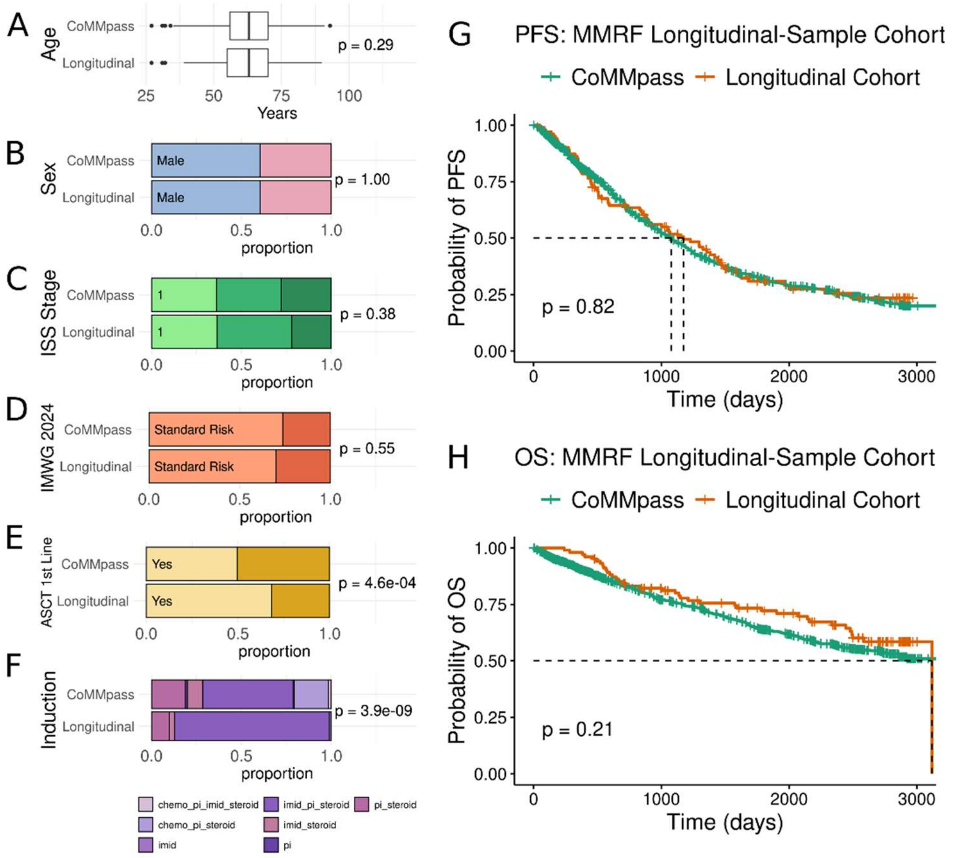


**Figure S1:** Comparison of clinical features of longitudinal cohort and CoMMpass as a whole. **A.** Box plot of patient age at diagnosis for the longitudinal cohort (n = 102) and the CoMMpass dataset (n = 1,143). P value was calculated using student's t test. **B-F.** Bar plots for the proportion of samples by sex (B), international staging system (ISS) disease stage (C), international myeloma working group risk category (D), autologous stem cell transplant (E), and induction treatment (F). P values were calculated using chi squared test. G-H Kaplan meier plot displaying the progression free survival (G) and overall survival (H) of patients in the longitudinal cohort and CoMMpass. The X axis indicates number of days from diagnosis, the Y axis indicates the proportion of patients without progression/death. P value was calculated using a Cox proportional hazards model.

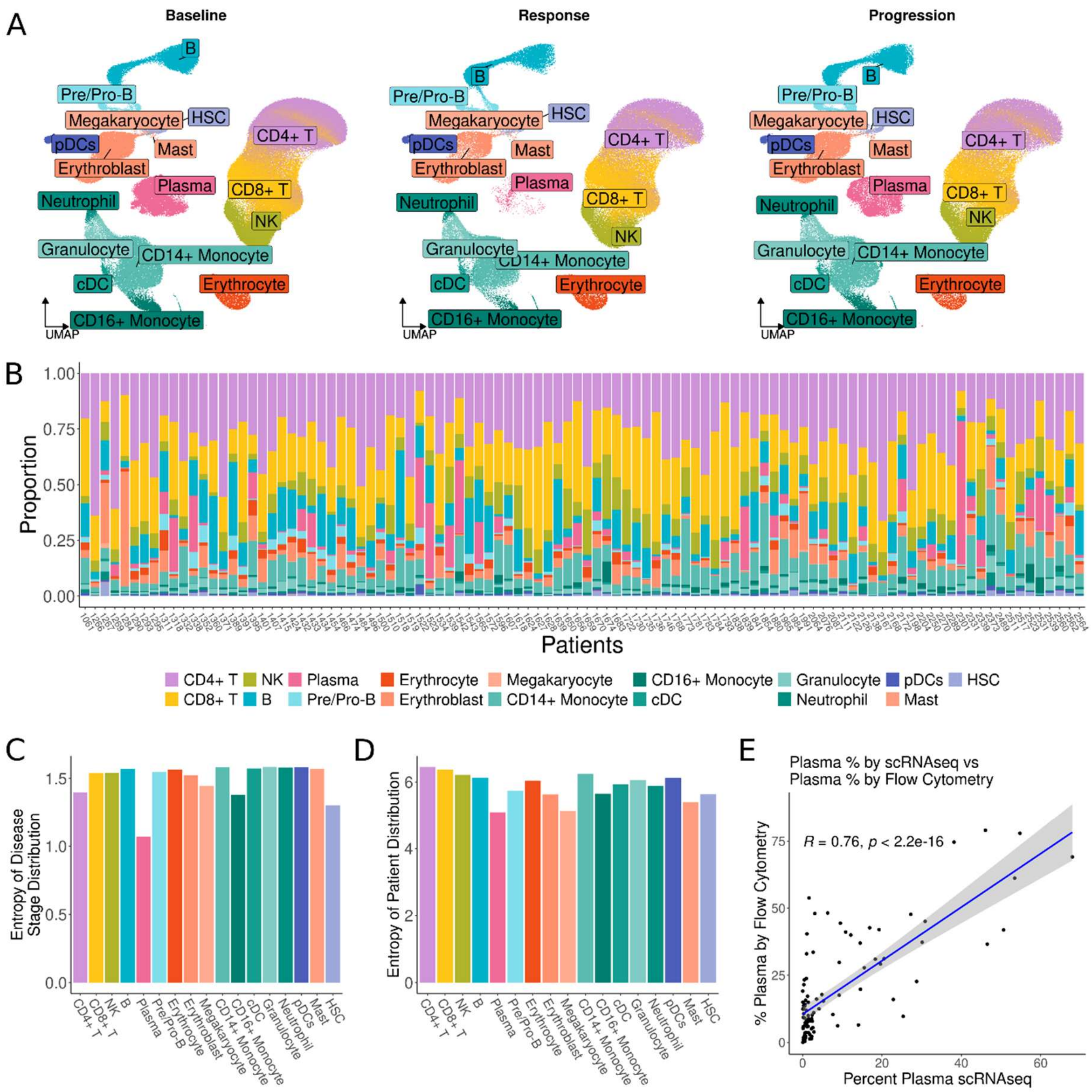


**Figure S2:** Distribution of cell types across disease stages and patients in CD138neg single cell RNA sequencing. **A.** UMAP embeddings as in Figure 1D split by disease stage with 300,232 cells from baseline samples, 156,255 cells from response samples, and 174,739 cells from progression samples. Points are colored by cell type. **B.** Bar chart of the proportion of cells annotated as a given cell type for each patient. The X axis displayed the 102 patients in the longitudinal cohort. The bars are colored by cell type as in A. **C.** Bar plot of the Shannon entropy of disease stage within each cell type. For each cell type, the Shannon entropy of the proportion of cells from each disease stage (baseline, response, progression) was calculated with high Shannon entropy indicating even distribution cell types across stages. **D.** Bar plot of the Shannon entropy of patients within each cell type. For each cell type, the Shannon entropy of the proportion of cells from each patient was calculated with high Shannon entropy indicating even distribution cell types across patients. **E.** Dot plot of the percentage of cells annotated as plasma in single cell RNA sequencing (x axis) against the percentage of plasma cells detected by flow cytometry. R and p values were calculated based on Pearson correlation.

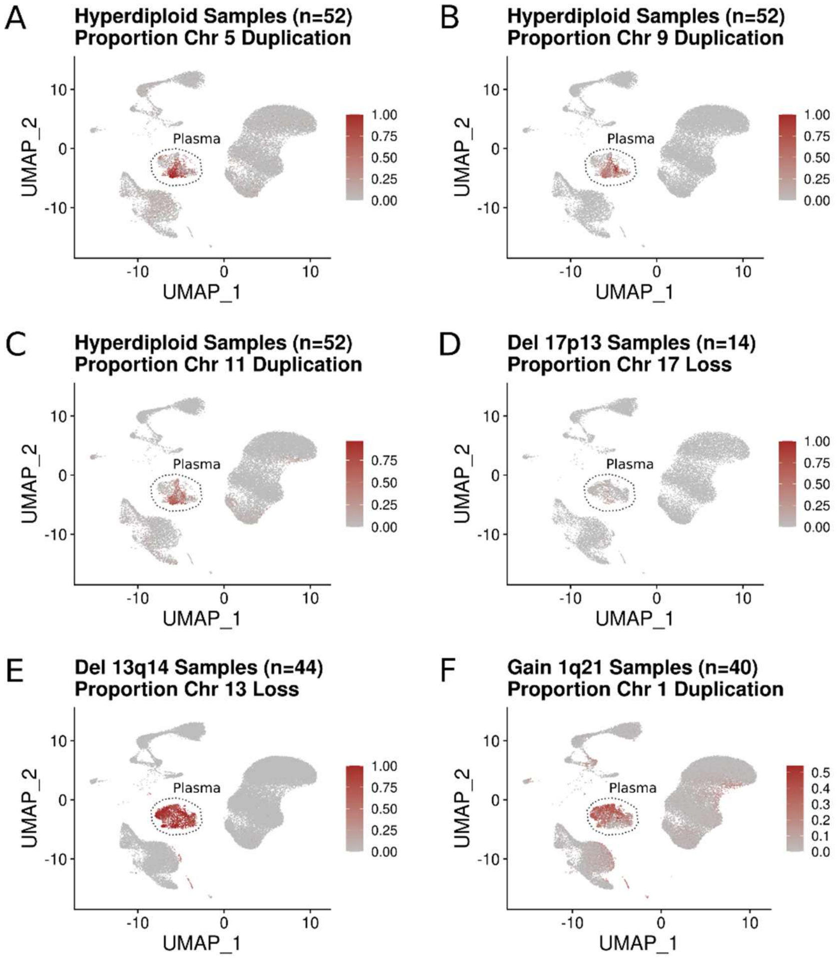


**Figure S3:** Inferred copy number indicates plasma cells are likely malignant. **A-F**. Uniform manifold approximation and projection (UMAP) embeddings for the 631,226 cells (243 samples from 102 patients) after quality filtering and doublet removal. Color scale indicates the proportion of chromosomes duplicated or lost as calculated using the inferCNV package. Copy number states were identified for each patient using the CD138$^{pos}$ whole genome sequencing data. To determine if plasma cells in scRNAseq harbored the corresponding copy number alteration (CNA), patients from a given CNA group (e.g. chromosome 1q gain) had their copy number state inferred using inferCNV on plasma cells using T, B, and myeloid cells as the reference set.

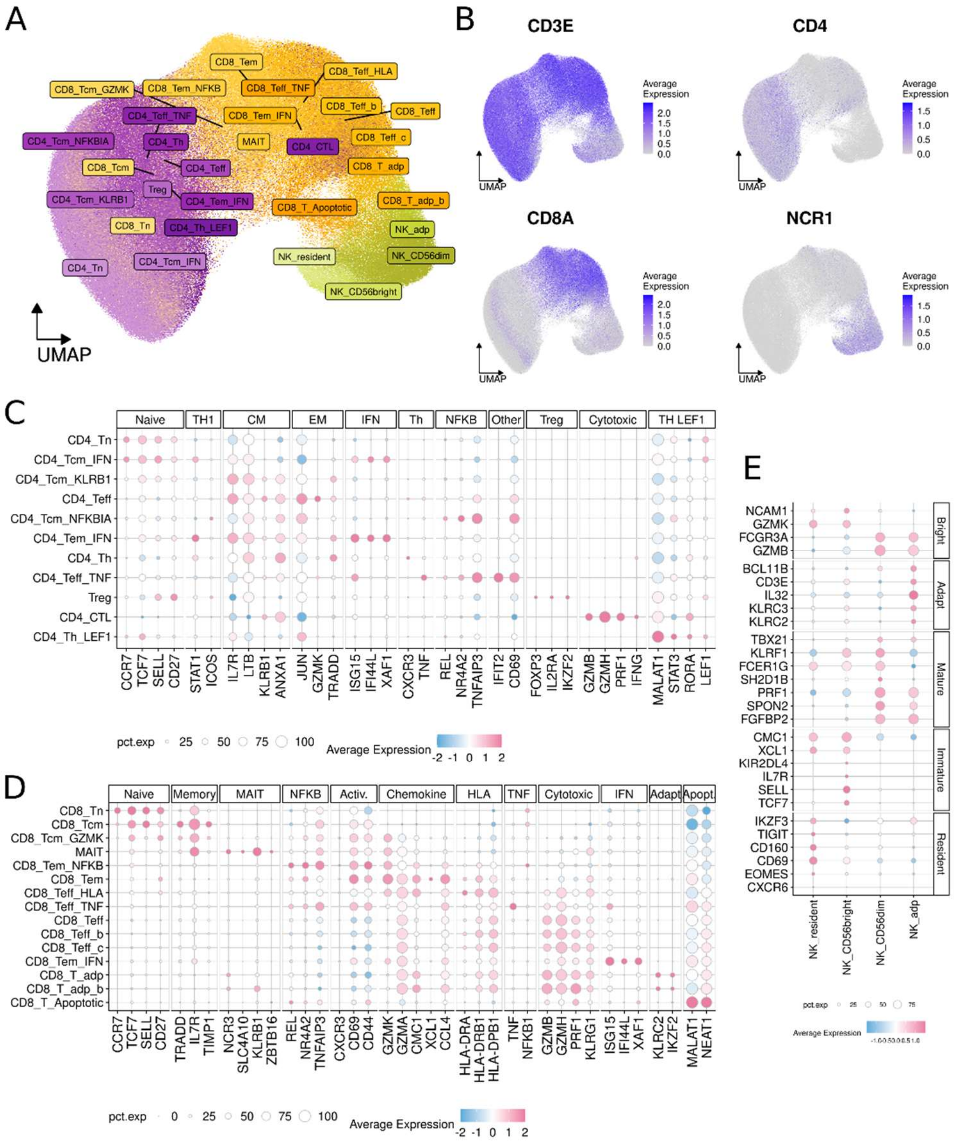


**Figure S4:** Annotation of T and NK subclusters. **A.** Uniform manifold approximation and projection (UMAP) embeddings of 30 subclusters of T and NK cells. Cells are colored by lineage (CD4⁺ T: purple, CD8⁺ T: orange, NK: yellow) and shaded by subcluster. **B.** Feature plots displayed the normalized gene expression per cell for markers distinguishing CD4⁺ T, CD8⁺ T, and NK cells. Cells are colored by gene expression with darker blue for higher expression and gray for low expression. **C-E.** Dot plots displayed the average scaled expression per cell for markers distinguishing subclusters within each cell type. Dot color indicates average normalized, scaled expression with blue for low values and pink for high values. Dot size indicates the percent of cells in the corresponding subcluster that expresses the gene.

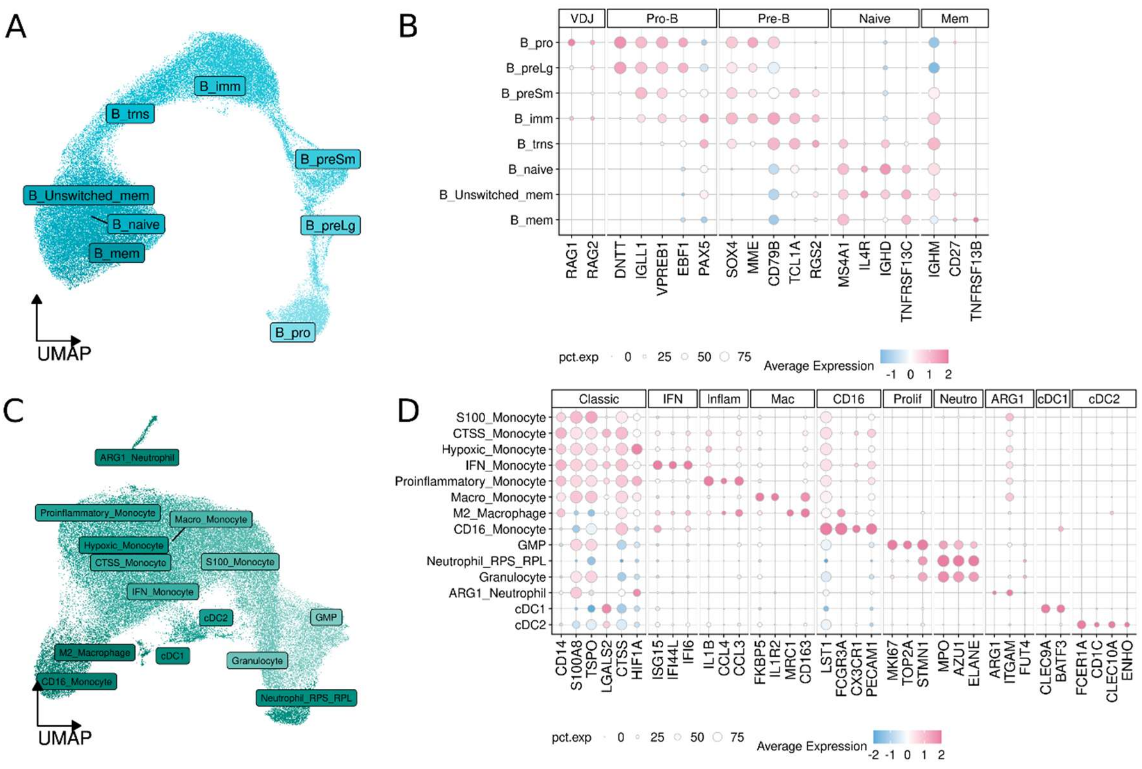


**Figure S5:** Annotation of B and myeloid subclusters. **A**. Uniform manifold approximation and projection (UMAP) of 8 subclusters of B cells. Cells are colored by lineage (B: blue) and shaded by subcluster. **B.** Dot plot displaying the average scaled expression per cell for markers distinguishing subclusters within each cell type. Dot color indicates average normalized, scaled expression with blue for low values and pink for high values. Dot size indicates the percent of cells in the corresponding subcluster that expresses the gene. **C.** UMAP of the 14 myeloid subclusters. Cells are colored by lineage (Myeloid: green) and shaded by subcluster. **D.** Dot plot displaying the average scaled expression per cell for markers distinguishing subclusters within each cell type. Dot color indicates average normalized, scaled expression with blue for low values and pink for high values. Dot size indicates the percent of cells in the corresponding subcluster that expresses the gene.

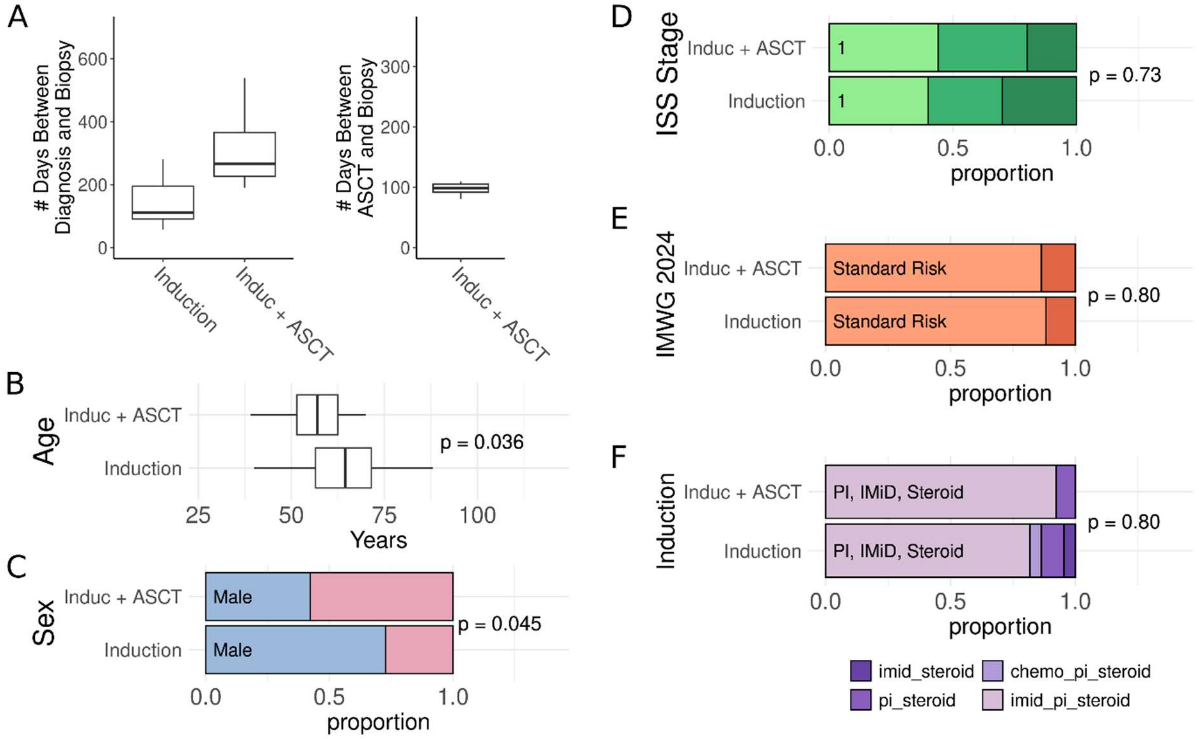


**Figure S6:** Distribution of clinical features in the baseline versus first response cohort. **A.** Box plots of the number of days from diagnosis to first response biopsy (left) and from autologous stem cell transplant to first response biopsy (right) in the longitudinal baseline to first response cohort. **B.** Box plot of the age distribution for patients with longitudinal samples from baseline and first response after induction, or baseline and first response after induction and autologous stem cell transplant (ASCT). P value was calculated using student's t test. **C-F.** Bar charts for sex, International Staging System (ISS) disease stage, International Myeloma Working Group (IMWG) risk group, and induction therapy in the induction cohort and the induction with ASCT cohort. Bar colors indicate sex (blue for male, pink for female), green for ISS stage (light-1, medium-2, dark-3), orange for IMWG risk (light-standard risk, dark-high risk), and purple for induction therapy. P values were calculated using chi squared test or Fisher's exact test if any group $n < 5$ (IMWG risk, induction therapy).

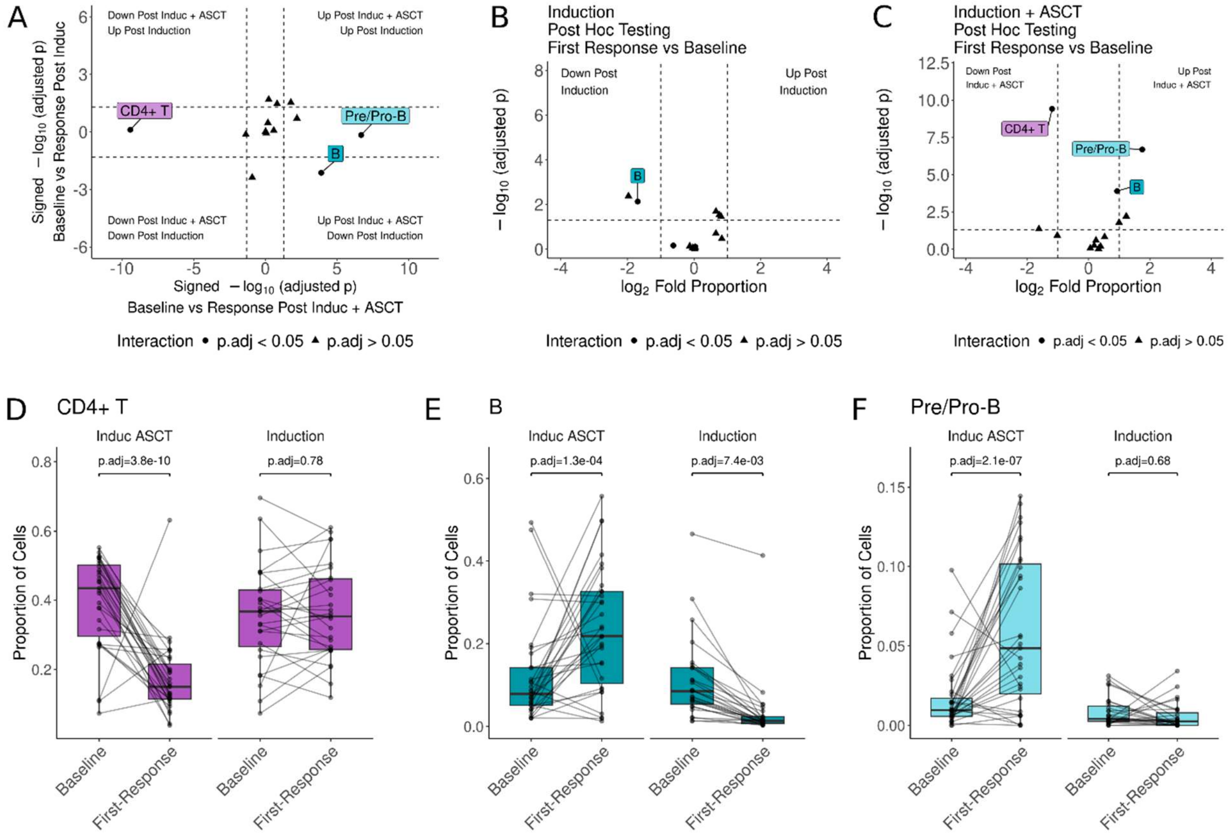


**Figure S7:** Induction with autologous stem cell transplant associates with decreased $CD4^+$ T cells and increased B cells. **A.** Dot plot displaying the increases and decreases in immune cell type proportion from baseline to first response after induction on the Y axis, and from baseline to first response post induction and autologous stem cell transplant (ASCT) on the X axis. A linear mixed effect model was used to compare subcluster proportions and changes were considered significant based on multiple comparisons adjusted p-value (adj. p) < 0.05 for interaction between time and treatment (i.e. the change from baseline to first response was different in the induction and induction with ASCT groups) and an adj. p < 0.05 by Tukey post-hoc testing (i.e. the change from baseline to first response was significant in at least one of the treatment groups). Dot shape indicates the adj. p value of the interaction term with position in the plot indicating the $log_2$ fold change signed adj. p for Tukey post hoc testing. The X-axis depicts induction and ASCT, with values on the right indicating a significant increase in proportion at first response and values on the left significant decreases. The Y axis depicts induction, with values on the top of the graph indicating significant increases at first response and values at the bottom significant decreases. **B-C.** Volcano plots of Tukey post-hoc testing of cell type proportions for baseline to first response post-induction (B) and baseline to first response post-induction and ASCT (C). The X-axis displays the $log_2$ fold of cell proportions of first-response versus baseline, and the Y-axis displays the $-log_{10}$(p.adj). The dot shape indicates the adj. p-value of the interaction term between time and treatment. **D-F.** Box and dot plots of cell proportions for $CD4^+$ T cells (D), B cells (E), and Pre/Pro B cells (F). Lines connect the dots corresponding to the same patient at each time point. P values were calculated using Tukey post-hoc testing as described in A.

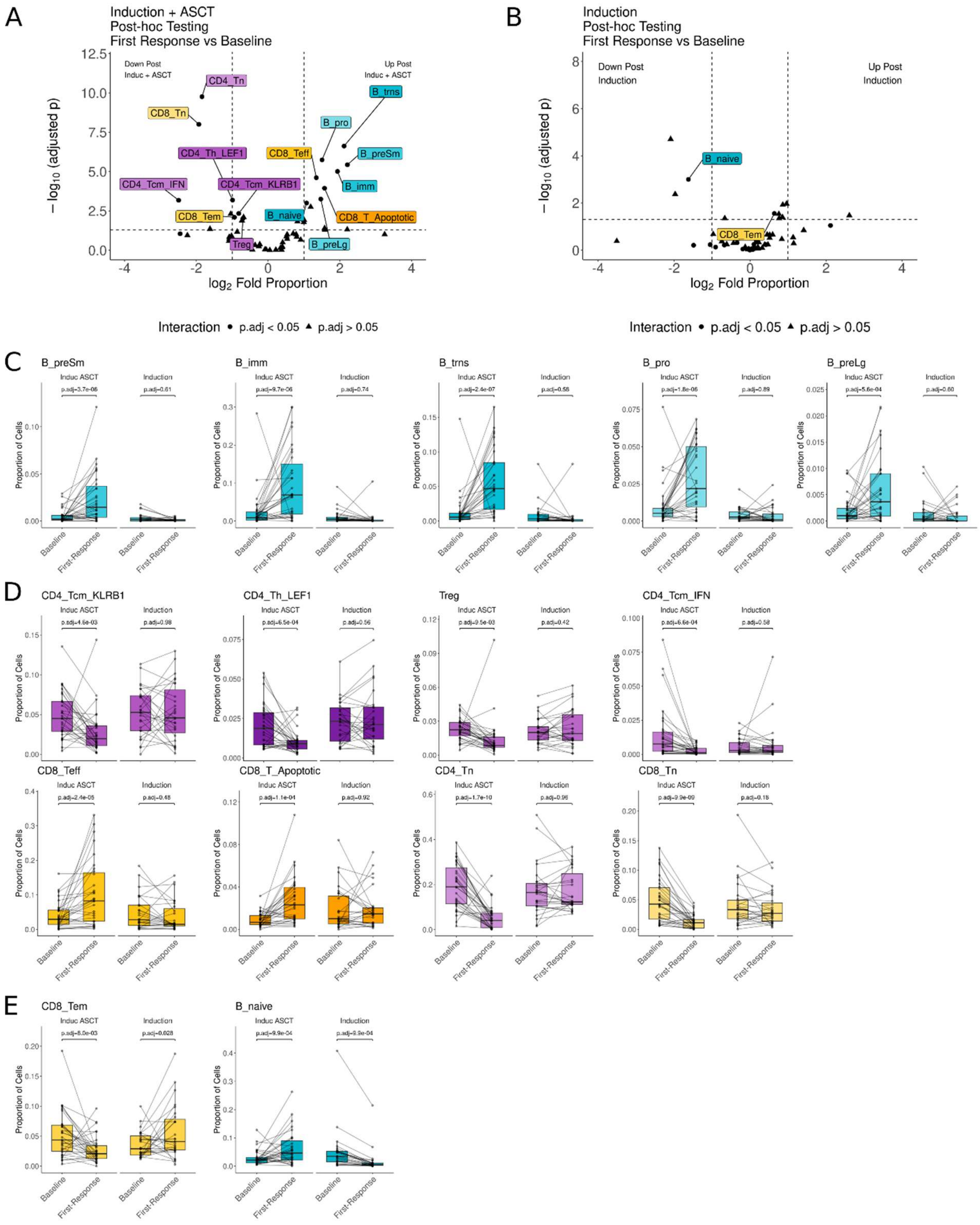


**Figure S8**: Induction with autologous stem cell transplant promotes an effector shift in CD8+ T cells, a decrease in CD4+ T cells, and increase in early B cells. **A-B.** Volcano plots of Tukey post-hoc testing of cell subcluster proportions for baseline to first response post-induction and autologous stem cell transplant (ASCT) (A) and baseline to first response post induction (B). The X axis displays the $\log_2$ fold of cell proportions of first-response versus baseline, and the Y axis displays the $-\log_{10}$(p.adj). Significance was calculated using a linear mixed effects model of cell proportion as a function of time and treatment while controlling for age and sex. Dot shape indicates the adj. p value of the interaction term between time and treatment. Post hoc testing corresponds with the results displayed in Figure 3B. **C-F.** Box and dot plots of cell proportions for B subclusters that increased only in the induction plus ASCT cohort (C), T subclusters that only changed in the induction plus ASCT cohort (D), and subclusters that changed in both cohorts in opposite direction (CD8+ effector memory T and naïve B) (E). Lines connect the dots corresponding to the same patient at each timepoint. P values were calculated using Tukey post-hoc testing as described in A.

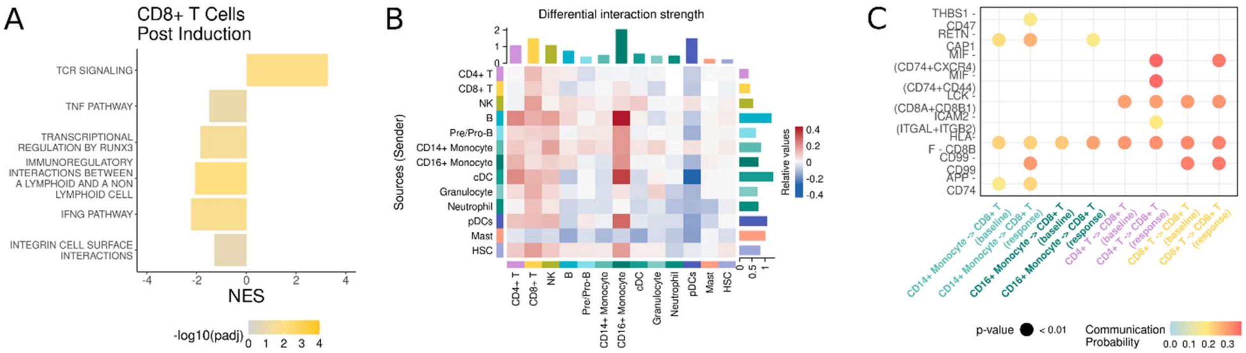


**Figure S9:** CD8+ T cells after induction do not display the same transcriptomic features as after induction and autologous stem cell transplant. These display the results for analyses comparing baseline to first response in the induction only cohort, corresponding to Figure 2D, F, and G. **A.** Bar plot of gene set enrichment analysis of CD8+ T effector cells comparing post-induction response to baseline. The X axis indicates normalized enrichment score (NES) and the bar color indicates -$\log_{10}$(adj. p) with grey for adj. p approaching 1.0 and yellow for small adj. p values. **B.** Heatmap depicting the difference in interaction strength between immune cells at first-response post-ASCT versus baseline as calculated by CellChat. Heatmap colors indicate the relative strength of interaction between two cell types at first-response post-ASCT as compared to baseline with red for increased and blue for decreased. Bars along the top of the plot indicate cumulative relative strength of incoming signals, and bars along the right side indicate cumulative relative strength of outgoing signals. **C.** Dot plot of pathways with differential communication probability between first response post-ASCT and baseline. The X axis indicates the interacting cell types and timepoint, the Y axis indicates the genes predicted to interact. Dots are colored based on communication probability with blue for low probability and red for high probability.

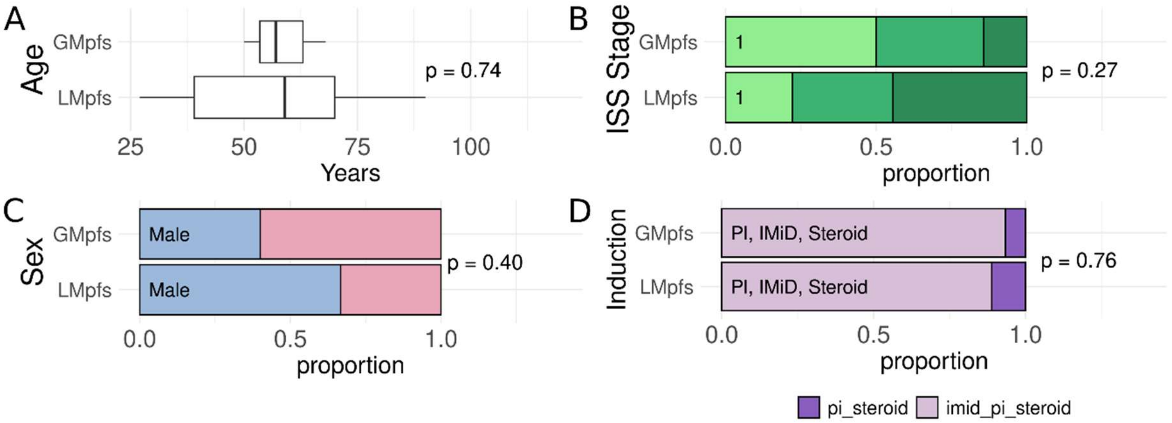


**Figure S10:** Distribution of clinical features in the baseline versus first-response post autologous stem cell transplant comparison of patients with greater than median progression free survival (GMpfs, n = 15) or less than median progression free survival (LMpfs, n = 9). **A.** Box plot for the age of patients in the GMpfs and LMpfs groups. P values was calculated using student's t test. **B-D.** Bar plots of the proportion of patients by International Staging System (ISS) disease stage (B), sex (C), and induction therapy (D). P values were calculated using Fisher's exact test.

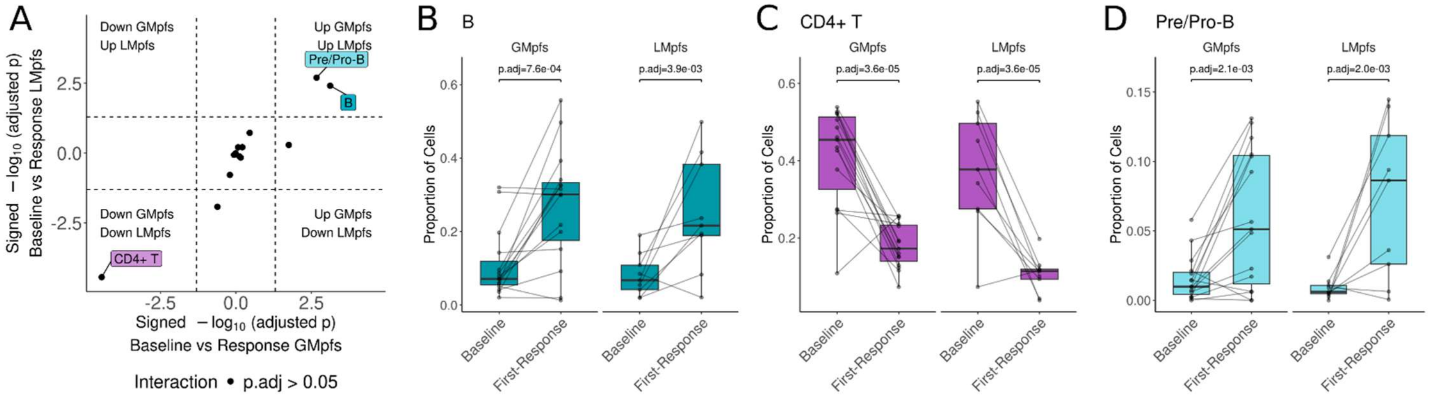


**Figure S11:** Cell proportion changes from baseline to first response post autologous stem cell transplant (ASCT) are comparable in patients with greater than median progression free survival (GMpfs, n = 15) and less than median progression free survival (LMpfs, n = 9). **A.** Dot plot displaying the increases and decreases in immune cell type proportion from baseline to first response after ASCT in the LMpfs group on the Y axis, and from baseline to first response post ASCT for the GMpfs group on the X axis. A linear mixed effect model was used to compare cell type proportions and changes were considered significant based on multiple comparisons adjusted p value (adj. p) < 0.05 for interaction between time and PFS group (i.e. the change from baseline to first response was different in the GMpfs and LMpfs groups) and an adj. p < 0.05 by Tukey post-hoc testing (i.e. the change from baseline to first response was significant in at least one of the PFS groups). Dot shape indicates the adj. p value of the interaction term with position in the plot indicating the $\log_2$ fold change signed adj. p for Tukey post hoc testing. The X axis depicts GMpfs, with values on the right indicating significant increase in proportion at first response and values on the left significant decreases. The Y axis depicts LMpfs, with values on the top of the graph indicating significant increases at first response and values at the bottom significant decreases. **B-D.** Box and dot plots of cell proportions for B cells (B) $CD4^+$ T cells (C), and Pre/Pro B cells (D). Lines connect the dots corresponding to the same patient at each timepoint. P values were calculated using Tukey post-hoc testing as described in A.

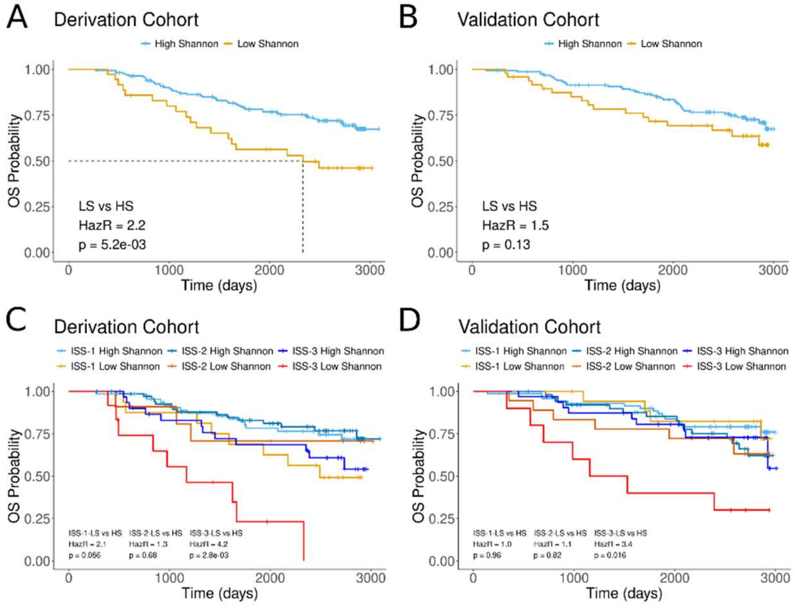


**Figure S12:** Overall survival in patients with high and low Shannon diversity index on serum immunoglobulins. **A-B.** Kaplan Meier curve for the derivation (A) and validation (B) cohorts of overall survival (OS) Lines are colored by group with yellow for low Shannon index on serum immunoglobulins and blue for high Shannon index. Patients were grouped into high and low Shannon index groups using the CutP method on progression free survival of the derivation cohort. Hazard ratio (HazR) and p value calculated by Cox proportional hazards model. **C-D.** Multivariate Kaplan Meier curves for OS of ISS stage and Shannon index group on serum immunoglobulins. Line colors indicate ISS-Shannon group with shades of blue for high Shannon and shades of yellow-orange for low Shannon index. Hazard ratios and p values calculated by Cox proportional hazards model.

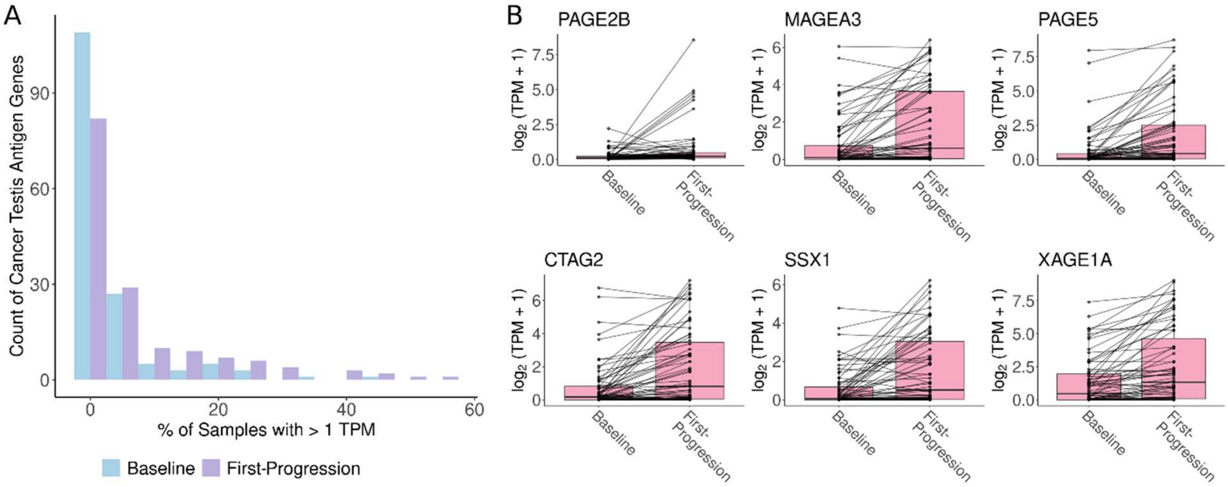


**Figure S13:** Expression of cancer testis antigens (CTAg) is specific to a subset of samples in the longitudinal cohort. **A.** Density plots of the relative quantity (Y axis) of CTAg genes by the percentage of samples expressing CTAg at least 1 transcript per million (TPM, X axis) in baseline and first progression samples. 154 CTAg genes were selected based on being a part of CTAg families identified in at least 2 CTAg databases, multiple genes within the CTAg family, located on the X chromosome, and expressed in < 90% of samples. **B.** Box plots of CTAg gene expression in CTAg-enriched samples versus all other samples. The y axis displays $\log_2$(TPM + 1).

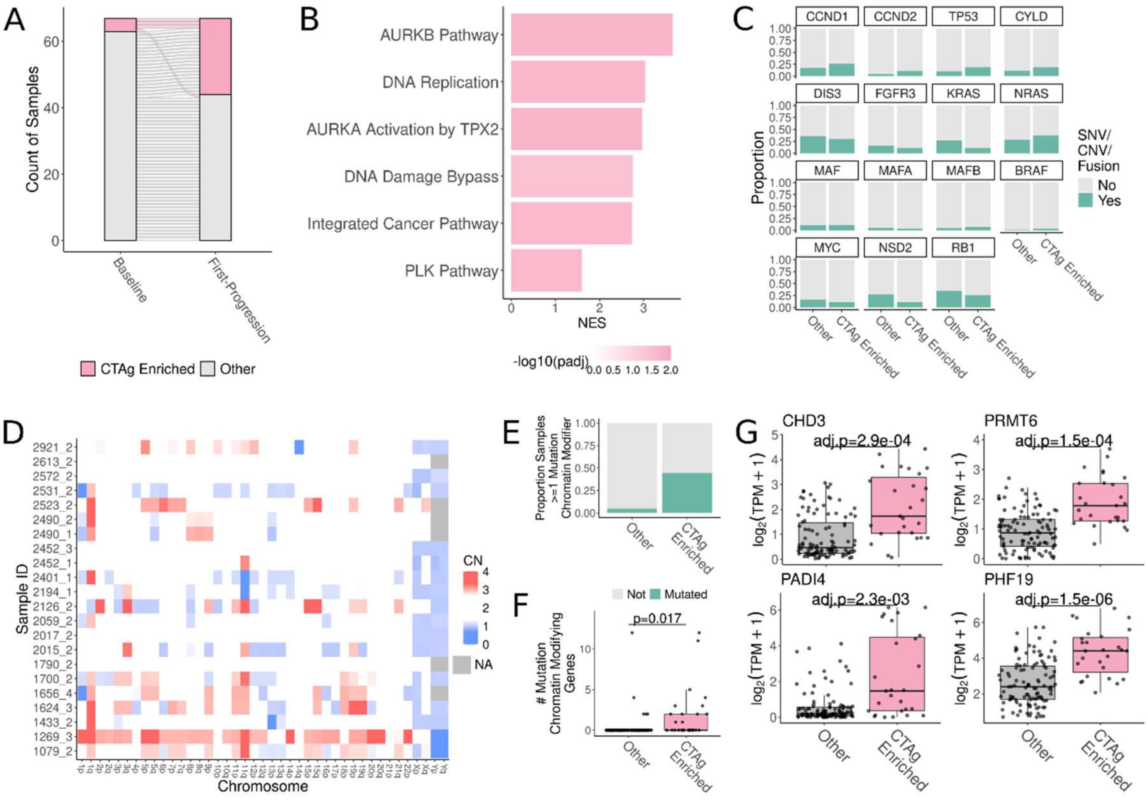


**Figure S14:** Samples enriched for cancer testis antigens (CTAg) are enriched for mutations and expression in genes for chromatin modifiers. **A.** Alluvial plot displaying samples with high expression of progression DEGs. While differential expression was executed on all 67 longitudinal sample pairs, less than half of the samples expressed many of the top DEGs (**Supplemental Fig 13**). To identify samples enriched for the progression-associated DEGs we calculated the average $\log_2$(TPM + 1) of positive DEGs and considered samples 1 standard deviation above the mean as enriched (labeled CTAg-enriched). **B.** Bar plot of gene set enrichment analysis of CD138$^{pos}$ bulk RNA sequencing comparing first progression to baseline. The X axis indicates normalized enrichment score (NES) and the bar color indicates $-\log_{10}$(adj. p) with grey for adj. p approaching 1.0 and pink for small adj. p values. **C.** Bar plots displaying common mutations seen in multiple myeloma in CTAg-enriched samples as compared to all others in the first progression longitudinal cohort. **D.** Copy number heatmap for the CTAg-enriched samples with red for copy number gain and blue for copy number loss. **E.** Bar plot of the proportion of CTAg-enriched and other samples with mutations in a selected list of 36 chromatin modifying genes (see **Supplemental Fig 16** for genes and types of mutations). A list of 389 genes from the Gene Ontology database was reduced by recursive feature elimination to find the minimum list of genes that maximizes the difference in proportion of CTAg-enriched samples and other samples having a mutation in the corresponding gene list. **F.** Box plot of the number of mutations per patient from the 36-gene list chromatin modifying gene mutations in CTAg-enriched samples and other samples. **G.** Box plots of chromatin modifying genes differentially expressed in CTAg-enriched samples compared to other samples. Adjusted p value as calculated in B.

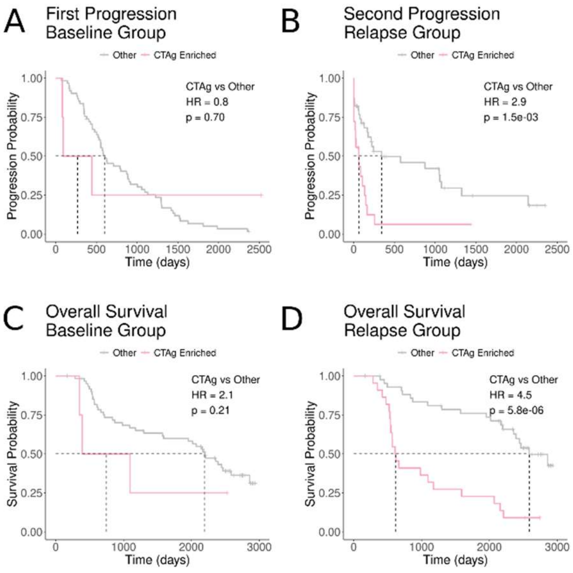


**Figure S15:** Patients enriched for cancer testis antigen (CTAg) have worse outcomes. **A-D.** Kaplan meier curves for progression free survival based on baseline CTAg group (A), second progression based on CTAg group at first progression (B), and overall survival based on baseline group (C) and first progression group (D). CTAg enrichment was calculated based average $\log_2$(transcript per million + 1) for differentially expressed genes ($\log_2$ fold change > 0.5, adjusted $p < 0.05$) at first progression versus baseline, four patients were enriched at baseline and 23 were enriched at first progression (see **Fig 3C**). Hazard ratios (HR) and p values were calculated by Cox proportional hazards model.

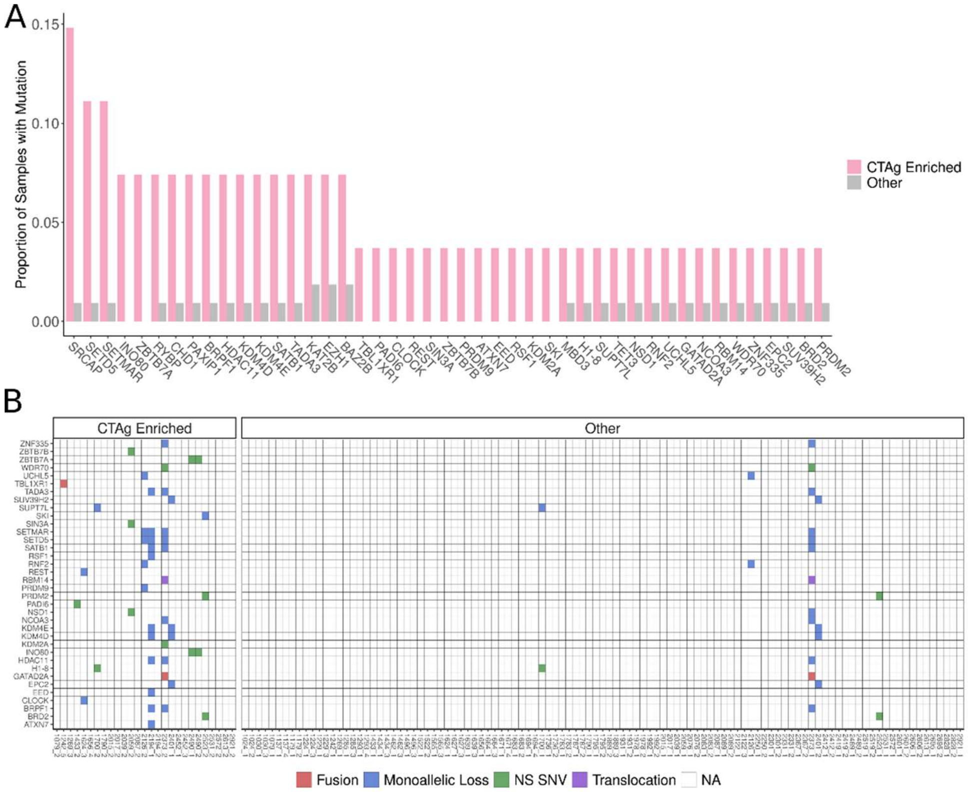


**Figure S16:** Mutations in chromatin modifying genes specific to cancer testis antigen (CTAg) enriched multiple myeloma (MM). **A.** Bar plot of 45 chromatin modifying genes present in greater proportion of CTAg-enriched samples as compared to other samples. **B.** Tile plot of 36 chromatin modifying genes specific to CTAg-enriched MM after recursive feature elimination from A. Tile color indicates mutation type with blue for copy number loss, green for non-synonymous single nucleotide variant, purple for translocation, and red for gene fusion.

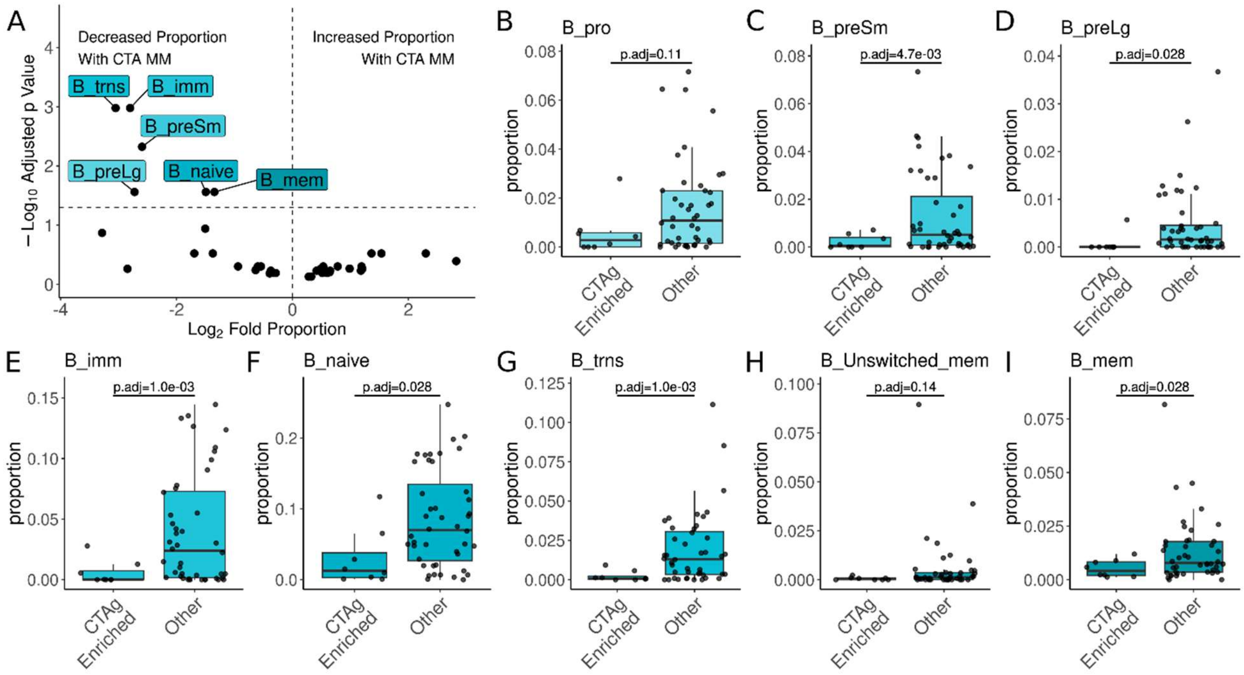


**Figure S17:** Differentially abundant cell subclusters in first progression samples with and without cancer testis antigen (CTAg) enriched myeloma. **A.** Volcano plots of the differential abundance of cell subclusters in CTAg-enriched MM versus all other samples. The X axis displays the $\log_2$ fold of cell subcluster proportions between CTAg-enriched and other samples. The Y axis displays the $-\log_{10}$(adjusted p value) based on a linear model of subcluster proportion and CTAg group controlling for age and International Staging System disease stage. **B-I.** Box and dot plots of cell proportions for B cell subclusters. Significance was tested by a linear model of cell type proportion and CTAg group while accounting for age and international staging system disease stage with Benjamini-Hochberg multiple comparisons adjustment.

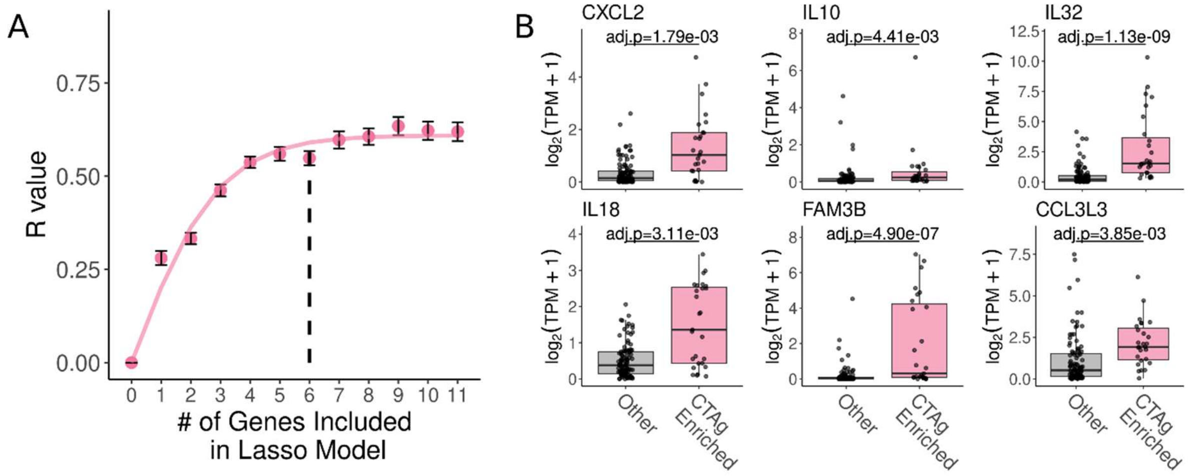


**Figure S18:** Lasso regression identifies cytokine/chemokine genes in CD138pos bulk RNA sequencing associated with B cell abundance. **A.** Dot plot of the cumulative correlation of Lasso models with increasing number of genes included from B. The x axis displays the number of genes included in the Lasso model and the points indicate mean r value with standard error bars for the one hundred boostraps. Starting with the highest ranked gene, one hundered bootstrapped Lasso models of MM gene expression against B cell proportion. Then, the next highest ranked gene was iteratively added to the model and assessed for an increase in r value. A saturating exponential model was fit to the resulting r values to estimate the number of genes at which the r value plateaus (i.e. the minimum number of genes to explain most of the association between MM gene expression and naïve B proportion). **B.** Box plots of cytokine/chemokine genes identified in the top 6 association with B cell proportion from Lasso regression. Statistical significance tested using limma voom with Benhamini Hochberg multiple comparisons correction.

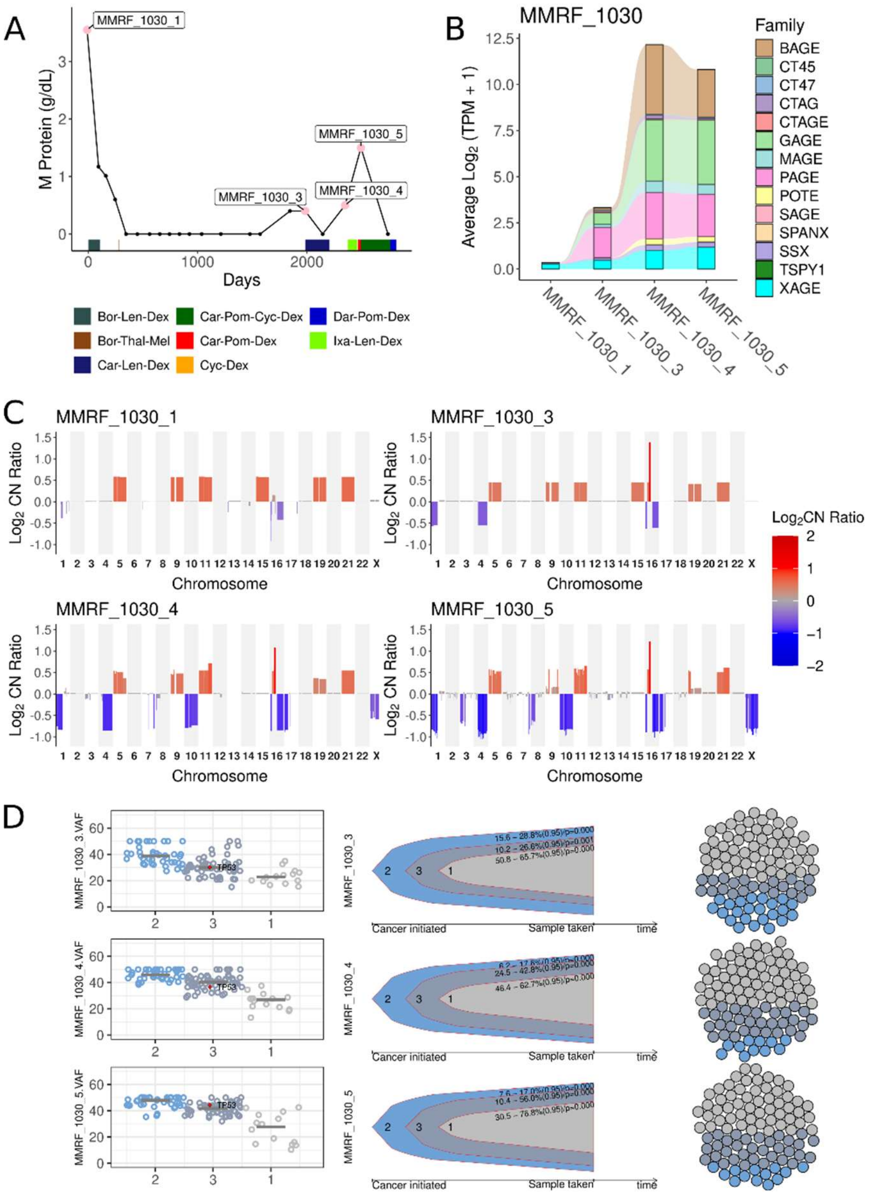


**Figure S19:** Longitudinal evaluation of CD138$^{pos}$ DNA and RNA from MMRF_1030. **A.** Line plot of serum M protein measurements over time. Pink dots and labels annotate the visits at which biopsies were taken for RNA and DNA sequencing. Bars below the line indicate lines of treatment. **B.** Alluvial plot of cancer testis antigen (CTAg) gene expression across longitudinal samples. Alluvium height indicates the average $Log_2$ transcripts per million (TPM) for genes in each CTAg family. **C.** Copy number plot split by longitudinal samples. The X axis displays chromosome number, the Y axis and bar color displays the $Log_2$ copy number ratio with red for copy number gain and blue for copy number loss. **D.** Subclonal reconstruction of longitudinal samples. Subclonal structure was inferred based on the cancer cell fraction of single nucleotide variants (SNV) using QuantumClone. SNVs in genes coding for chromatin modifying proteins are labeled and highlighted in red (left). Bell plots display inferred subclonal evolution from the SNV cancer cell fractions (middle). Dot chart depicts the relative proportion of each subclone across time points (right).

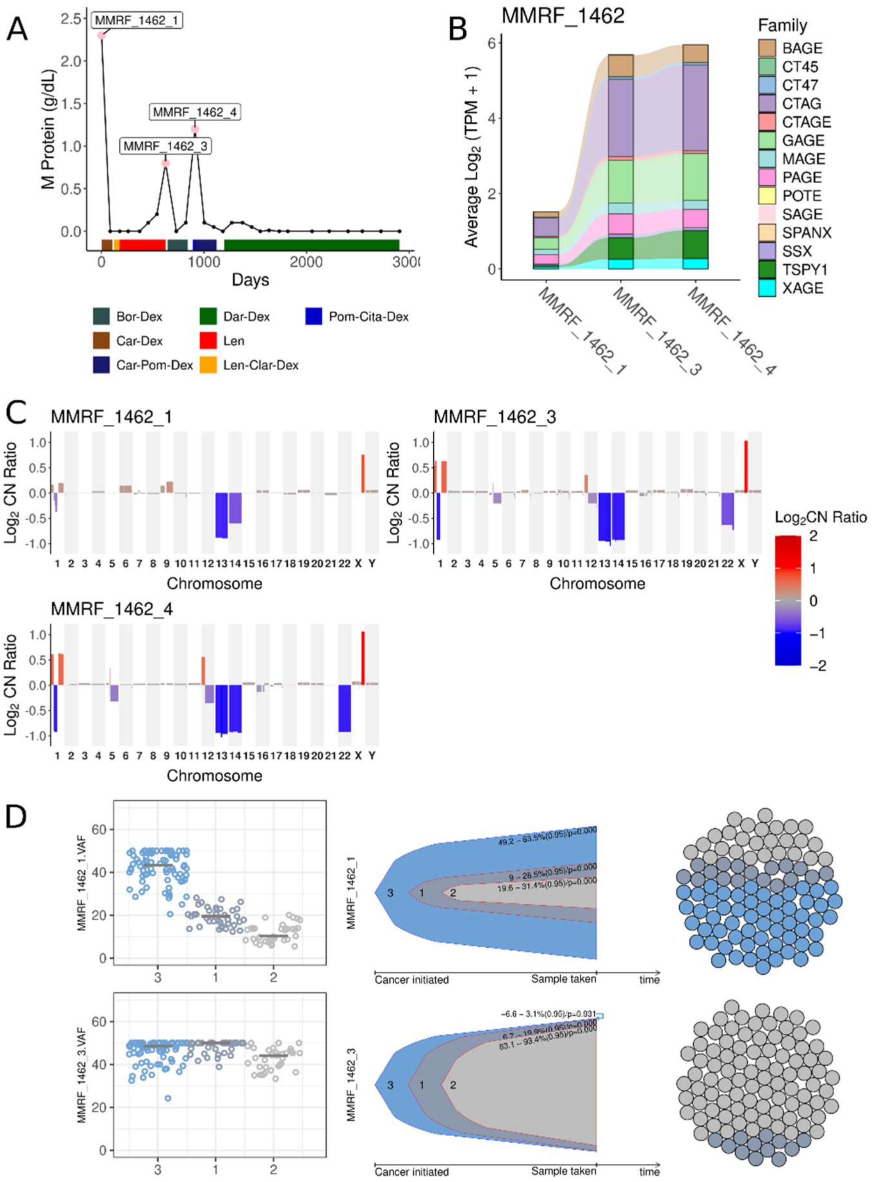


**Figure S20:** Longitudinal evaluation of CD138$^{pos}$ DNA and RNA from MMRF_1462. **A.** Line plot of serum M protein measurements over time. Pink dots and labels annotate the visits at which biopsies were taken for RNA and DNA sequencing. Bars below the line indicate lines of treatment. **B.** Alluvial plot of cancer testis antigen (CTAg) gene expression across longitudinal samples. Alluvium height indicates the average $Log_2$ transcripts per million (TPM) for genes in each CTAg family. **C.** Copy number plot split by longitudinal samples. The X axis displays chromosome number, the Y axis and bar color displays the $Log_2$ copy number ratio with red for copy number gain and blue for copy number loss. **D.** Subclonal reconstruction of longitudinal samples. Subclonal structure was inferred based on the cancer cell fraction of single nucleotide variants (SNV) using QuantumClone. SNVs in genes coding for chromatin modifying proteins are labeled and highlighted in red (left). Bell plots display inferred subclonal evolution from the SNV cancer cell fractions (middle). Dot chart depicts the relative proportion of each subclone across time points (right).

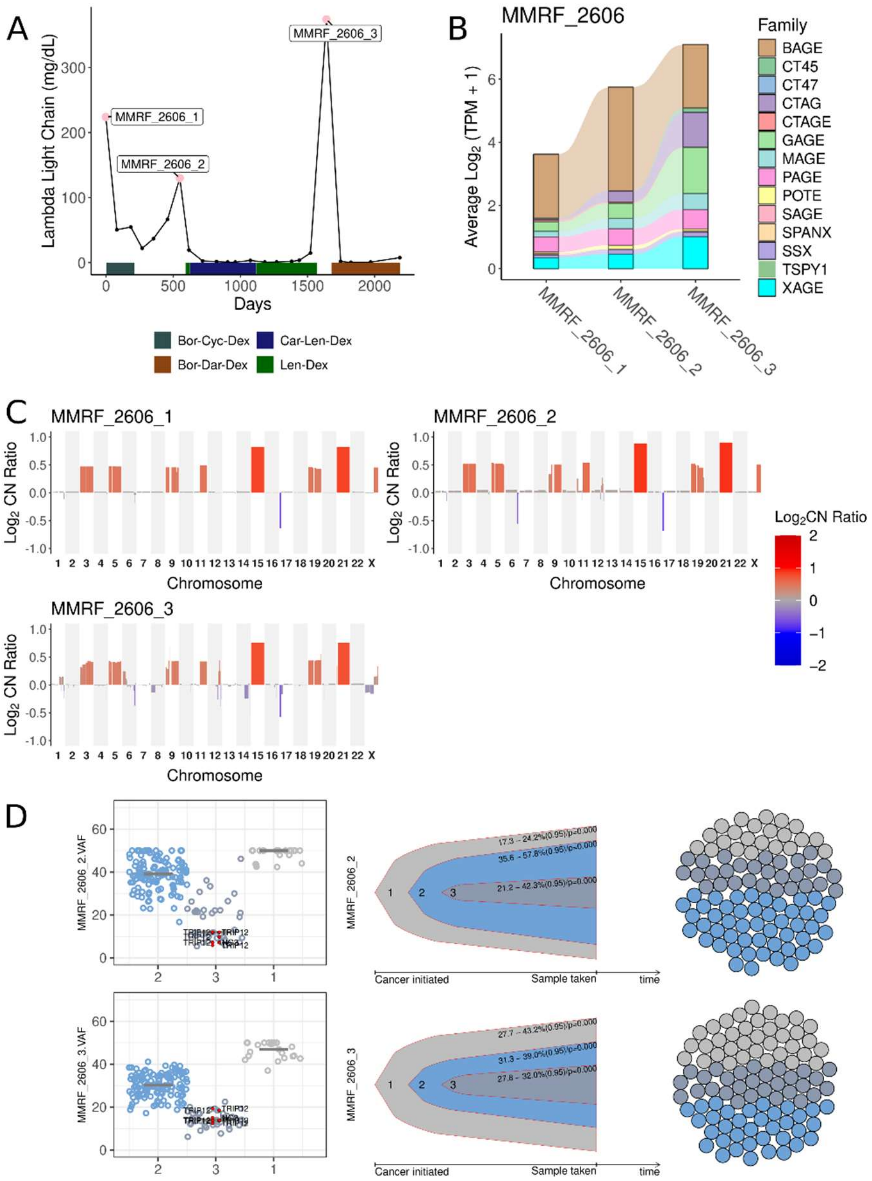


**Figure S21:** Longitudinal evaluation of CD138$^{pos}$ DNA and RNA from MMRF_2606. **A.** Line plot of serum lambda light chain measurements over time. Pink dots and labels annotate the visits at which biopsies were taken for RNA and DNA sequencing. Bars below the line indicate lines of treatment. **B.** Alluvial plot of cancer testis antigen (CTAg) gene expression across longitudinal samples. Alluvium height indicates the average $Log_2$ transcripts per million (TPM) for genes in each CTAg family. **C.** Copy number plot split by longitudinal samples. The X axis displays chromosome number, the Y axis and bar color displays the $Log_2$ copy number ratio with red for copy number gain and blue for copy number loss. **D.** Subclonal reconstruction of longitudinal samples. Subclonal structure was inferred based on the cancer cell fraction of single nucleotide variants (SNV) using QuantumClone. SNVs in genes coding for chromatin modifying proteins are labeled and highlighted in red (left). Bell plots display inferred subclonal evolution from the SNV cancer cell fractions (middle). Dot chart depicts the relative proportion of each subclone across time points (right).

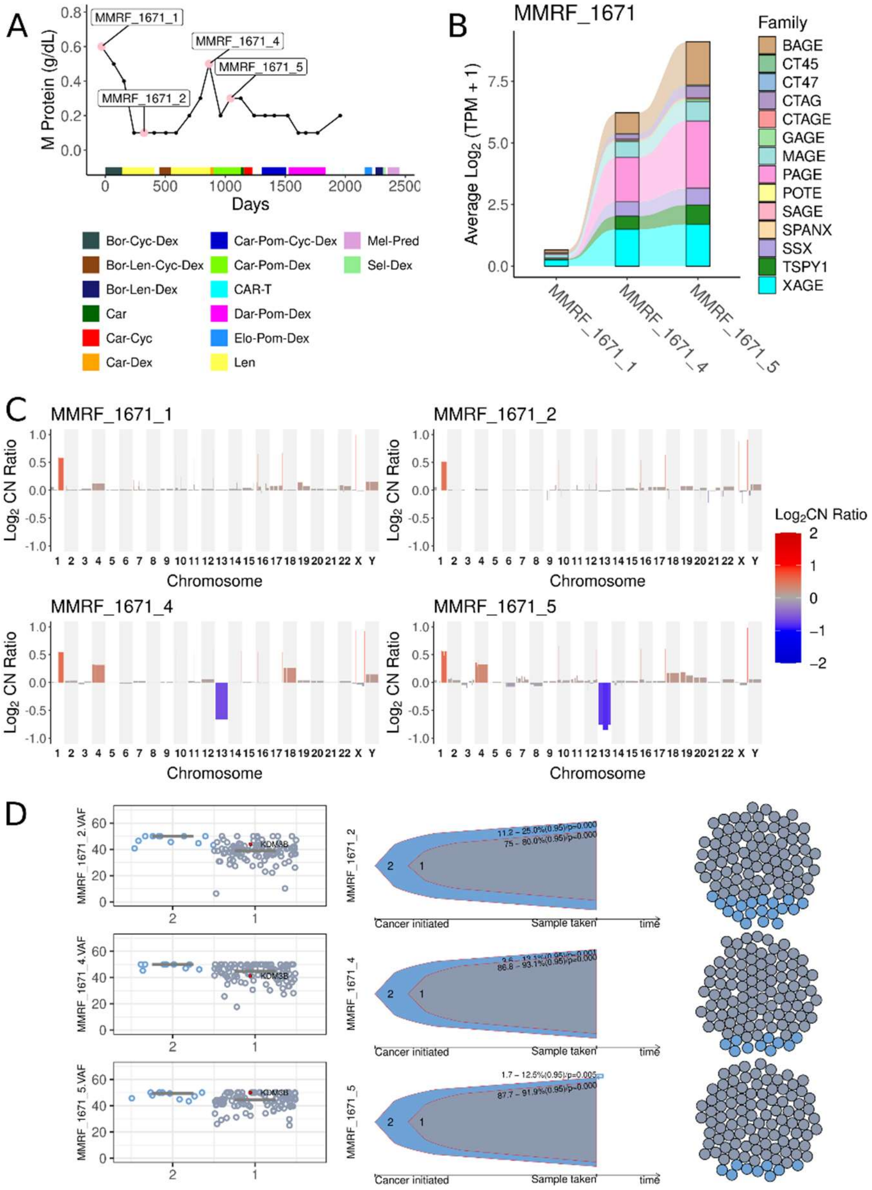


**Figure S22:** Longitudinal evaluation of CD138$^{pos}$ DNA and RNA from MMRF_1671. **A.** Line plot of serum M protein measurements over time. Pink dots and labels annotate the visits at which biopsies were taken for RNA and DNA sequencing. Bars below the line indicate lines of treatment. **B.** Alluvial plot of cancer testis antigen (CTAg) gene expression across longitudinal samples. Alluvium height indicates the average Log$_2$ transcripts per million (TPM) for genes in each CTAg family. **C.** Copy number plot split by longitudinal samples. The X axis displays chromosome number, the Y axis and bar color displays the Log$_2$ copy number ratio with red for copy number gain and blue for copy number loss. **D.** Subclonal reconstruction of longitudinal samples. Subclonal structure was inferred based on the cancer cell fraction of single nucleotide variants (SNV) using QuantumClone. SNVs in genes coding for chromatin modifying proteins are labeled and highlighted in red (left). Bell plots display inferred subclonal evolution from the SNV cancer cell fractions (middle). Dot chart depicts the relative proportion of each subclone across time points (right).

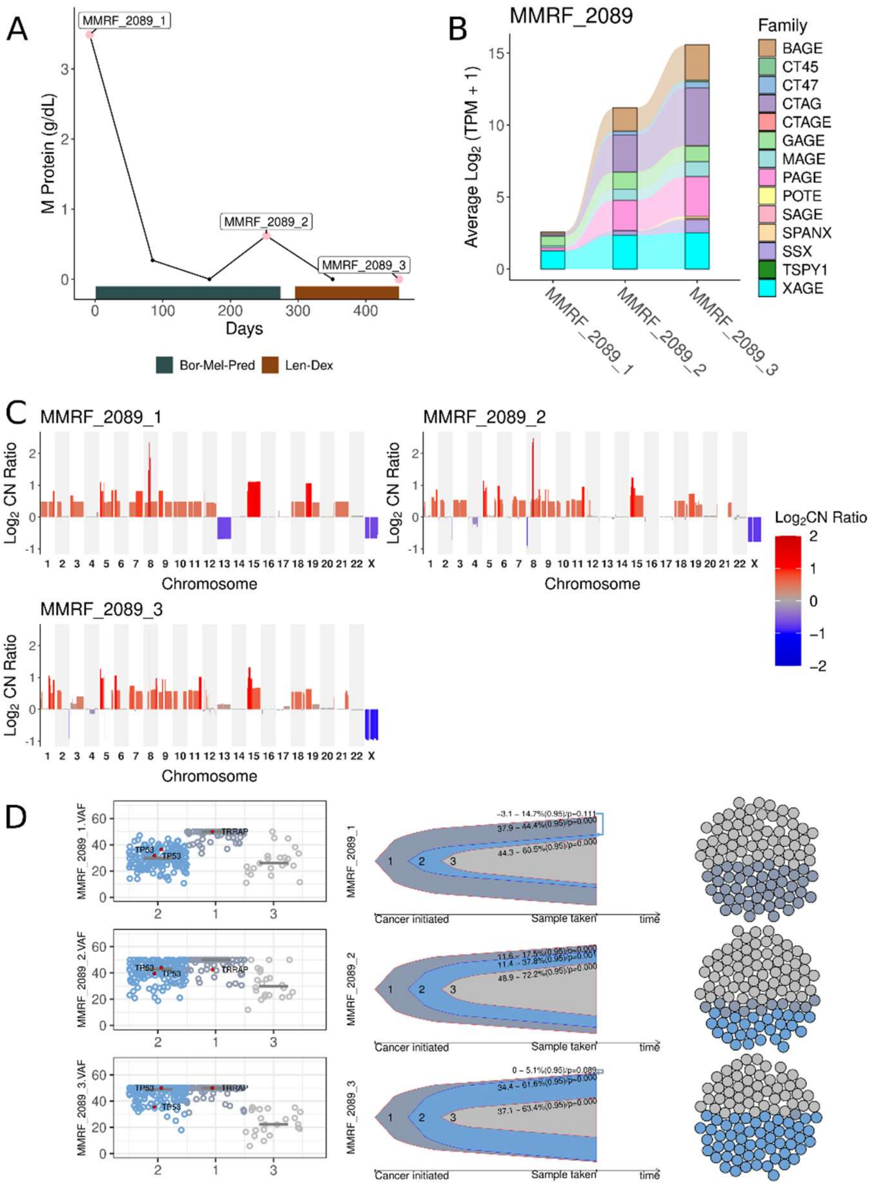


**Figure S23:** Longitudinal evaluation of CD138$^{pos}$ DNA and RNA from MMRF_2089. **A.** Line plot of M protein measurements over time. Pink dots and labels annotate the visits at which biopsies were taken for RNA and DNA sequencing. Bars below the line indicate lines of treatment. **B.** Alluvial plot of cancer testis antigen (CTAg) gene expression across longitudinal samples. Alluvium height indicates the average Log$_2$ transcripts per million (TPM) for genes in each CTAg family. **C.** Copy number plot split by longitudinal samples. The X axis displays chromosome number, the Y axis and bar color displays the Log$_2$ copy number ratio with red for copy number gain and blue for copy number loss. **D.** Subclonal reconstruction of longitudinal samples. Subclonal structure was inferred based on the cancer cell fraction of single nucleotide variants (SNV) using QuantumClone. SNVs in genes coding for chromatin modifying proteins are labeled and highlighted in red (left). Bell plots display inferred subclonal evolution from the SNV cancer cell fractions (middle). Dot chart depicts the relative proportion of each subclone across time points (right).
